# A late Neanderthal reveals genetic isolation in their populations before extinction

**DOI:** 10.1101/2023.04.10.536015

**Authors:** Ludovic Slimak, Tharsika Vimala, Andaine Seguin-Orlando, Laure Metz, Clément Zanolli, Renaud Joannes-Boyau, Marine Frouin, Lee J. Arnold, Martina Demuro, Thibaut Devièse, Daniel Comeskey, Michael Buckley, Hubert Camus, Xavier Muth, Jason E. Lewis, Hervé Bocherens, Pascale Yvorra, Christophe Tenailleau, Benjamin Duployer, Hélène Coqueugniot, Olivier Dutour, Thomas Higham, Martin Sikora

**Author notes:** These authors contributed equally to this work (Ludovic Slimak, Tharsika Vimala, Andaine Seguin-Orlando).

## Abstract

Neanderthal genomes have been recovered from sites across Eurasia, painting an increasingly complex picture of their populations’ structure, mostly indicating that late European Neanderthals belonged to a single metapopulation with no significant evidence of deep population structure. Here we report the discovery of a late Neanderthal individual, nicknamed “Thorin”, from Grotte Mandrin in Mediterranean France, and his genome. These dentognathic fossils, including a rare example of distomolars, are associated with a rich archeological record of their final technological traditions in this region ∼50-42 thousand years ago. Thorin’s genome reveals a deep divergence with other late Neanderthals. Thorin belonged to a population with small group size that showed no genetic introgression with other known late European Neanderthals, revealing genetic isolation of his lineage despite them living in neighboring regions. These results have important implications for resolving competing hypotheses about causes of the Neanderthals’ disappearance.

**One Sentence Summary:** A new French Neanderthal fossil and its genome reveal complex population dynamics during the past 100,000 years.

## Main Text

### Introduction

The reasons behind the extinction of the Neanderthals ∼40 thousand years ago (ka) are still widely debated. Multiple theories have been presented over the years, including competition or interbreeding with modern humans, but it remains unknown if the factors involved in this process were primarily ecological or social and then based on the historical inter-relations between these populations^1–3^. Some researchers suggest that social, technical, or ethological differences between Neanderthals and modern humans may have played a direct role in their demise but the precise cause(s) of such extinction remains uncertain^1–7^. Paleogenomic and osteological studies have revealed low effective population sizes and signatures of inbreeding in Siberian and late European Neanderthals^6, 8^, suggesting social structure characterized by small group sizes and low intergroup mobility. This contrasts with recent results from early Eurasian modern humans, which showed low levels of inbreeding and higher intergroup mobility despite small group sizes^9,10^. Whether these results are representative of wider Neanderthal social organization remains inconclusive.

Since the publication of the first draft of the Neanderthal genome in 2010^11^, Neanderthal genomes have been recovered from sites across Eurasia, painting an increasingly complex picture of Neanderthal genetic structure. The deepest divergence among Neanderthal genomes sequenced to date is found between eastern and western Eurasian Neanderthal populations represented by the ∼120 ka Altai Neanderthal from Denisova Cave^7^ and the >44 ka Vindija 33.19 individual from Croatia^12^. Genomic data of all other available Neanderthal remains, the earliest in western Europe being ∼120 ka (Scladina and Hohlenstein-Stadel (HST)), while the latest being ∼40 ka, suggest genetic continuity in western Eurasia for ∼80 ka^13^. Recent results obtained from sedimentary DNA suggest that the genetic landscape was significantly altered by expansions of Neanderthal populations ∼105 ka^14^. This gave rise to lineages in western Europe represented by samples from Central Europe (Vindija), the Caucasus (Mezmaiskaya Cave), and Siberia (Chagyrskaya cave 8)^15^, the latter likely replacing the earlier Altai-like population. The genomes of late (<50 ka) European Neanderthals, including an individual from the Caucasus (Mezmaiskaya 2), were all found to be more similar to Vindija than to other known lineages, indicating further population turnover towards the last stages of Neanderthal history in the Caucasus or western Europe^16^. The close correlation between genetic similarities and geographic location suggested an absence of major population structure among the sampled late Neanderthal populations. It remains unknown whether these patterns result from long-term *in situ* evolution of late European Neanderthal populations, or as a consequence of a recent expansion of Vindija-like lineages into Europe.

Here we report the discovery of a late Neanderthal individual, nicknamed “Thorin”, in 2015 and progressively excavated since then at Grotte Mandrin in Mediterranean France, a site which also was temporarily occupied by early modern humans at 54 ka^1^. Thorin is one of the best represented Neanderthal individuals found in France since the discovery from Saint-Césaire in 1979^17^. Combining archaeological, chronostratigraphic, isotopic, and genomic analyses, we show that Thorin belonged to a late Neanderthal population which had stayed genetically isolated for some 50 ka. We further find evidence of gene flow from a deeply divergent lineage distinct from the Thorin lineage in the Neanderthal individual from Les Cottés^16^. Our results suggest the presence of multiple isolated late Neanderthal communities in Europe close to their time of extinction, and shed light on their social organization with limited, if any, level of interactions in between different Neanderthal populations in their last millennia.

## Results

### Thorin is a late European Neandertal

Grotte Mandrin is a rockshelter located in Mediterranean France directly overhanging the Rhône River Valley. The site records 12 main sedimentary layers dating from Marine Isotope Stages (MIS) 5 to 3. Geological and micromorphological analyses show that all archeological levels were well preserved by rapid wind deposition of sands and silts^1^. The upper sequence is divided into 8 archeological levels chronologically placed between 65.6 to 31.0 ka at 95% CI, encompassing the last Neanderthal societies and the arrival of the first modern human groups. Each of these levels provide rich archeological records, totalling more than 60,000 lithics and 70,000 faunal remains. Fireplaces and hominin remains were also found in most of Mandrin’s levels^1^. These 8 archeological levels were divided in 5 cultural phases: Level F: Rhodanian Quina, Level E: Neronian, Level D: Post-Neronian I (PNI), Levels C2 to B2 Post-Neronian II (PNII), Level B1: Protoaurignacian. The cultural determinations of the Neronian, PNI, and PNII phases at Mandrin^1^ show major technical and cultural divergences with the coeval Mousterian and Châtelperronian societies^4^ found in the neighboring regions of south-western France and _Burgundy_^3,5,18,19^.

Thorin was discovered in 2015 at the entrance of the rockshelter in lateral contact between the upper layers and the bedrock in Level B2 (Fig. 1) associated with abundant fauna and artifacts attributed to the PNII, the last Mousterian phase from Grotte Mandrin^1, 3, 5, 18, 19^. Thorin is represented by several fragments, including a portion of the left palatal process at the level of the molars, a fragmentary mandible, as well as 31 permanent maxillary and mandibular teeth (Fig. 2). While the upper right premolars and the upper left canine were lost post-mortem, it is noteworthy that two supernumerary lower molars are present (fourth molars). They are heteromorphic and exhibit a reduced and simplified (non-conical) crown with a single but large root from the cervix to the apex. The marked inclined wear facet affecting the occluso-mesial crown aspect of these two teeth fits with the distal interproximal facet of the lower third molars, indicating that the distomolars impacted the third molar crowns during the eruption process.

**Fig. 1.**
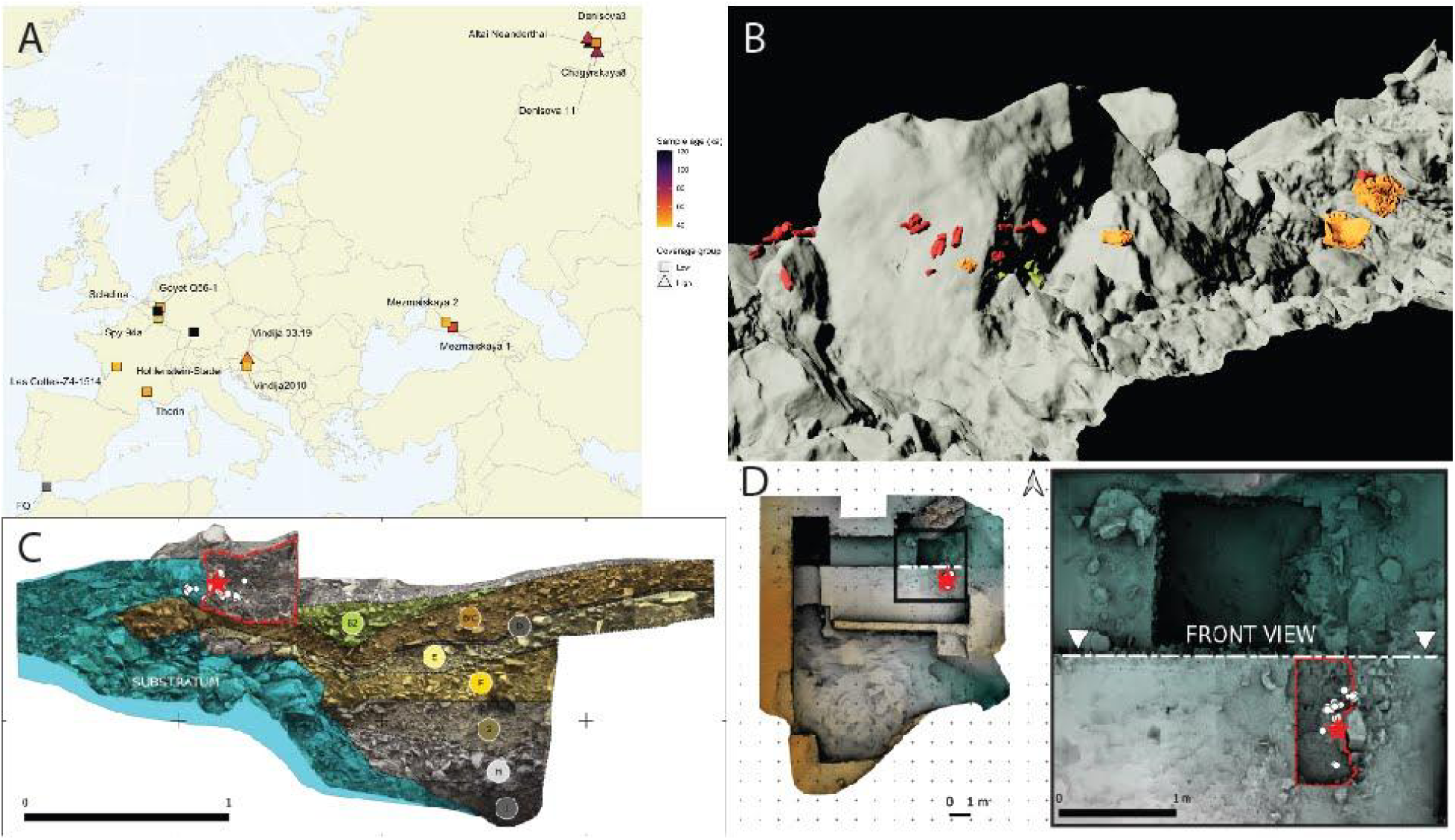
(A) Map showing geographic location, ages, and coverage group of Neanderthal fossils with genome-wide data used in this study. (B) 3D model of the disposition of the Thorin fossils during discovery. Stratigraphic (C) and plan (D) views through Mandrin showing Thorin’s discovery location.

**Fig. 2.**
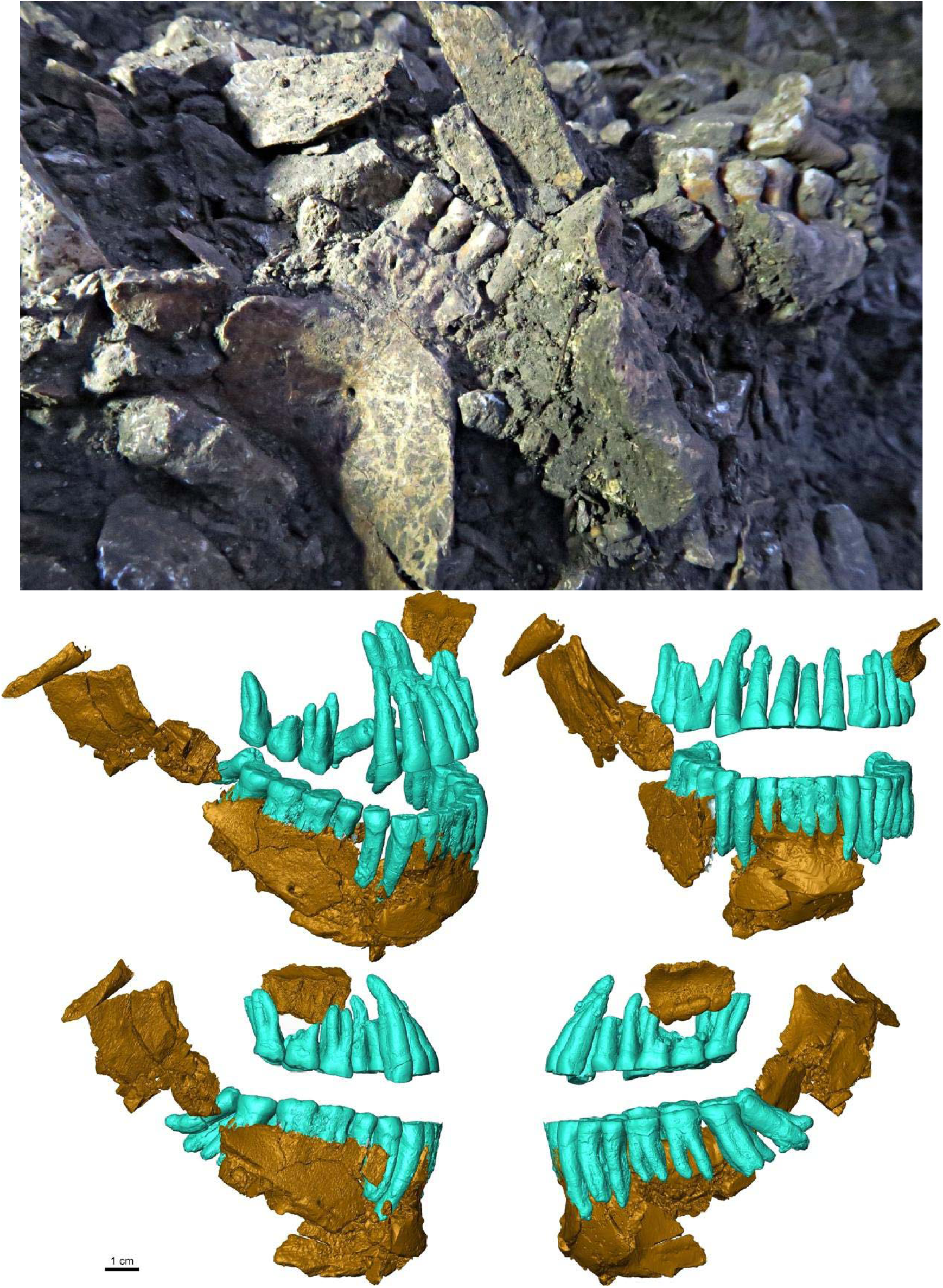
The Thorin Neanderthal. Top. View of the mandible *in situ* when found in September 2019. **Bottom.** Virtual reconstruction of the jaw and dental elements of Thorin in tilted (upper left), anterior (upper right), and lateral (bottom right and left) views.

Overall, the dental morphology of this individual is typical of Neanderthals, with shovel-shaped maxillary central incisors, marked labial convexity on the maxillary lateral incisors that also show a large tuberculum dentale on the lingual aspect of the crown, a well-developed hypocone projected lingually in the maxillary molars, and high root stem/branches ratio (i.e., taurodontism^20, 21^; Fig. S1). Most of the dentition shows advanced occlusal wear associated with hypercementosis at the root apex, notably on the anterior teeth, and the fully developed third fourth molars indicate that it is an adult individual. The advanced occlusal wear is also associated with hypercementosis and exostoses on the maxilla, indicating that the teeth and jaw were under heavy (para)masticatory stress during the life of this individual. Near these cranial elements, the remains of five adult phalanges of the left hand were found (Fig. S2). They showed typical Neanderthal features: ulnar deviation of the pollical distal phalanx and expansion of the distal phalangeal tuberosity^22, 23^. All of the human remains recovered so far are of adult age and the anatomical representation of the different elements is compatible with the presence of a single individual. While the teeth show typical Neanderthal features, the presence of two supernumerary fourth molars is remarkable. Mandibular distomolars are extremely rare in extant humans (around 0.02%)^24^ and, to the best of our knowledge, have not been reported in Pleistocene *Homo* so far, though other kinds of supernumerary teeth have been described in a few instances for Neanderthal and Paleolithic modern humans^25–28^. The aetiology of the presence of distomolars is still debated^24^. Studies of odontoskeletal anomalies found in early-generation hybrids of living primates display a relatively high incidence of distomolars^29, 30^.

Recent analyses of Paleolithic sites in Western Europe suggest that Mousterian lithic industries, traditionally attributed exclusively to Neanderthals, ended 39-41 ka cal. BP^4^. Throughout Eurasia, ten sites have yielded Neanderthal remains directly dated between 50 and 40 ka cal.

BP^16, 31–39^, while only the four French sites, Arcy, Les Cottés, La Ferrassie, and Saint-Césaire, underwent ultrafiltration and provided ages between 45 and 40 ka cal. BP^16, 36, 38, 39^. Neanderthal remains safely attributed to the final stage of their long existence are thus particularly rare and come essentially from sites excavated decades ago^16, 31–39^, often with little or disputable stratigraphic and archeological context.

In order to provide a wider range of less-precious specimens for the destructive process of radiocarbon dating, we screened 80 fragmentary bone remains suspected as deriving from Thorin by Zooarchaeology by Mass Spectrometry (ZooMS) collagen peptide mass fingerprinting^40, 41^ following the methods outlined in ref. *42*. Specimens that yielded spectra matching a Hominidae signature^43^ were radiocarbon dated at the Oxford Radiocarbon Accelerator Unit. Hydroxyproline was extracted for AMS dating to ensure reliability and contamination removal^44^. A selection of hominin remains were also further explored by paleoproteomic sequencing and its ability to distinguish archaic from modern hominin taxa^38^, but comparison with known modern human remains from Holocene deposits proved this approach to be problematic (Supplementary Material 5).

Direct U-series dating and combined U-series – electron spin resonance (US-ESR) dating of Thorin was also undertaken on a fragment of the Neanderthal’s lower left third premolar crown. Additional faunal remains from Level B2 were directly dated using the same approaches. Uranium diffusion and accumulation patterns in the dentine and enamel were obtained prior to the isotopic analysis. According to the diffusion model and the U-series age distribution in the fossils, a minimum age of 43.5±4.1 ka can be assigned to the Neanderthal remains from Level B2. US-ESR modeling yields statistically indistinguishable finite ages of 48 +5/-13 ka and 49 +5/-10 ka for Thorin and the Level B2 fauna, respectively.

We undertook Bayesian modeling of the broader stratigraphic sequence at Mandrin to determine a robust age estimate for Thorin within the PNII levels (C2-B2). The model yielded an age range for Thorin of 51,300-48,900 cal. BP (at 68.2% prob.) and 52,900 - 48,050 cal. BP (95.4% prob.; Fig. 3; see Materials and Methods).

**Fig. 3.**
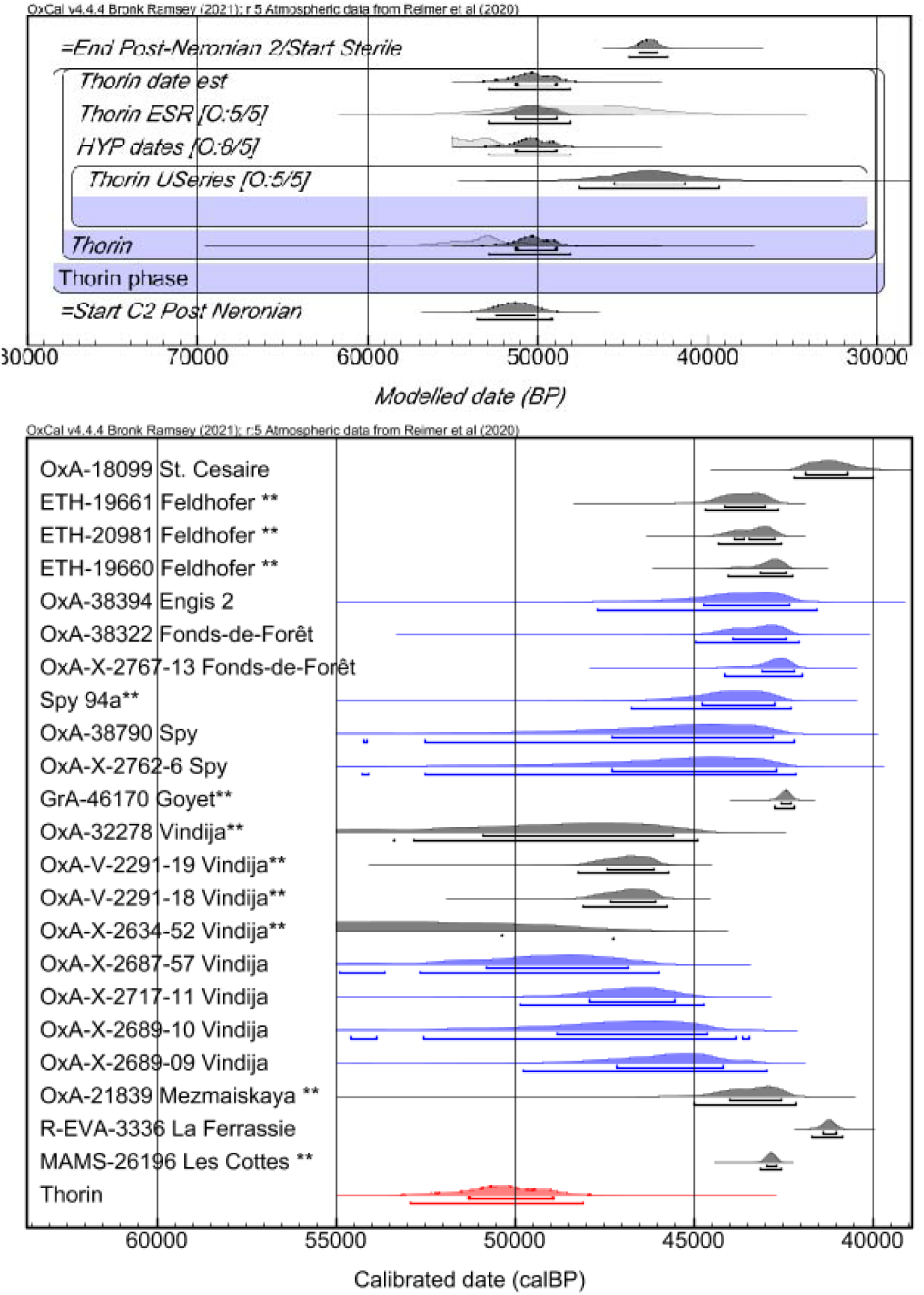
Bayesian modeling of Mandrin’s Post-Neronian II levels containing Thorin. Top. The PN2 phase of the Mandrin Bayesian model used to determine the age of Thorin. The direct ages on Thorin comprise an ESR age, a U-series age constrained as a minimum age and three pooled mean Hydroxyproline AMS ages). Outliers are included in the format [Outlier:posterior/prior]. The boundaries are cross-referenced to the main Bayesian model established for the site, and represent the start and end dates of the PN2. **Bottom.** Comparison of the Thorin modeled age against other ‘late’ Neanderthals. Calibrated likelihoods in blue represent AMS dates obtained using HYP protocols. ** indicates high or low autosomal coverage of the dated specimen. Other likelihoods in black represent dates obtained on bulk or ultrafiltered/purified collagen samples. Dates are after refs. 16 and 31-37.

The carbon, nitrogen, oxygen and strontium isotopic ratios measured on one of the Thorin teeth are fully compatible with an individual living in an open landscape and cold climatic conditions, consistent with the sedimentary characteristics of the C2-B2 deposits and direct dating results, rather than forested and temperate conditions as would have been the case during MIS 5 (Fig. S3; Methods).

### Thorin represents a distinct Neanderthal lineage

A first molar root fragment was used to generate a whole genome sequence from Thorin by performing three sequential DNA extractions (E1, E2 and E3), drastically reducing modern human contamination (Table S9, S10), as well as whole-genome in-solution capture to increase the fraction of endogenous human DNA. Libraries built on raw (non-USER treated) DNA extracts exhibited elevated terminal C>T / G>A substitution rates consistent with authentic ancient DNA data (Figs. S4-S9, Tables S8-S9). However, analyses of contamination rates using mitochondrial DNA and X-chromosome data and grade-of-membership models on the nuclear DNA revealed substantial levels of modern human DNA contamination in the data generated from the first extract E1 (mtDNA-based estimate 13-60%, Table S9; X-based estimate 13-29%, Table S10). We therefore restricted all subsequent analyses to data from extracts E2 and E3, which show re-estimated mtDNA and nuclear contamination rates of <1% and 0.01%, respectively, yielding a final average depth of coverage of 1.3X of the nuclear genome and 561X for the mtDNA.

We rule out the potential of reference and capture bias in our data with D statistics from which we in both cases obtain non-significant D-values (Figs. S17).

Molecular sex determination using reads mapped to the X and Y chromosome showed that the Thorin individual was male. Phylogenetic analyses of the mitochondrial (MT) genome revealed that the Thorin MT genome was most closely related to that of the FQ individual from Gibraltar. Both of them form part of a clade including other recently described European Neanderthal samples (Stajnia Cave, Poland; Galeria de las Estatuas, Spain) and the ∼65 ka Mezmaiskaya 1 individual from the Caucasus, distinct from other late Western Eurasian Neanderthals sequenced to date (Figs. 4A & S10). Analyses of the Y chromosome showed a similar result, with the Thorin sequence diverging prior to the other two male late Neanderthals (Spy94a, Mezmaiskaya 2), albeit with limited bootstrap support (Figs. 4B & S11).

**Fig. 4.**
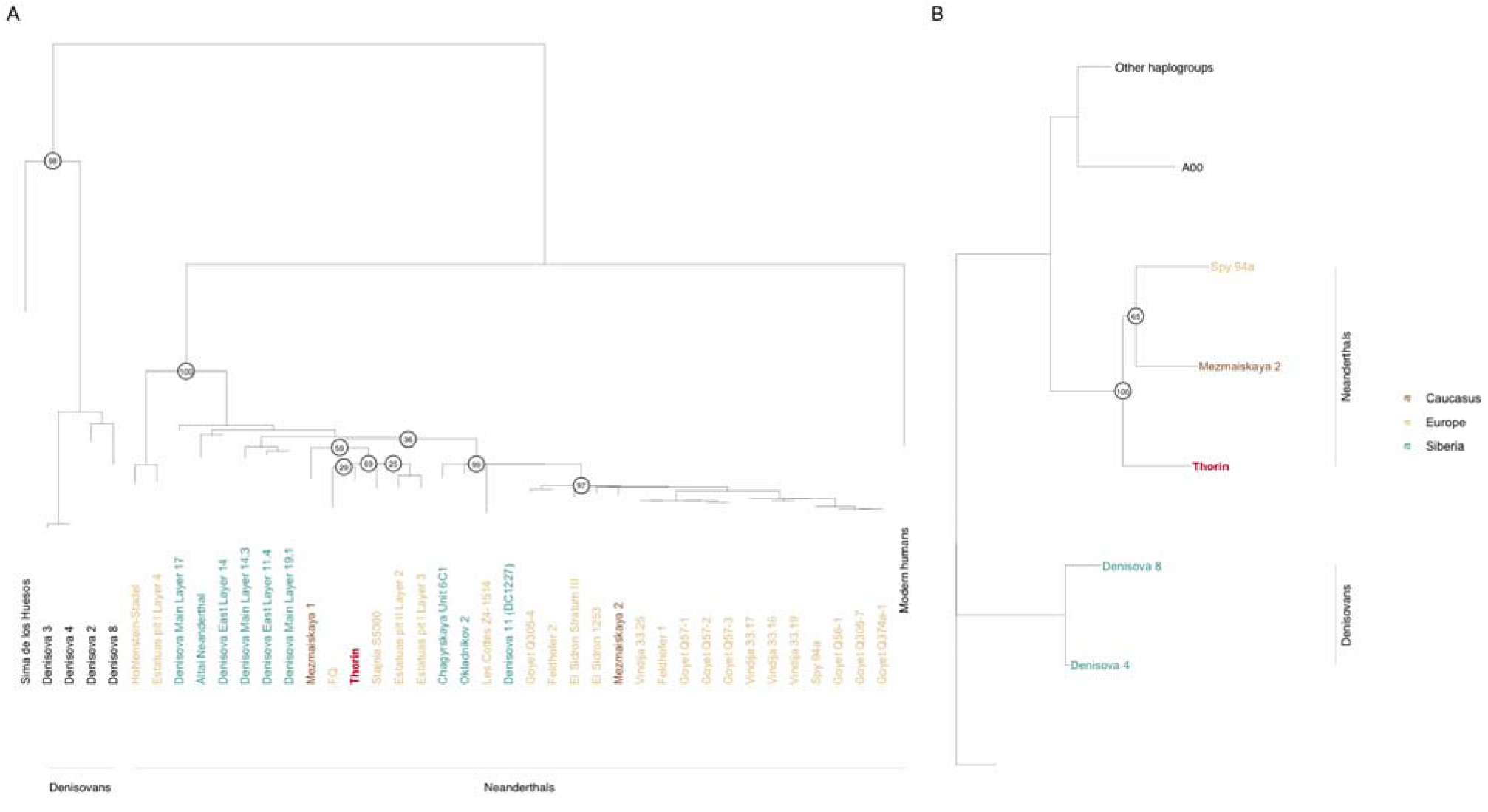
Genetic affinities of Thorin Neanderthal with respect to previously published archaic hominins. (A) Maximum likelihood tree of mitochondrial DNA sequences. (B) Neighbor joining tree of Y chromosome sequences.

Using BEAST2^45^ we obtained a molecular age estimate of ∼100 ka (Table S12, Figs. S12-S13) for Thorin, some ∼50 ka older than the ^14^C, U-series, and OSL ages obtained from the sediment layer from which Thorin was excavated. Similar discrepancies in ages have previously been observed for Chagyrskaya 8^15^ and Stajnia S5000^46^. Notably, directly dated samples used for tip calibration are restricted to the clade of late Neanderthals (Fig. 4A, Table S11) and cover only a shallow part of the entire tree, possibly leading to inaccurate estimates if substitution rates vary across the phylogeny^47^. In order to test this, we carried out an additional BEAST2 analysis including Thorin as an additional calibration point using an age of 50 ka (95% CI: 45– 55 ka), allowing for variation in substitution rates along the tree. The resulting tip ages for Chagyrskaya 8 (70 ka; 95% CI: 48–94 ka) and Stajnia S5000 (77 ka; 95% CI: 53–103 ka), were found to be substantially closer to ages obtained from their respective archaeological contexts (∼60 and ∼50 ka). Similarly, the molecular age estimate of Mezmaiskaya 1 of ∼74 ka also aligned with its previous estimate of 60-70 ka^48^ (Table S12, Figs. S12-S13). The estimated substitution rates remained within a relatively narrow range, suggesting that the initial molecular ages for samples in the Thorin MT clade were likely overestimated. Under this model we estimate a divergence time of the Thorin clade of 123 ka, while we estimate the divergence between Hohlenstein-Stadel and the rest of the Neanderthals to 215 ka and the split between modern humans and all Neanderthals to ∼330 ka (Figs. S12-S13).

We investigated broad population structure among the low coverage Neanderthals and Chagyrskaya 8 by projecting them onto a principal component analysis (PCA) of Vindija 33.19, Altai Neanderthal and Denisova 3, the three deepest diverged archaic lineages with high quality genomes currently available. The projected individuals formed a cline towards Vindija 33.19, consistent with their previously reported sharing of a more recent common ancestor than with the Altai Neanderthal (Fig. S14). Interestingly, the placement of Thorin fell within the cline but further from Vindija 33.19 than any other late Neanderthal individual, suggesting a more distant relationship to Vindija 33.19. D-statistics confirmed that Neanderthals from Europe, the Caucasus, and Siberia younger than 80 ka shared significantly more alleles with Vindija 33.19 than with Thorin, and that the Thorin lineage forms an outgroup to those lineages (Figs. 5 & S15-S19). The exception was the Neanderthal sample FQ from Gibraltar^49^, which showed a weak but significant signal of excess allele sharing with Thorin, consistent with their closely related MT sequences (Figs. 6 & S15-S19). Furthermore, Thorin does not show excess allele sharing with modern humans in comparison to all other west Eurasian Neanderthals, indicating that the lineage interbreeding with modern humans diverged prior to the Thorin lineage, and ruling out the possibility of recent interbreeding with early modern humans at Mandrin cave^1^ (Fig. S17).

**Fig. 5.**
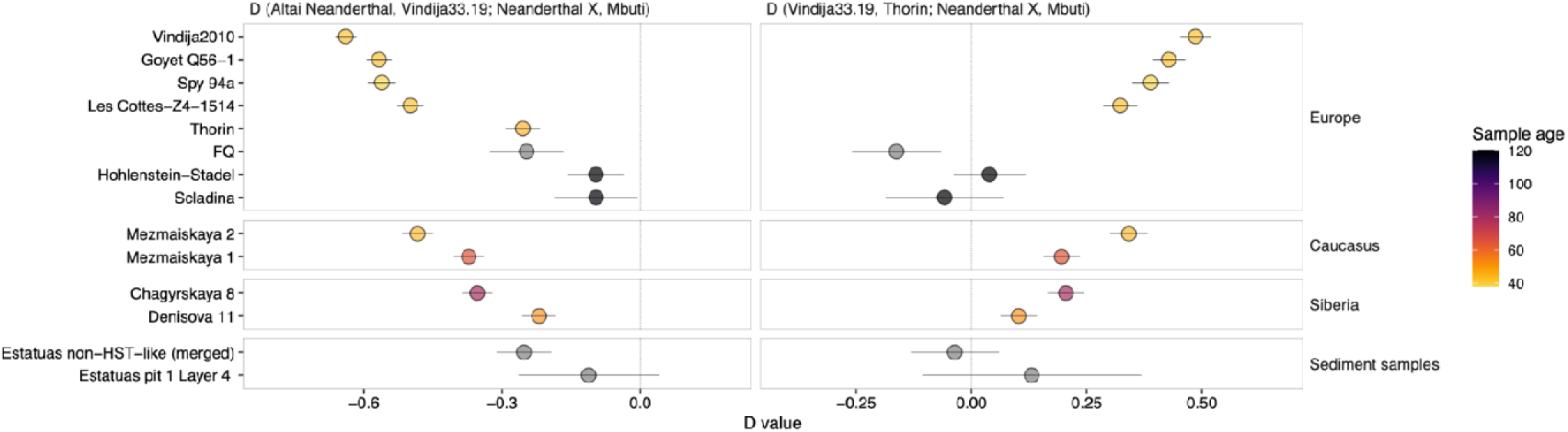
Genetic affinities of Thorin. *(left)* D-statistics of the form D(Altai Neanderthal, Vindija 33.19; Neanderthal X, Mbuti), showing that all Eurasian Neandertal samples share more alleles with Vindija 33.19 than with the Altai Neandertal. Thorin shares relatively more alleles with Vindija 33.19 than early European Neanderthals of the putative first radiation (HST, Scladina, Estatuas pit 1 Layer 4), but less than the second radiation (Mezmaiskaya 1, Chagyrskaya 8). (*right)* D-statistics of the form D(Vindija 33.19, Thorin; Neanderthal X, Mbuti), showing that Neanderthals from Europe, the Caucasus, and Siberia younger than 80 ka share more alleles with Vindija 33.19 than with Thorin. The exception is the Gibraltar Neanderthal sample (FQ), which shows increased affinity with Thorin. Error bars indicate 3 ✕ standard error (|Z| = 3).

**Fig. 6.**
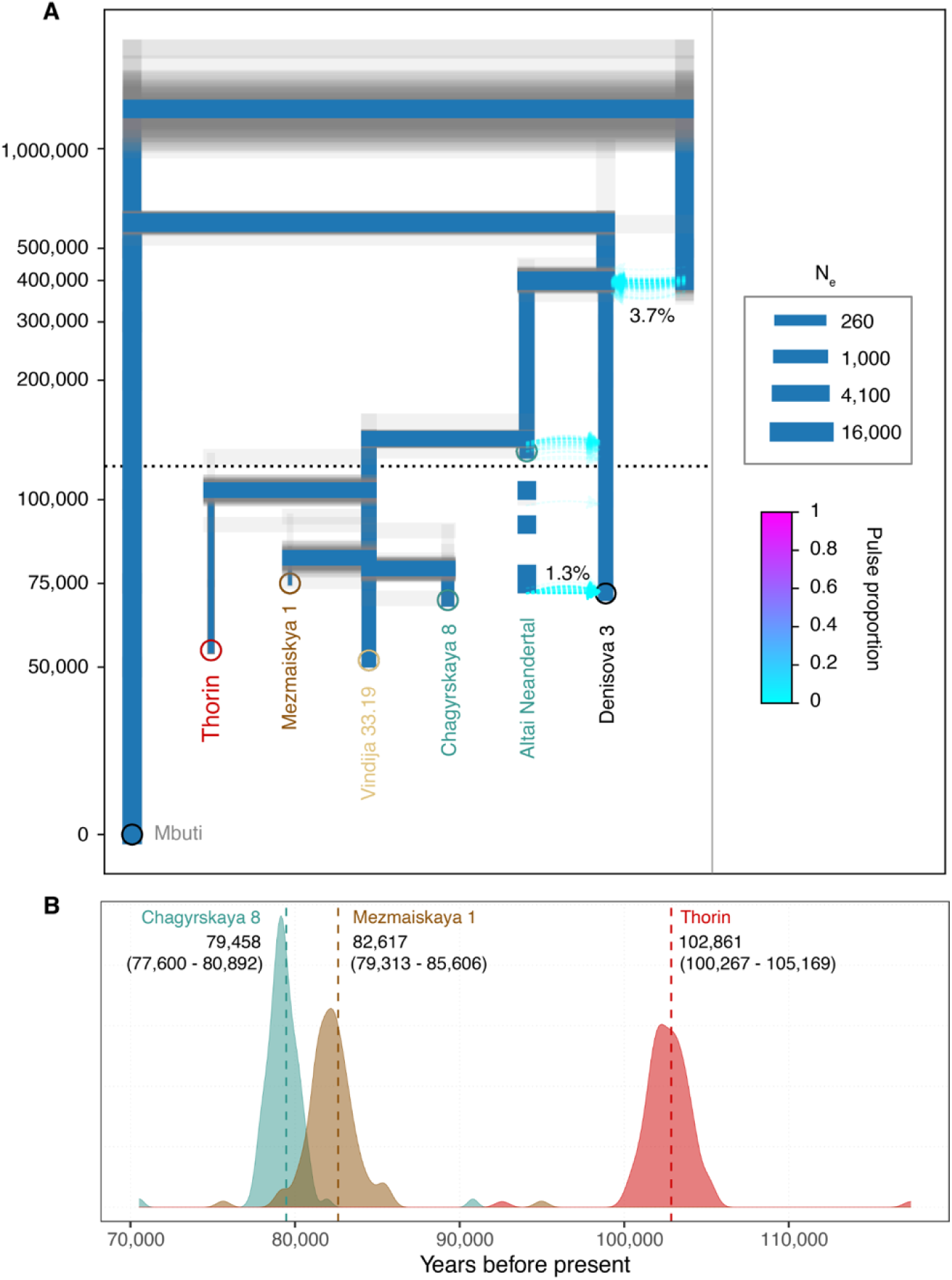
Demographic history of the Thorin lineage. **(A)** Best-fitting demographic model relating Thorin to other Neanderthal and Denisovan genomes. Blue branches show point estimates, whereas gray transparent branches indicate estimates obtained using 100 nonparametric bootstrap replicates. **(B)** Point estimates (dashed line) and density of 100 parametric bootstrap replicates for divergence time parameters of Thorin, Mezmaiskaya 1 and Chagyrskaya 8 Neanderthal genomes from the late European Neanderthal Vindija 33.19.

We carried out demographic modeling using the site-frequency-spectrum based approach implemented in *momi2*^50^, which allows the placement of low coverage individuals onto a scaffold inferred from high quality genomes. We first fit a scaffold demography including the three high coverage Neanderthals (Altai Neanderthal, Chagyrskaya 8, Vindija 33.19) as well as the Denisovan, incorporating previously inferred demographic events^8^. The low coverage samples Thorin and Mezmaiskaya 1 were then added to this scaffold, allowing for a divergence from the Vindija 33.19 lineage at any point after the split from the Altai Neanderthal. The best-fit model indicates a divergence of the Thorin lineage from Vindija 33.19 at 102,861 years ago (95% CI 100,267 - 105,169), considerably earlier than those of Mezmaiskaya 1 (82,617 ya; 95% CI 79,313 - 85,606) or Chagyrskaya 8 (79,458 ya; 95% CI 77,600 - 80,892; Fig. 6), and consistent with results from D-statistics and mtDNA.

Using a novel approach to detect runs of homozygosity in low coverage Neanderthal genomes, we found evidence for increased homozygosity in the Thorin genome compared to other late European Neanderthals. Thorin harbors ∼7% of its genome in homozygous segments of 5Mb or longer, including 45 Mb (∼1.5%) in segments longer than 20Mb indicative of recent inbreeding (Figure 7). Taken together, our results suggest small group sizes and long-term genetic isolation of the Thorin population from other late Neanderthal populations with genomic data available.

**Fig. 7.**
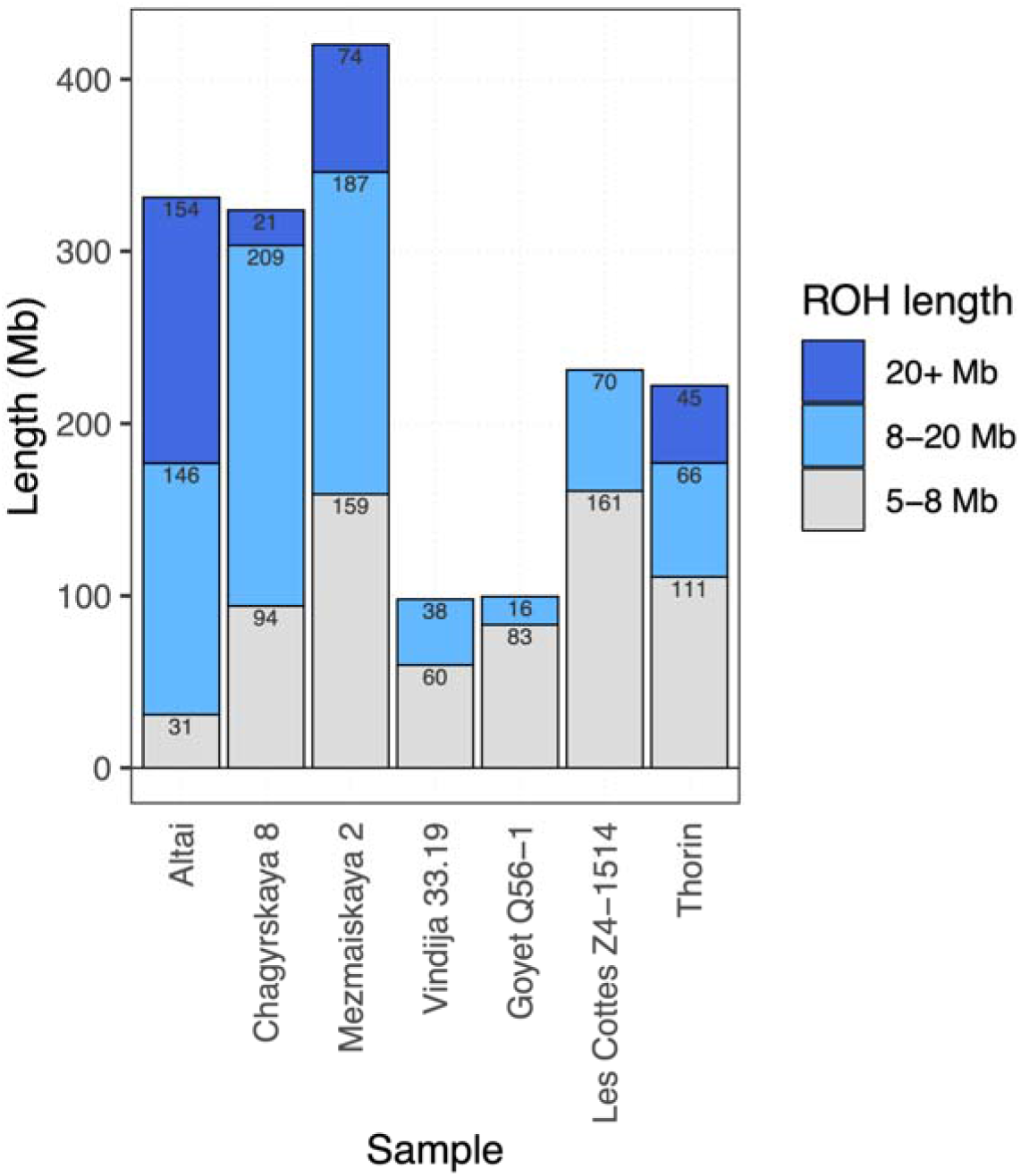
Runs of homozygosity in Neanderthals. Bar plot showing cumulative total length of ROHs ≥ 5Mb in Thorin and other Neanderthal genomes with average genomic coverage ≥ 1.5X. ROH length classes are distinguished by bar colors, with total length in each class indicated.

### Other isolated lineages present 50 ka?

We further investigated the possibility of population turnover in Europe during the late Neanderthal period. Using D-statistics testing whether the late Caucasus lineage of Mezmaiskaya 2 forms an outgroup to other late European Neanderthals, we find evidence of gene flow with a “deep” Neanderthal lineage in the ∼43 ka Les Cottés Z4-1514 sample from France (Fig. S18). Interestingly, this individual carries a mtDNA lineage most closely related to the Siberian Neanderthals from Okladnikov and Chagyrskaya caves, diverging earlier than the clade of Vindija-like late Neanderthals sampled to date (Fig. 4). Demographic modeling of Les Cottés Z4-1514 and Mezmaiskaya 2 onto the previous best-fitting model revealed that a model with gene flow into Les Cottés Z4-1514 from an unsampled ghost lineage diverging ∼89 ka provided a significantly better fit than one without gene flow (Fig. S24). An alternative model involving a ghost lineage constrained to diverging from the Thorin lineage also yielded a poorer fit, with a divergence time of the ghost lineage close to the diverging of the Thorin lineage (Fig. S24). Our results thus suggest the presence of at least two deeply divergent and isolated lineages in close geographic proximity during the late Neanderthal period, subsequently partially replaced by an expansion of Vindija-like lineages into western Europe within the last 10,000 years of their existence. Interestingly, the eastern European late Neanderthal from Mezmaiskaya cave (Mezmaiskaya 2) also shows high levels of homozygosity (Figure 7), suggesting small group sizes were likely also common among late Neanderthals outside the expanding Vindija-like population.

## Discussion

Thorin is the most complete Neanderthal individual found in France since 1979^17^ and falls amongst a group of other Neanderthals dating to the last millennia of their existence in western Europe. So far population genetic analysis of other late Neanderthals has indicated they belonged to a single metapopulation with no significant evidence of deep population structure among them^16^. The genome of Thorin sheds new light on the population structure of late Neanderthals as our genomic analyses demonstrate that Thorin belongs to a deeply diverging European Neanderthal lineage, representing a remnant of earlier European Neanderthals. Interestingly, the divergence of this lineage began at ∼100-105 ka, during the MIS 5 interglacial, a period that saw fast climatic and environmental changes across Eurasia and repopulation by warm adapted fauna through the continent^51, 52^. The timing of this divergence also coincides with a period of population replacement detected in northern Spain among Neanderthal populations^14^.

Our analysis testing for gene flow between Thorin’s lineage and other known Neanderthal and modern human lineages suggests the existence of an isolated group of late Neanderthals in western Europe 50 ka. This population is associated with a distinctive PNII lithic tradition^1^, which is continuously attested in the last four Mousterian levels of Mandrin (Levels C2 to B2), from 52.9-43.0 ka at 95% CI, overlapping with the final disappearance of Neanderthal populations in Eurasia^1, 4^. Thorin therefore likely belonged to one of the last representative Neanderthal populations in this area of Mediterranean France, and poses the first direct genomic evidence of deep population structure among late European Neanderthals. The genetic relationship observed between Thorin and FQ (also indicated in demographic modeling, Supplementary Note 4) indicates that the Gibraltar Neanderthals might have been members of an extended southwest European metapopulation, and raises the possibility of a much later dating for those individuals than previously anticipated^49^. However, due to sparsity of data from FQ, we are unable to draw further conclusions hereof.

The genetic differences between Thorin and the other Western European Neanderthals may signify a major process of population replacement following, or related to, the expansion of anatomically modern humans through Europe. Interestingly, Thorin corresponds to the phase of Neanderthal reoccupation of Grotte Mandrin after the earliest modern human incursions in Europe^1, 2^. The millennia-long genetic isolation of the Thorin-lineage raises new questions of relevance to the Neanderthal extinction debate and the types of interactions between the last Neanderthals and early *H. sapiens* in Europe. Additional DNA analyses and secure direct dating of late Neanderthal remains are now crucial to understand whether this population was only locally spread -in the middle Rhône Valley-or if the Thorin lineage was more widely distributed across Europe, as suggested by the Gibraltar connections. The sedimentary autosomal DNA data from the Galeria de las Estatuas population was unfortunately not of sufficient coverage to establish closer affinity with Thorin. However, the sampling locations of European Neanderthals within the mtDNA clade of Thorin, from Iberia (Gibraltar, Galeria de las Estatuas) and southern France to Poland (Stajnia) would support a broader distribution, and be consistent with a suggested radiation of Neanderthal populations ∼105 ka. While it is commonly inferred that the interactions between the first *H. sapiens* and the last Neanderthals may have played an important role in the latter’s extinction in Europe, the unexpected identification of an hitherto unrecognized late Neanderthal population reveals a much more complex population structure among late Neanderthals and raises new lines of questions to further explore their social or ethological organization, which potentially could have played an important role in their later extinction.

Besides the lineage that is represented by Thorin, our demographic modeling provides indirect evidence of another deeply diverged “ghost” lineage present through the French Neanderthal Les Cottés. Our demographic modeling suggests that the introgressing lineage diverged some time after the Thorin lineage, closer to the divergence of Mezmaiskaya1 and the Siberian individual Chagyrskaya8 from the Altai region, with which Les Cottés also shares a closely related MT lineage. Whether this ghost lineage forms part of an as yet unknown further radiation of lineages after 100 ka but before the classical late Neanderthals remains unknown without a denser sampling of genomic data from around that time period. Our results nevertheless suggest a minimum of two distinct Neanderthal lineages present in Europe during the late Neanderthal period. In the absence of any detectable gene flow between Thorin and other Neanderthal lineages after its divergence, we conclude that Thorin represents a lineage that has stayed isolated for ∼50 ka. Deep cultural and technical specificities distinguishing Rhône Valley late Mousterian industries have been long proposed^18, 19, 53^, underlining that from MIS 5 to 3 these French Mediterranean Neanderthal societies possessed a distinct technical background. These cultural traits distinguishing Neanderthal societies from neighboring regions can now be paralleled with deep genetic isolations among these societies. Our results thus also shed light onto the social organization of Neanderthals, suggesting that small isolated populations with limited, and potentially without, inter-group exchange as a possibly more general feature of Neanderthal social structure.

## Acknowledgements

We deeply acknowledge the Service Régional de l’Archéologie Auvergne Rhône-Alpes, the city of Malataverne, and all of the students/volunteers that supported the 30 years of continuous field research in Grotte Mandrin. The authors acknowledge Professors E. Willerslev and L. Orlando for fruitful discussions and continuous support. We thank the laboratory technicians of the Danish National High-throughput DNA Sequencing Center, as well as Tina B. Brand, Pernille V. S. Olsen and Jesper Stenderup from the Lundbeck Foundation Center for GeoGenetics for technical assistance. Thanks also go to Manasij Pal Chowdhury for assistance with the paleoproteomics data availability.

## Author contributions

L.S. and L.M. lead the scientific project and excavated the site. Microtomographic-based data were collected and analyzed by C.Z. L.S. analyzed the lithic technical systems. H.Co., O.D. and C.Z. studied human remains. Spatial analyses were performed by P.Y. and X.M. H.Ca. recorded and analyzed stratigraphy and deposit dynamics. Radiometric dating analyses and Bayesian age modeling were performed by T.H., R.J-B, L.A., M.D, M.F., T.D. and D.C. H.B. performed and interpreted the isotopic measurements on bones and teeth. M.S. coordinated the ancient DNA analyses. A.S.O performed the ancient DNA laboratory work and sequencing.T.V. and M.S. analyzed the genomic data. L.S., C.Z., M.S., T.V., and J.E.L. wrote the manuscript with contributions from all other authors. A.S.O., T.V., and L.S. wrote the supplementary materials with contributions from all other authors.

## Data and materials availability

Include a note explaining any restrictions on materials, such as materials transfer agreements. Note accession numbers to any data relating to the paper and deposited in a public database; include a brief description of the data set or model with the number. If all data are in the paper and supplementary materials include the sentence “All data is available in the main text or the supplementary materials.” All data, code, and materials used in the analysis must be available in some form to any researcher for purposes of reproducing or extending the analysis.

## Supplementary Materials

Materials and Methods

Supplementary Note 1. Site and fossil descriptions

Supplementary Note 2. Quality control and contamination estimates

Supplementary Note 3. Uniparental markers: Mitochondrial & Y-chromosome analysis

Supplementary Note 4. Population genetic analyses

Figures S1-S24

Tables S1-S36

Supplementary References

