## Supplementary Information for "A late Neanderthal reveals genetic isolation in their populations before extinction"

**This PDF file includes:**

Materials and Methods

Supplementary Note 1. Site and fossil descriptions

Supplementary Note 2. Quality control and contamination estimates

Supplementary Note 3. Uniparental markers: Mitochondrial & Y-chromosome analysis

Supplementary Note 4. Population genetic analyses

Supplementary Note 5. Paleoproteomic analyses

Tables S1 to S15

Figs. S1 to S20

**Materials and Methods**

**ZooMS**

ZooMS collagen fingerprinting was carried out on 33 spatially-plotted bone fragments discovered in contact with Thorin and considered as potential hominin by basic characteristics of bone size, thickness, etc. Following ref. 42, this involved removal of collagen through decalcification with 0.6 M hydrochloric acid (HCl) overnight, ultrafiltration into 50 mM ammonium bicarbonate using 10 kDa molecular weight cut-off filters, and digestion with sequencing grade trypsin at 37^o^C overnight. The digests were then acidified to 0.1% trifluoroacetic acid (TFA) and then ziptipped (Varian OMIX C18 pipette tips) for peptide purification, being eluted in 50% acetonitrile (ACN) in 0.1% TFA and dried to completion by centrifugal evaporation. Samples were then rehydrated with 10 µL 0.1% TFA and 1 µL co-crystallized with an equal amount of 10 mg/mL alpha-cyano hydroxycinnamic acid in 50% ACN/0.1% TFA and allowed to dry. Peptide mass fingerprints were then acquired using a Bruker Ultraflex II Matrix Assisted Laser Desorption Ionisation Time of Flight Mass Spectrometer over the range *m/z* 700-3,700 and compared with reference spectra for humans^41,54^. The acid-insoluble pellets from specimens identified as hominin were then further processed for radiocarbon dating.

**Radiocarbon Dating**

Samples from the Thorin specimen were AMS dated at the Oxford Radiocarbon Accelerator Unit (ORAU) at the University of Oxford. Collagen was initially extracted using the ‘AG’ protocol, comprising a simple demineralisation and gelatinisation. Following this the collagen was treated using the method outlined in ref. 44. This comprised an initial acid hydrolysis in 6M HCl. Underivatized amino acid solutions were dried down and then resuspended in 0.1 M NaOH. Following this they were separated using a Varian prep-Liquid Chromatography system. The hydroxyproline was collected in MilliQ^TM^ water, then concentrated and dried using a Genevac EZ‐2 Plus vacuum evaporator. Following this the samples were combusted via a PDZ-Europa Robo-Prep combustion elemental analyzer coupled to a PDZ-Europa 20/20 mass spectrometer operating in continuous flow mode using He gas as a carrier. C/N atomic ratios as well as δ^15^N and δ^13^C values, %carbon and nitrogen values, were obtained (Table S1). Following combustion the samples were graphitised and reacted with an iron catalyst in an excess of H_2_ at 560℃^55^. The graphite was then AMS dated. Measurements were background corrected for the presence of trace carbon on the HPLC^44^.

**Uranium series direct dating of teeth**

Two complete faunal teeth and one fragment of a *Homo neanderthalensis* tooth were submitted for U-series dating. The teeth showed very little signs of diagenetic alteration with superficial discoloration of the outer enamel and a small amount of sediment insertion in natural cracks. The sample set consisted of a small bison premolar and a fragmented Neanderthal tooth (half upper crown of a molar or premolar) from the Thorin specimen, both associated with Level B2, and a caprinid molar (1996-E0-B#65) from Level B3. Teeth were sectioned in half using a high precision slow speed rotary diamond saw, exposing dentine and enamel tissues. The exposed surface of each sample was then polished to 5 microns smoothness to offer a clean ablation surface.

Uranium-series dating of the teeth was undertaken by laser ablation multi-collector ICP-MS at the Biomics Laboratory of the Geoarchaeology and Archaeometry Research Group (GARG) facility, Southern Cross University. Laser ablation was performed with a New Wave Research 213 nm laser, equipped with a TV2 cell. Thorium (^230^Th, ^232^Th) and uranium (^234^U, ^235^U, ^238^U) isotopes were measured on a Thermo Neptune XT multi-collector ICP-MS mounted with jet sample and x-skimmer cones. All five isotopes were collected in static mode, with both ^234^U and ^230^Th collected in the ion counter and CDD, respectively. Helium flow rate and ICP-MS parameters were tuned with NIST610 element standard to derive a ^232^Th/^238^U ratio for this standard greater than 0.85 and thus minimize differences in fractionation between Th and U. For tuning, a fluence of 13.1 J/cm2, pulse rate of 20 Hz, spot size of 110 μm and scan speed of 5 μm/s were used. This yielded 1.75V of ^238^U and 1.50V of ^232^Th on NIST610.

Teeth were ablated using rasters of 4min 54s each (two passes of ~750 μm long). Before and after each sample, NIST612, MK10 and MK16^55^ standards were measured, as well as a fossil *Hippopotamus* tooth with known isotopic ratios. ^234^U/^238^U and ^230^Th/^238^U isotopic ratios were corrected for elemental fractionation and Faraday cup/SEM yield by comparison with MK10 coral for which ratios were previously characterized internally by solution analysis. Concentrations of U and Th were determined using NIST612 glass as a calibration standard. Background subtraction, concentration quantification and ratio corrections were performed using Iolite™ software. The corrected (^234^U/^238^U) and (^230^Th/^238^U) isotope ratios for the secondary standard (MK16 coral; 1.110 ± 0.01 and 0.809 ± 0.042, respectively) were within error of the values determined by solution analysis (1.110 ± 0.002 and 0.764 ± 0.007). The calculated closed-system ^230^Th-U age for MK16 was 120 ± 5.2 ka (2σ, n=8), within error of the value determined by solution analysis (124 ± 2 ka). The hippopotamus tooth was used as a control on matrix effect.

Analyses of the caprinid, bison and Thorin specimens consisted of 8, 14 and 20 rasters respectively, each two passes of 750 μm long. Each raster was averaged to obtain one isotopic and age data point. The caprinid tooth didn’t offer any exploitable results (thorium concentration was under detection limit) and therefore ages were not calculated and are not reported. Both the bison tooth and Thorin fossil tooth exhibit similar isotopic values, corroborating the same stratigraphic provenance and similar burying condition. Parts of the Neanderthal tooth fragment showed some significant ^232^Th incorporation, as well as some leaching towards the edge close to the fractured dentine. The human tooth especially seems to have suffered from a higher migration of detritic thorium, which can be explained by the fractured and fragmented structure of the tooth. Both samples showed rather heterogeneous distribution of U-series isotopes, with clusters of accumulation towards natural cracks and the enamel dentine junction. Large areas of the teeth had obvious diagenetic processes with low to severe impact on the uranium series isotopes distribution and age calculations. Overall, both uranium and thorium concentrations were rather low, and several areas of the tooth were at detection limits, hampering our ability to obtain exploitable data. This also explains the relatively large uncertainty associated with the measurements and propagated in the final age uncertainty. Both fossils were recovered from the same stratigraphic layer and the results indicate a similar deposition and age bracket (Table S2). Despite a complex isotopic distribution, the diffusion pattern remains typical of what is usually observed in dental tissues. The U-series age of 43.5 ± 4.1 ka (2-sigma) for Thorin corresponds to the time at which the uranium migrated into the dental tissues. In that regard, the U-series age should be considered a minimum age for the sample, since some time might have passed between the death of the individual and the initial incorporation of uranium into the tooth. Yet, mapping the spatial isotopic distribution within the teeth gives a better understanding of the uranium uptake pathways and history, ultimately increasing the analytical protocol and the correctness of the age by measuring in accumulation clusters. Unless complicated burial processes and history have altered significantly the initial diffusion of uranium into the samples, it appears likely that 43.5±4.1 ka is close to the “true” age of the stratigraphic deposition.

**US-ESR direct dating**

An enamel fragment from each tooth was separated using a hand-held diamond saw following the protocol developed in ref. 56 and stripped of the outer ~100 microns ±10% on each side. Fragments were mounted into a parafilm mold within a Teflon sample holder to record the angular dependency in the ESR response^58,59^. Fragments were then measured at room temperature on a Freiberg MS5000 ESR X-band spectrometer at a 0.1mT modulation amplitude, 10 scans, 2mW power, 100G sweep, and 100KHz modulation frequency for ESR dating. Irradiation was performed with the Freiberg X-ray irradiation chamber, which contains a Varian VF50 X-ray gun at a voltage of 40KV and 0.5mA current on the fragment exposed to X-rays without shielding (apart from a 200um Al foil layer^60^ ). Each fragment was irradiated, following exponentially increasing irradiation times (around 90s, 380s, 900s, 1800s, 3600s, 7200s, 14400s, 25000s and 50000s, although the exact irradiation time and dose rate varied for each sample). For each irradiation step, the energy output of the X-ray gun is recorded at the beginning and end and averaged, which allows correction for the dose rate received by the sample. For each irradiation step the fragment was measured over 180^o^ in x, y and z-configurations with a 20^o^ step^61,62^. ESR intensities were extracted from T1-B2 peak-to-peak amplitudes on the merged ESR signal. Isotropic and baseline corrections were applied uniformly across the measured spectra^61^. The amount of NOCORs was estimated using the protocol described in ref. 58, yet no influence was observed on the spectra after x-ray irradiation steps. The ESR dose response curves were obtained by using merged ESR intensities and associated standard deviations from the repeated measurements over one orientation only. Fitting procedures were carried out with the MCDOSE 2.0 software using a Markov Chain Monte Carlo (MCMC) approach based on the Metropolis-Hastings algorithm^62^. D_E_ values were obtained by fitting a single saturating exponential (SSE) at the appropriate maximum irradiation dose (D_max_), following the recommendations of ref. 63. The dose equivalent of the Neanderthal “Thorin” and fauna Bison teeth from B2 was estimated to be 64.1 ± 4.5 Gy and 67.0 ± 5.6 Gy respectively. Using this D_E_ and the parameters shown in Table S3, the US-ESR ages were unable to be modeled, likely because of U leaching and making the ESR ages younger or equivalent to the U-series ages. Thus, the equivalent dose for each sample was offset, as follows: 66.3 +2.3/-6.7 Gy for Thorin and 68.8 +3.8/-7.4 Gy for the Bison tooth to allow realistic modeling and creating asymmetrical uncertainties. Using this protocol, the US-ESR age obtained for Thorin is 48 +5/-13 ka and 49 +5/-10 ka for the Bison tooth. If we assume a closed system (CS) model, Thorin's age estimation becomes 56±41 ka and the Bison fauna age becomes 56 ± 39 ka, which would correspond to maximum ages for these fossils. Additionally, a simple early uptake (EU) model gives an age estimation of 42±5 ka for the fauna. We also modeled the US-ESR ages of Thorin and the bison tooth by considering dosimetry from the C2 layer or a mix B2/C2 values. Using the parameters from C2 or average C2/B2, Thorin age estimates are slightly shifted to 51+4/-14 ka and 50+5/-13 ka, and the bison tooth to 51+4/-10 ka and 50+5/-10 ka, respectively. Therefore, all age estimates are statistically indistinguishable regardless of the stratigraphic association with B2, C2 or an average B2/C2. For the Bayesian model, all systematic errors were removed and all associated uncertainties defined at 1-sigma. This resulted in a US-ESR age estimate of 48 ± 4 ka for Thorin, which has been included as the likelihood in the OxCal model.

The sediment elemental concentrations, external beta and gamma dose rate contributions, and water content shown in Table S3 correspond to those obtained for OSL sample X6717 from B2^1^. The external beta dose rates have been calculated from the U, Th and K contents measured on a portion of sediment sub-sample (~8 g) by ICP-MS and ICP-AES. The external gamma dose rates were determined from carbon-doped aluminum oxide dosimeters buried in close proximity to the location of the sediment sample for a period of 396 days. The cosmic dose rate has been calculated for the specific burial depth of the fossils according to ref. 64, taking into account the altitude, geomagnetic latitude, density of sediment overburden, and the time-averaged geometry of bedrock overburden.

**Bayesian model**

We built a simple Bayesian model using OxCal 4.4.3^65^ and the INTCAL20 calibration curve^66^ to determine a modeled age for Thorin. Thorin is linked stratigraphically with the PN2 (Post-Neronian 2) levels of the site, encompassing levels C2 to B2. For this reason we used the dated boundaries from the Bayesian model derived in ref. 1 as cross-referenced constraints on the age in a single Phase model aimed at providing a probabilistic density function for the age of the Thorin remains. For Thorin’s age estimate we used an error-weighted mean of the three HYP radiocarbon ages in fraction modern of 0.0007 ± 0.0007 fM (T=2.64, ꭓ2=5.99, df.=2), as well as the U Series and ESR age estimates constrained as minimum and direct ages respectively. Outliers were set at 0.05 probability within a General Outlier_Model^67^. All posterior outliers were set at <5%. The CQL code for this is below.

The results of the modeling are shown in Fig. 4. We used a Date command embedded with the phase to determine an age range for Thorin at 51,300-48,900 cal. BP (at 68.2% prob. and 52,900-48,050 cal. BP (95.4% prob.)).

***Bayesian Model CQL code for Thorin age***

Sequence()

{

Boundary("=Start C2 Post Neronian");

Phase("Thorin phase")

{

Combine("Thorin")

{

Before()

{

Date("Thorin USeries",N(2021-43500,2050))

{

Outlier("General", 0.05);

};

};

R_F14C("HYP dates", 0.0007, 0.0007)

{

Outlier("General", 0.05);

};

Date("Thorin ESR",N(2021-48000,4000))

{

Outlier("General", 0.05);

};

Date("Thorin date est");

};

};

Boundary("=End Post-Neronian 2/Start Sterile");

};

**X-ray microtomographic scanning of the teeth**

The teeth and jaw fragments of Thorin were scanned using the X-ray microfocus instruments (X-µCT) Phoenix Nanotom 180 (FERMAT Federation from the Inter-university Material Research and Engineering Centre, UMR 5085 CNRS, University of Toulouse) and GE v|tome|x s (Placamat platform, UMS 3626 CNRS, University of Bordeaux). Acquisitions were performed according to the following parameters: 100-130 kV, 170-250 µA, 1440-2550 images taken over 360° (0.14-0.25° of angular step), 0.1 Cu filter. The final volumes were reconstructed with a voxel size of 19.5-26.3 µm for the isolated teeth and 48.1-50.0 µm for the jaw fragments.

**Isotopic analyses**

The goal of performing stable isotopic analyses of tooth enamel (carbon *d*^13^C, oxygen *d* ^18^O, strontium ^87^Sr/^86^Sr) and bone or dentine collagen (carbon *d* ^13^C, nitrogen *d* ^15^N) was to gain information on the paleoenvironmental context of Thorin, its position in the large mammal trophic system and its mobility pattern. These newly obtained data could be compared with those obtained for faunal remains from sites in the same region and for other late Neanderthal specimens in western Europe. Since carbon, nitrogen and oxygen isotopic ratios depend on the type of vegetation at the basis of the food webs (open or forested), aridity, plant productivity, temperature, geological bedrock, and, for humans, on the trophic position and the prey selection^68,69^, this multi-isotopic approach allows us to refine the palaeoenvironmental context of this Neanderthal and to confirm its chronological attribution. For this study, fossil tooth material from faunal and human specimen from Grotte Mandrin were analyzed (see below) and published data for the sites of Payre in the same region were used for comparison of carbon, oxygen and strontium isotopic values of tooth enamel^70-72^, while sites such as Aldène Cave, Mialet Cave, Abri du Maras, Trou de la Mère Clochette, Saint-Marcel Cave were used for comparison of carbon and nitrogen isotopic values of bone collagen^73-77^ as well as previously published data on identified fauna from Grotte Mandrin^4^ were used for comparison of the carbon and nitrogen isotopic values of bone and tooth collagen.

***Carbonate isotopic analysis***

For carbonate isotopic analysis, tooth enamel was sampled from animal teeth from levels B and C in Grotte Mandrin, for the following large mammal taxa: bison *Bison priscus* (n = 4), red deer *Cervus elaphus* (n = 6), reindeer *Rangifer tarandus* (n = 3) and horse *Equus cf. germanicus* (n = 4). In addition, one tooth enamel fragment from Thorin was also analyzed, as well as two modern human teeth from Neolithic context in the same cave as comparison.

For the newly analyzed specimens, about 5 to 10 mg of tooth enamel were drilled using a rotating tool with diamond coated drill bit and the powder was then pretreated following the protocol from ref. 78 to remove possible organic and carbonated contamination. Subsequently, the C and O isotopic analysis was performed at the Department of Geosciences of the University of Tübingen as follows: pretreated enamel was reacted with 100% H_3_PO_4_ for 4 h at 70^◦^C using a MultiFlow-Geo interfaced with an Elementar IsoPrime 100 IRMS. Final isotopic ratios are reported as delta values per mil (‰) relative to an international standard (V-PDB for carbon and oxygen), calibrated with international standards (IAEA-603: *d*^13^C =+2.46 ‰ / *d*^18^O = −2.37 ‰ and NBS-18: *d*^13^C = −5.014 ‰ / *d*^18^O: −23.2 ‰), as well as three in-house standards. Multi-point standard isotope calibration was carried out using the IonOS software (Version 3.2) by Elementar by generating a trend line (y = mx+c) that maps measured vs. expected isotopic results, which is then used to calibrate sample results.

The measurement uncertainty was monitored using three in-house standards. The overall analytical precision is higher than 0.1‰ for *d*^13^C values and better than 0.2‰ for *d*^18^O values. The conversion of *d*^18^O values towards the V-SMOW standard has been done using the formula: *d*^18^O_V-SMOW_ = *d*^18^O_V-PDB_.1.03096 – 30.86.

The strontium isotopic analysis was performed on pretreated enamel powders at the clean laboratory facilities of the Curt-Engelhorn Centre for Archaeometry at Mannheim, Germany, following a well-established protocol^79^. The determination of ^87^Sr/^86^Sr ratios was performed by a high-resolution multi collector ICP-MS (HR-MC-ICP MS, Neptune). Raw data were corrected according to the exponential mass fractionation law to ^88^Sr/^86^Sr = 8.375209. Blank values were lower than 50 pg Sr, corresponding to less than 0.1% of the Sr-content of the analyzed sample.

***Collagen isotopic analysis***

A very small fragment (~100 mg) of Neanderthal tooth dentine was treated following a well-established protocol^80^ at the University of Tübingen. The sample was ultrasonicated in acetone and then rinsed with distilled water. Once dried, it was powdered and sieved to a particle size of less than 0.7 mm. For collagen extraction, bone powder was decalcified in 1 M HCl for 20 min at room temperature and filtered through a 5 mm filter. The insoluble residue was then soaked in 0.125 M NaOH for 20 h at room temperature. Subsequently, the rinsed residue was heated in closed tubes at 100^◦^C for 17 h in HCl pH2 solution, in order to gelatinize the collagen. After filtration through a 5 mm filter, the filtrate containing gelatinized collagen was freeze-dried.

The stable carbon and nitrogen isotopic compositions were performed in duplicate at the Institute of Environmental Science and Technology (ICTA, Barcelona, Spain) using a Thermo Flash 1112 (Thermo Scientific VC) elemental analyzer coupled to a Thermo Delta V Advantage mass spectrometer with a Conflo III interface. This measures the ratios of ^13^C/^12^C and ^15^N/^14^N relative to an international standard (V-PDB for carbon and AIR for nitrogen). The international laboratory standard, IAEA 600 (caffeine), was used. Analytical uncertainty was determined to be ±0.20 ‰ for both *δ*^13^C and *δ*^15^N, based on multiple measurements of collagen extracted from modern bones of camel (*Camelus dromedarius*) and elk (*Alces alces*).

***Trophic position of Mandrin Neanderthal***

*Collagen carbon and nitrogen isotopes and trophic position of the Mandrin Neanderthal*

The dentine fragment from the Mandrin Neanderthal yielded good quality collagen, with C/N ratio of 3.4 and carbon and nitrogen content of 34.7% and 12.0%, respectively, well in the range of well-preserved collagen^81,82^. The *δ*^13^C and *δ*^15^N values of -19.5‰ and +11.7‰, respectively, are within the range of those obtained for late Neanderthals from Belgium^83^ and western France^84^. A direct comparison is however difficult, since the specimen from Mandrin comes from a tooth while the measurements of the other specimens were made on bone, as the previously published data, and this could lead to slightly higher *δ*^15^N values. Another possible interfering factor is the change of baseline that has been documented in other regions of France around the time of Mandrin Neanderthal^85^. To investigate this possibility, we gathered isotopic data from other sites from the same Southeastern France region at around the same time and compared these values to those of Southwestern France around the Neanderthal site of Saint-Césaire (Fig. S3). Despite slightly lower *δ*^15^N values in Southeastern France compared to those of Southwestern France, the isotopic values of Thorin are plotting in the same position compared to the herbivorous species as in Saint-Césaire, indicating that Thorin had a similar trophic position in its ecosystem as the late Neanderthal from Saint-Césaire. A more detailed paleodietary reconstruction will be performed when collagen isotopic data will be obtained from the ungulate fauna from Grotte Mandrin, but there is no indication that the Neanderthal from Grotte Mandrin could not be part of the ecosystem represented by the faunal remains in the region.

*Enamel carbon and oxygen isotopes and environmental context of the Mandrin Neanderthal*

The *d*^13^C values of Mandrin Neanderthal are about 3-4‰ lower than those of the coeval ungulates and in the same range as those of Neanderthals from Payre^71^. Such a difference in *d*^13^C values between coeval hominids and ungulates is due to the differences in fractionation between diet and enamel carbonates between both trophic groups and is typically found for Pleistocene hominins in Europe^71,86^. Since the *d*^18^O values of Mandrin Neanderthal are similar to those of coeval fauna, and much lower than those of Holocene humans living under interglacial conditions from Mandrin and Payre, the combined *d*^13^C and *d*^18^O values of this hominin are fully consistent with a life in the same environmental conditions as the ungulates from the MIS3 layers where it was found.

*Strontium isotopes and mobility of the Mandrin Neanderthal*

The ^87^Sr/^86^Sr ratios measured on the Neanderthal tooth enamel (0.70882±0.0005) were very similar to those of coeval horses and bison (0.70803±0.000003 to 0.70863±0.00002) and a human Neolithic tooth from Grotte Mandrin (0.70803±0.00003). These values are close to those measured on Neanderthal tooth enamel from Payre, a few dozen kilometers north of Grotte Mandrin along the Rhône Valley (070909±0.00003 to 0.71080±0.00005; Fig. S3)^71^. When compared to the range of variation for ^87^Sr/^86^Sr ratios in Southeastern France, as reported in Fig. 7 of ref. 71, such values are broadly spread so it is not possible to establish an exact location of the Mandrin Neanderthal at the time of its tooth formation, but nothing indicates that this individual would have spent some of this youth far away from the location of Grotte Mandrin.

**Sample processing and DNA sequencing**

*DNA extractions*

Sample processing was performed in a dedicated clean laboratory facility at the Lundbeck Foundation Centre for GeoGenetics, GLOBE Institute, University of Copenhagen, Denmark, following strict ancient DNA procedures. Each extraction, and subsequent USER^®^ treatment, library building, and PCR setup session included a mock reaction where no sample (for extraction) or QIAGEN^©^ EB buffer (for the following steps) was added to the reagents instead of DNA solution. These pre-PCR steps were performed in the ancient DNA facility, physically separated from laboratories where post-PCR and fresh modern samples are processed.

The surface of a first molar root (sample MAN-15-B2-1600, also registered under the GeoGenetics accession number CGG_2_016357) was cleaned by gentle abrasion, and 280mg of the root tip was manually crushed using a mortar. DNA was extracted following a silica-column based method described in ref. 87 and adapted by ref. 88 (method Y). Root fragments were pre-digested for 2h at 37ºC by incubating in 1mL of an extraction buffer (0.45M EDTA, 0.25mg/mL Proteinase K and 0.5% N-LaurylSarcosyl). After centrifugation for 2 minutes at 13,000 rpm, the supernatant was removed and the remaining pellet was subjected to three sequential digestions as described below.

After 24h of incubation at 37ºC in fresh digestion buffer, the microtube was centrifugated for 2 minutes at 13,000 rpm and the supernatant E1 (labeled as BE1 for the extraction blank) was recovered without disturbing the remaining undigested pellet. This pellet was further incubated with 1mL of fresh digestion buffer for an additional 42 h at 37ºC, centrifugated for 2 minutes at 13,000 rpm and the supernatant E2 (BE2 for the extraction blank) was recovered. As there was still undigested material, a final incubation of 72h at 37ºC in fresh digestion buffer was performed, and supernatant E3 (the extraction blank being labeled as BE3) recovered after centrifugation for 2 minutes at 13,000 rpm. For any incubation lasting for more than 24h, 0.25mg/mL of fresh Proteinase K was added after each 24 h of incubation.

Each of the supernatants E1, BE1, E2, BE2, E3 and BE3 was concentrated down to 200μL volume using an Amicon Ultra-4 30kD device (Merck Millipore). DNA solutions were further purified on a MinElute column (QIAGEN^©^) and eluted in 60μL elution buffer (QIAGEN^©^ EB supplemented with 0,05% Tween 20) after 15 minutes incubation at 37ºC. All extracts were stored at -20ºC in siliconized tubes (Eppendorf® LoBind).

*USER treatment, sequencing library building and indexing*

A fraction of 14.9μL of each extract was directly built into Illumina sequencing libraries to validate the presence of post-mortem DNA damage signatures. Another fraction (32.5μL) of each extract was incubated for 3h at 37ºC with 10μL Uracil-Specific Excison Reagent (USER^®^, NEB^TM^ reference M5505) enzyme mix in order to remove uracil residues and limit the impact of nucleotide mis-incorporations in the analysis^89^.

On each extract, multiple independent library constructions and amplifications aim at limiting PCR duplicates and thus reducing sequencing costs (Table S4). Blunt-End Illumina sequencing libraries were built on both USER-treated and non-USER-treated extracts, following ref. 90 with slight modifications as described in ref. 91, using the NEBNext^®^ DNA Library Prep Master Mix Set (NEB^TM^ reference E6070) without the ssDNA isolation module.

To determine the optimal number of Polymerase Chain Reaction (PCR) cycles to be used for amplifying libraries, an aliquot of 1μL of each library was diluted 20 times and subjected to a real-time PCR assay, as recommended in ref. 90. Subsequently, a volume of 12μL (out of the 25μL total library volume) of each unpurified library was amplified in a PCR reaction volume of 50μL, using the KAPA HiFi HotStart Uracil+ ReadyMix (KAPA Biosystems, Inc), 200nM of PCR primer IS4^90^ and 200nM of a custom-designed primer containing a 6-bp known index sequence used for post-sequencing demultiplexing. Thermocycling conditions were as follows: 1 min at 94ºC, followed by N cycles of 15 sec at 94ºC, 20 sec at 60ºC, and 20 sec at 72ºC, and a final elongation step of 1 min at 72ºC. The cycle numbers N used for each library amplification are summarized in Table S4. When less than 12 cycles were necessary, libraries were subjected to a single PCR amplification round, purified on a MinElute column (QIAGEN^©^) and eluted in 20μL EB with 0.05% Tween.

Otherwise, PCR amplification was carried out in two rounds: after the first 10-12 cycles amplification round, PCR products were purified on a MinElute column, eluted in 20μL EB (QIAGEN^©^) and split over four PCR reactions. The four final PCR products were then pooled and purified on a MinElute column and eluted in 10μL EB with 0.05% Tween. A 1/10 dilution of each purified product was quantified on an Agilent 2200 TapeStation instrument (Agilent Technologies).

*Whole Genome Enrichment*

To increase the human DNA fraction in the sequencing libraries, we performed an in-solution Whole Genome Enrichment (WGE), using MYBaits RNA probes synthesized at MYcroarray (Ann Arbor, MI, USA^92^). Following manufacturer’s instructions (MYbaits manual v2.2), 70 to 650ng of amplified libraries were incubated for 40h at 55ºC with 5μL of the synthetic probes (one capture reaction per library), in presence of blockers (Human Cot-1 DNA, Salmon Sperm DNA and Proprietary Blocking Agent). After purification using Dynabeads™ MyOne™ Streptavidin C1 beads (Invitrogen), captured libraries were eluted in 30μL EB with 0.05% Tween. Real-time PCR was performed on a 1/20 dilution of the eluate, to determine the minimal number of PCR cycles required to amplify the captured libraries up to a molarity compatible with the Illumina sequencing technology. The whole volume of WGE-enriched libraries was then amplified in a 100μL final PCR reaction, with 2μL of AmpliTaq Gold DNA Polymerase (Applied Biosystems) and 200nM of each primer IS5_reamp.P5 and IS6_reamp.P7^90^, for 7 to 13 cycles (Table S4). Thermocycling conditions consisted of initial denaturation for 5 min at 94ºC, followed by N cycles of 30 sec at 94ºC, 30 sec at 60ºC and 40 sec at 72ºC, and lastly an elongation step of 7 min at 72ºC. Amplification products were purified on MinElute column and eluted in 20μL EB + 0.05% Tween.

*Sequencing*

Purified libraries were quantified on an Agilent 2200 TapeStation instrument (Agilent Technologies) and pooled, requiring at least three base differences between any indices pair. Pools were quantified using a Kapa Library Quantification Kit Illumina (KapaBiosystems, KK4854) real-time PCR assay and sequenced at the Danish National High-Throughput DNA Sequencing Centre, Copenhagen University, Denmark, on an Illumina HiSeq2500 platform, with 100SR, 100PE or 80PE mode, with single i7 index read. Basecalling and demultiplexing were performed using CASAVA v.1.8.2.

**Read processing and mapping**

Illumina reads were processed for each individual library using the PALEOMIX v1.1.1 pipeline^93^ with default parameters, except that minimal mapping quality threshold was set to 30 and seeding was disabled. Briefly, sequencing reads were trimmed for known adapter sequences, low quality termini, and filtered out if shorter than 30 nucleotides using AdapterRemoval^94^ . Paired-end reads overlapping for 11 nucleotides or more, with a maximal edit distance of 1, were collapsed and further treated as single-reads. Trimmed and collapsed reads were aligned to the human reference genome hg19 available from the UCSC genome browser (<http://hgdownload.cse.ucsc.edu/goldenPath/hg19/chromosomes/>) using BWA version 0.5.9-r26-dev^95^, collapsed for duplicates and re-aligned locally around indels using GATK^96^. Uncollapsed paired-end reads most likely correspond to long contaminating DNA templates of modern origin and were thus filtered out of the final BAM file.

The whole procedure was repeated against the revised Cambridge reference mitochondrial sequence (rCRS, Accession Number NC_01920) to generate mitochondrial read alignments. Summary statistics obtained from PALEOMIX are presented in Table S5.

**Data authentication and contamination estimation**

Presence of ancient DNA was confirmed using mapDamage2.0^97^ by detecting excess C to T substitutions in the strand termini of each generated library. aRchaic^98^ was used to detect additional types of damage patterns by applying a grade membership model, which clusters the tested samples into groups based on their mismatch profiles. We, furthermore, estimated the level of sequencing errors in the libraries following the approach in ref. 99. The method is based on measuring an excess amount of derived alleles relative to a high quality genome, which is assumed to be error-free, using ANGSD^100^. Mitochondrial contamination estimates were carried out using contaMix^101^ by first generating a consensus sequence using BCFtools *v.1.10.2* (<https://github.com/samtools/bcftools>) for each library and then aligning them to 311 present-day modern humans with mafft^102^. X chromosome contamination estimates were carried out using ANGSD by generating a binary count file covering sites on the X chromosome with a base quality of at least 20 (-r X: –doCounts 1 –iCounts 1 –minQ 20). The count file was used as input along with a HapMap file of CEY (Europeans) to the contamination estimator, from which maximum likelihood estimates of contamination levels were obtained.

**Mitochondrial DNA (mtDNA) and Y chromosome analysis**

*Metagenomic classification of reads from sediment samples*

Sediment samples^14^ included in the mtDNA analysis were processed through the metagenomic classifier *KrakenUniq v. 0.5.8*^103^. We filtered the raw alignments (obtained from European Nucleotide Archive, accession PRJEB42656) to only include reads with at least 35 base pairs and mapping quality of at least 25 and performed metagenomic classification using a custom database of all full mitochondrial and plastid genomes in RefSeq. All reads classified within the genus of primates were used for further analysis. Variant calling was carried out with ‘*bcftools call’* (-m --ploidy 1) and filtered for sites with read depth of ≥ 5 and genotype quality of ≥ 25. To ensure that only confidently called sites were used for analysis we excluded sites where the genotype was supported with less than 75% of the reads. Excluded sites were set to missing in the final consensus calling generated by using ‘*bcftools consensus’*.

*mtDNA analysis*

We carried out a maximum likelihood (ML) analysis to infer a tree topology based on the mitochondrial genomes of Thorin, 43 ancient humans^104-106^, 34 Neanderthals^8,11,12,16,33,44,46,49,107-112^, and 4 Denisovans^113-116^, using RaxML-NG^117^. The tree was inferred using 100 bootstraps under the GTR-I-G4 substitution model (options: --all --bs-trees 100). We furthermore performed a maximum parsimony analysis using Denisova 3 and 23 Neanderthals (included in the ML analysis) only considering sites within the coding region (rCRS coordinates: 577-16,023). Using the pratchet function from the R package phangorn^118^, a distance matrix was computed including raw substitution rates, which was used to perform a branch-shortening analysis^8^. Relative age estimates were obtained using Goyet Q56-1, which has been radiocarbon dated to be 42,540 years old^16^, as a reference sample. The final age estimated were computed using the substitution rate of the coding region in modern humans, 1.57×10^-8^ substitutions/bp/year (95% confidence interval (CI): 1.17×10^-8^-1.98×10^-8^ substitutions/bp/year^118^). BEAST v. 2.6.3^45^ was used to perform molecular tip dating on Thorin and to estimate divergence dates. The analysis included 24 Neanderthals^11,16,33,110^, 1 Denisovan^120^, 9 ancient human genomes^105,106^, and 53 contemporary human mitochondrial genomes^121^. We only considered sites within the coding region. For the initial runs, we performed two sets of analysis, one with a fixed substitution rate of 1.57×10^-8^ substitutions/bp/year and another with an estimated substitution rate. Previously radiocarbon- and molecularly dated samples were used as calibration points, where the point estimates were used as initial values and their corresponding 95 Confidence Intervals (CI) were used as a uniform prior for the respective samples. For undated Neanderthals, an initial value of 50,000 years ago along with a uniform prior ranging from 30,000-300,000 years ago. A uniform prior to the TMRCA of all Neanderthals was set to 100,000-infinity, while a uniform prior for the TMRCA of modern humans was set to 50,000-infinity. A substitution model was obtained using bModelTest^122^, while the best fitting tree/clock model was found using stepping stone analysis from the BEAST2 model-selection package.

*Y chromosome analysis*

Phylogenetic analyses of the Thorin Y chromosome sequence were carried out using a reference panel of shotgun sequencing data for 21 human samples (19 high coverage ancient samples; 2 modern samples with A00 haplogroup) and five archaic humans (three Neanderthals and two Denisovans; Table S6). Individual sample genotypes were obtained by selecting the majority allele at each genomic site covered by a minimum of two sequencing reads, with the further requirement that >80% of reads show the majority allele. A multisample genotype matrix was then obtained by identifying biallelic SNPs across all samples, restricting to the ~6.9 Mb long “accessible” Y chromosome regions targeted by the capture probes in ref. 123. Analyses including the El Sidrón 1253 sample were further restricted to the reduced ~560 kb region captured in that individual. Phylogenetic trees were constructed by applying the neighbor joining algorithm on the pairwise mismatch distance matrix, using the ape R package, with the alleles observed in the Chimpanzee genome as outgroup.

**Analysis panel**

A panel of 30 contemporary individuals^137^, 18 high quality ancient genomes^9,88,104,124-130^, and 14 archaic genomes^8,11-13,15,16,46,49,114,131^ (Table S7). Low coverage Neanderthals were filtered with SAMtools v. 1.3.1^132^ to discard reads with a mapping quality below 25. We kept two versions of each low coverage sample, one BAM file including all reads, while the second only contained deaminated reads. Both sets were obtained using pseudo haploid genotyping by sampling a random allele on each site.

We carried out a two-step approach to combine the data of the high and low coverage Neanderthals. We first selected all biallelic SNPs from previously published diploid genotype calls for the four high-coverage Neanderthals, and extracted pseudo-haploid genotypes from the low coverage data subset at the same sites. Genotypes in the low-coverage samples that did not match either of the two alleles observed in the high-coverage samples were set to missing. We then extracted biallelic SNPs segregation among the low-coverage samples only, and merged both sets. For the final analysis panel, we including only biallelic transversion SNPs, as well as restricting to regions within the 1000 Genomes Phase 3 strict accessible genome mask (ftp://ftp.1000genomes.ebi.ac.uk/vol1/ftp/release/20130502/supporting/accessible_genome_ mask s/20141020.strict_mask.whole_genome.bed).

**Population genetic analyses**

*Procrustes analysis*

The data was filtered with PLINK v. 1.9^133^ to keep sites with minor allele frequencies above 0.05 and no missing genotypes. Procrustes analysis was then carried out by first performing a principal component analysis (PCA) with EIGENSOFT^134^ using the Altai Neanderthal, Vindija 33.19, and Denisova 3. Subsequently, we obtained raw PC coordinates for each low-coverage Neanderthal sample by individually projecting them onto the three reference samples. The obtained raw PC coordinates were then transformed to align with the reference PCA coordinates by using the *procrustes* function from the R package *vegan*^135^.

*D-statistics*

D-statistics was carried out using ADMIXTOOLS 2.0 (<https://uqrmaie1.github.io/admixtools/>) in R. All estimates were obtained using a block jackknife with a block size of 5 centiMorgan.

*Treemix*

Treemix^136^ was carried out by using two samples from Mbuti (sample IDs: HGDP00456, SS6004471)^137^ as root population (Fig. S19). For the low coverage Neanderthals, we used deaminated reads from Mezmaiskaya 1 and 2, while we used all obtained reads passing the initial filtering steps from Thorin, Goyet Q56-1, and Les Cottés Z4-1514. The analysis by accounting for linkage disequilibrium (-k 300) and without sample size correction (-noss).

*Runs of homozygosity*

We called runs of homozygosity (ROH) using a novel approach employing spatial smoothing of read allele frequencies along individual Neanderthal genomes. We first identified a set of bi-allelic reference SNPs polymorphic across the four high coverage archaic genomes (Altai, Chagyrskaya 8, Vindija 33.19, Denisova) and one human genome (HGDP01029, San). For each target Neanderthal genome, we then tabulated the frequencies of the two alleles for each reference SNP covered by ≥ 2 reads in the target genome. The obtained vector of minor read allele frequencies along each chromosome was smoothed by fused lasso regression^138^, using the function ‘fusedlasso1d’ implemented in the *genlasso*^139^ R package (version 1.6.1). To call ROHs, we collapsed consecutive SNPs with estimated coefficients β<0.005 into segments. We evaluated the performance of this approach in low coverage settings by randomly subsampling the high coverage Neanderthal genomes to lower coverage (1X, 2X).

*Demographic modeling*

We used the site-frequency spectrum (SFS)-based method *momi2*^50^ to carry out demographic modeling from allele count data. Modeling was carried out using an SFS with ascertainment restricted to the four high-quality archaic samples with diploid genotypes (Altai Neanderthal, Chagyrskaya 8, Vindija 33.19, and Denisovan). To account for the exclusion of transition SNPs in the dataset, we used a transversion-only mutation rate of 0.16e-9 / bp / year as estimated by Fu et al. The generation time interval was assumed to be 29 years. Each demographic model was fit using ten independent replicates from randomized parameter starting values, to evaluate consistency of parameter estimates at convergence. We used the “TNC” method for model optimization, with a maximum number of 400 iterations. Confidence intervals for parameter estimates were obtained using 100 parametric bootstrap replicates, as implemented in the *momi2* package. To further investigate the model fit, we used the *f*_4_* statistic, calculated as log(BABA) - log(ABBA) as previously described. We compared distributions of Z-scores comparing *f*_4_* statistics expected under the best-fit model with those observed from the real data.

**Supplementary Note 1 - Site and fossil descriptions**

Grotte Mandrin is a vaulted rock shelter directly overlooking the middle valley of the Rhône River, one of the largest and most important rivers of Western Europe. The Rhône reaches the current shore of the Mediterranean coastline some 120 km south of the site (Fig. 1). Research conducted since 1990 has allowed the excavation of most of the floor surface from under the vaulted area of the cave. Mandrin provides a reference archaeological succession for it contains all of the phases currently known from the end of the Middle Palaeolithic (Levels F to B2) up to the early Protoaurignacian (Level B1), a culture marking the advent of the full Upper Palaeolithic in Western Europe at around 42 ka^1^.

Each archaeological unit has yielded a rich lithic industry (~60,000 lithics total) associated with numerous paleontological materials (~70,000 fauna). Faunal remains are almost exclusively related to human activities in the cave, as shown by their high frequency of anthropogenic modifications. Geoarchaeological study has shown that the overall preservation of the stratigraphic units is good (Levels B to D) to even excellent (Levels E-F). This Pleistocene sequence is covered by recent Holocene sediments (Level A) recording different phases of human cremations from the late Neolithic^1^. While Levels B1 to F are known from a large surface of excavation (ranging from 100 to 50 m²) and concerning most of the area of the cave, the lower part of the sequence, Levels G to J, is only known from ~5m² from different test-pits at the site. The lithic industries from these oldest layers are also rich and allow a good documentation of these technologies based on a total of 7,687 lithics. These layers are attributed to the Marine oxygen-Isotope Stage 5 (MIS 5) based on initial optically stimulated luminescence (OSL) dates and warm-adapted faunal communities.

The bulk of the Pleistocene deposits were formed by sedimentation of windblown local sands and silts entering the cave. Carnivore activity in the shelter is limited to a few traces of wolf and fox activity represented by rare coprolites and gnawing marks on bones initially abandoned by hominins. Levels B to D have yielded several anthropogenic combustion areas.

All archaeological units are separated by thin sterile layers free of any archaeological material. A total of 59,338 lithics were recovered from all the archaeological levels. The lithics are well preserved. The good preservation of the thin edges of the lithic artifacts show rare or no taphonomic movement, in agreement with their rapid covering by aeolian deposits. In the upper part of the Pleistocene sequence eight archaeological units (B1 to F) have been unearthed and divided into five cultural phases (from base to top): 1) Level F, Rhodanian Quina Mousterian, 2) Level E, Neronian^1^, 3) Level D, Post-Neronian I Mousterian, 4) Levels C2, C1, B3, and B2, Post-Neronian II Mousterian, and 5) Level B1, Protoaurignacian.

These 8 stratified levels encompass ~15 millennia and provided most of the lithic collections of Mandrin with 51,650 artifacts including 13,440 that are ≥2 cm in maximum size.

Thorin was found in the Post-Neronian II units associated with a very rich archeological material. Thorin has been progressively uncovered since 2015 and appears as a very fragmented but potentially fairly complete individual, recovered on a very small surface of about 50 cm² (Figs. 1, 2, & S1).

The post-Neronian II Mousterian, associated with Thorin, is found in Levels C2, C1, B3 and B2, which provided 7,853 lithics ≥2cm. Level C2 provided 3,410; Level C1 provided 1,567; Level B3 provided 1,653; and Level B2 provided 1,223 lithics ≥2cm. These four Levels, well distinguished by spatial distribution analyses and by their chronologies (52.9 – 44.1 ka cal BP) (*1*) encompassing ~8-9 ka, are attributed to the Post-Neronian II Mousterian culture. These 4 Mousterian levels are technically identical to one another, based on the production of flakes mainly obtained by Discoid flaking. These flakes are retouched in a large variety of Mousterian scrapers. Altogether, a total of 497 Mousterian scrapers were found dominated by lateral convex scrapers and transversal scrapers. None of these 497 scrapers present a Quina retouch, categories of retouches that concern more than a quarter of the scrapers in Level F. Some rare large points are attested in these Post-Neronian II levels. Some Levallois flakes are produced, obtained mainly by recurrent unipolar and preferential methods.

*Thorin’s hand skeleton*

Five manual skeletal elements were collected during the 2016 and 2017 excavation campaigns, comprising 3 distal phalanges and 2 intermediate phalanges. These specimens, found in close proximity to each other, are not redundant, and when they show elements of lateralization, they are all located on the left side. They show a mature stage of development.

*Distal phalanges*

There are 3 distal phalanges, numbered 1261, 1586 and 1579 (Fig. S2). They were respectively attributed to the distal phalanges of the first, second and third fingers of the left hand.

*Specimen 1261: distal phalanx of the left thumb (DP1)*

The specimen is incomplete, the base and a proximal part of the shaft are missing, with the bone showing a taphonomic V-shaped fracture with a proximal point. The distal tuberosity (apical tuft) is large (medial-lateral width greater than 9.5 mm, a minimum value due to taphonomic erosion of the lateral part); it is widely implanted on the diaphysis, rounded in shape, with a slight asymmetry projecting its apex to the medial (ulnar) side. These morphometric characteristics bring this distal phalanx closer to those preserved for Neanderthals. In contrast, the distal phalanx of the modern human thumb is more slender, with a distal tuberosity that is less wide, relative to its length, and is lanceolate and symmetrical in shape^22,140-142^ . Despite its fragmentary nature, the morphology of this 1261 specimen is thus clearly Neanderthal.

*Specimen 1586: distal phalanx of the left index finger (DP2)*

This is an upper end of a distal phalanx, with the entire distal tuberosity and two-thirds of the shaft, a taphonomic fracture running obliquely from bottom to top and from the lateral to the medial edge has separated the base of the phalanx which is missing. The distal tuberosity is complete, rounded in shape, with a medio-lateral width of 8.5 mm. This value is within the biometric variability of the distal phalanges of the Neanderthal index (from 8.5 to 10.5 mm) and is greater than that of AMH Qafzeh 9 (6.8 mm) and the fossils of the Early Upper Paleolithic (6.6 to 7.4 mm)^140^. While this value of 8.5 mm is higher than the average value for contemporary Europeans (7.9 ± 1 mm), it falls within the upper range of their variability. However, the very rounded morphology of this distal tuberosity present on this piece 1586 is observed in Neanderthals and differs from that of modern humans, for whom the phalangeal tuft is lanceolate^140,142^.

*Specimen 1579: distal phalanx of the left middle finger (DP3)*

This specimen is complete. Although the distal tuberosity is very slightly eroded around the edge, its shape is clearly rounded and not lanceolate. Its length is 19 mm (minimum value) and is above the average for contemporary males (18.3 ± 1.8 mm)^22^ while falling within the upper range of their variation. This value is slightly lower than that of the Neanderthals of Ferrassie 1 (21.9 mm), Regourdou 1 (21.4 mm), Shanidar 4 (21.5 mm) and 5 (21.2 mm) and Kebara 2 (21.4 mm)^143^ and is closer to that of Kiik Koba EM179 (20.3 mm)^22^. Furthermore, the value of this measurement is higher than that of the specimens from La Ferrassie 2 (17.4 mm) and Shanidar 3 (17.6 mm)^23^. The width of the distal tuberosity is 9.5 mm (minimum value due to taphonomy), reflecting a phalangeal tuft width greater than that of Qafzeh 9 (8 mm) and the average of contemporary males (8.7 ± 1.2 mm) but within the upper range of their variation. This value measured here is lower than that measured for other Neanderthals such as La Ferrassie I (12.2 mm), Kiik Koba I (13.8 mm), Shanidar 4 (12.2 mm), Shanidar 5 (11 mm) but remains close to those of Shanidar 3 (10.8 mm) and Regourdou 1 (10 mm)^23^.

*Proximal and intermediate phalanges*

They are represented by two fragmentary pieces, numbered 1262 and 1587 (Fig. S2), identified respectively as the proximal phalanx of the thumb and the intermediate phalanx of the index finger, the fragmentation of which does not provide any taxonomic information, unlike the distal phalanges, which have autapomorphies.

*Specimen 1262: proximal phalanx of the thumb*

This corresponds to the distal half of the phalanx, a taphonomic fracture having occurred mid-diaphysis. The distal articular surface is fairly well preserved, with a medio-lateral width of 9 mm.

*Specimen 1587: middle phalanx of the index finger*

It consists of two unrelated fragments, one distal and one proximal, each with its corresponding articular surface. Their morphology allows them to be attributed to the second finger, but this attribution must be confirmed by the study of the new elements of the hand skeleton, recently discovered

Distal phalanges of the left hand of Thorin (DP 1, DP2, DP3) display clear Neanderthal features. The distal pollical phalanx presents the medial (ulnar) deviation of the distal end described in Neanderthal specimens^23^. All three phalanges show a rounded morphology of the distal tuberosity that differs markedly from that of modern humans. This morphology is described as specific to Neanderthals^23,142^.

**Supplementary Note 2 - Quality control and contamination estimates**

**Authentication of ancient DNA fragments**

We restricted the analysis to reads with a mapping quality of at least 30 and only reads covering the highly mappable regions of the genome^144^. To confirm the presence of ancient DNA in the libraries, the C to T and G to A damage patterns in the strand termini were quantified along the fragment lengths using the software *mapDamage2.0*^97^. This was carried out by measuring the mismatch rates along the fragments and subsequently plotting them as a function of the position. *mapDamage2.0* statistically models the post-mortem patterns in a Bayesian framework and returns the posterior distributions of the parameters, which are related specifically to the cytosine deamination. Our non-USER-treated libraries show signs of deamination in the strand termini confirming the presence of ancient DNA fragments in the tested libraries. However, we find that the non-USER treated library from extract 1 (Thorin_1600_E1L1P2) has significantly less fragments with deamination patterns compared to the other non-USER treated libraries (Fig. S2, Table S8).

We further studied the damage patterns in the libraries using *aRchaic*^98^, which uses a grade membership model in order to detect different types of damage patterns. In contrast to mapDamage, *aRchaic* not only looks for signs of cytosine deaminations, but detects a range of non-predefined damage patterns. Therefore, BAM files from all libraries, regardless of whether they have been USER-treated or not, are relevant as input. *aRchaic* returns Mismatch Feature Format (MFF) files for each of the tested libraries containing the information of mismatch types, flanking bases, strand break bases and positions of the observed mismatches. Based on the detected features, the mismatches are clustered into groups called mismatch profiles. The number of profiles used for the analysis are defined by *K*. The grade of membership is determined by the relative frequency of each mismatch profile observed in the individual samples. The implemented model assumes each of the mismatches observed in the samples belong to one of the *K* mismatch profiles.

All BAM files were down-sampled to 10,000,000 reads. We used data from 22 libraries of the Thorin sample along with 33 archaic individuals^8,11,12,15,16,49,114^, 10 ancient samples^9,104,124-131^, and 39 modern human individuals^137^ for reference. The results of *K* = 2 display the difference in the level of cytosine deamination patterns between modern, non-USER treated ancient, and USER-treated ancient libraries. Both the modern samples and USER-treated libraries show a high membership in the first cluster due to the removal of deaminated cytosines, which causes the ancient fragments to mimic modern DNA. Although modern DNA is not expected to show a significant amount of damage, we find an elevated level of C to A and T to C in the 5’ end along with reduced levels of T to A and T to G substitutions. The non-USER treated libraries show a high membership in the second cluster, where the driving damage pattern is the C to T substitution in the strand termini (Fig. S5). Results from *K* = 4 show the modern samples primarily assigned to cluster 1, while the libraries generated on the Neanderthal fossils presented in ref. 16 are mainly assigned to cluster 2. The non-USER treated ancient libraries are displaying a high level of membership in cluster 3 and USER-treated libraries are mainly assigned to cluster 4 (Fig. S6). Thus, we can distinguish USER-treated ancient libraries from libraries containing modern DNA, meaning that a significant membership in cluster 1 for the ancient libraries is an indication of contamination of modern DNA. Considering this, we observe that extract 1 of the Thorin sample contains relatively high amounts of modern contamination compared to extracts 2 and 3.

**Estimation of sequencing error**

We also estimated the rate of sequencing error of our libraries. Similar to the approach used in ref. 98 both an overall as well as a type specific error estimation were performed using ANGSD^100^. The analysis is based on the observed number of excess derived alleles in the data relative to a high-quality individual, which we assume to be error free. In the case of a sample with no sequencing error, the expectation is to observe an equal number of derived alleles in the data and the high-quality individual. As our high quality sample, we used present-day Mbuti individuals from Simons Genome Diversity Panel^137^. In order to define the ancestral allele states of each polymorphism, the method requires an outgroup, for which we used a chimp genome. Only the intersection of sites covered by the test sample, the error free individual and the outgroup were considered by the method.

The results of the overall error rates (Fig. S7) show that the non-USER treated libraries show a high error rate compared to the USER-treated libraries. However, we find relatively lower error rates in the non-treated libraries from extract 1 (Thorin_E1 and Thorin_E1L1P2), which could be explained by modern contamination reducing the fraction of ancient DNA fragments in the libraries. Taking a closer look at the type specific error rates, we observe that the most dominant types of substitutions are C to T and G to A, which is expected in non-treated libraries containing ancient DNA (Fig. S8).

**Mitochondrial contamination estimates**

We first carried out contamination estimations based on the mitochondrial (mt) DNA using *ContaMix*^101^. We used *BCFtools v.1.10.2* (<https://github.com/samtools/bcftools>) to generate a consensus sequence for each library. Each of the consensus sequences were aligned to 311 present-day human mtDNA sequences with *mafft*^102^. We, additionally, mapped the BAM files to their respective consensus sequences with the *BWA* package^95^. We discarded unmapped reads, reads below mapping quality 30, and reads with alternative hits using *SAMtools*^133^, so that only uniquely mapped reads were kept for further analysis. Both the mt alignment as well as the remapped BAM file were used as input to *ContaMix*.

The resulting point estimates (Table S9) of the mtDNA contamination indicate that libraries built on extract 1 are significantly contaminated compared to extracts 2 and 3. Additionally, we ran the analysis on a merged BAM file including data from all libraries with a resulting point estimate of authentic reads at 0.77. Excluding extract 1 decreases the level of contamination significantly to a proportion authentic of 0.997.

**X chromosome contamination**

Given that Thorin is a male individual, we further assessed the level of nuclear contamination using the X chromosomal reads. We carried out the analysis following the procedure described in ref. 145, which utilizes the contamination estimator provided by ANGSD. Based on the idea that males only have one allele per site on the X chromosome, different alleles observed at one site are assumed to be an indicator of either sequencing error or contamination. The true allele is defined by the most frequently observed allele at each site requiring that the data have a decent coverage.

We generated a binary count file only covering the X chromosome including reads with base quality of at least 20 (-r X: –doCounts 1 –iCounts 1 –minQ 20). The count files were then used as input to the contamination estimator along with a HapMap file of CEU (European) individuals. We find that extract 1 is significantly more contaminated than extracts 2 and 3, which influences the overall level of contamination in the merged BAM file containing the reads of all libraries. Due to this issue, we assessed the level of nuclear contamination in the BAM file including all reads excluding extract 1, from which we obtain a reduced amount of contamination at ~0.01 (Table S10).

Due to the consistent observations of extract 1 being more contaminated relative to extracts 2 and 3, we chose to exclude data from these libraries in all subsequent analyses.

**Supplementary Note 3 - Uniparental markers: Mitochondrial and Y-chromosome analysis**

We studied the phylogenetic relationship between Thorin and previously published Neanderthals using a ML approach based on the mtDNA. We included 43 ancient humans^104-106^, 34 Neanderthals^8,11,12,16,33,43,46,49,107-112^, and 4 Denisovans^113-116^. We carried out the genotype calling for Thorin using ‘*bcftools* *call’* (-m --ploidy 1) and only considered sites covered by at least 5 reads with a genotype quality of at least 30. We generated the consensus sequence with ‘*bcftools consensus’*. Sites not passing the filtering were set to missing in the final sequence. In order to identify possible ambiguous genotypes in the consensus sequence we computed the frequency of the alternate allele across all sites along the mitochondrial genome. We found only two possible ambiguous sites where both the reference and alternative alleles were observed at intermediate frequency (between 10-90 %; Fig. S9). At both sites the minor allele frequency was ~30%, suggesting possible mtDNA heteroplasmy of two closely related haplotypes. All the sequences were aligned using *mafft*^102^ and inputted to the ML tree inference tool *RAxML-NG*^117^. Both ML search and bootstrapping (n = 100) were performed under the GTR+I+G4 model (options: --all --bs-trees 100). We found Thorin falling outside the genetic variation of the already published late European Neanderthals and forming a clade with the Forbes’s Quarry Neanderthal and Stajnia S5000 with Mezmaiskaya 1 as the closest outgroup (Fig. 4).

**Maximum Parsimony (MP) analysis between Neanderthal mtDNA sequences**

We carried out a MP analysis including Denisova 3 and 23 of the Neanderthals used in the above ML analysis. Only the coding region ranging from 577 to 16,023 base pairs (bp) (rCRS coordinates) of the MT genome was considered for the analysis. The parsimony analysis was run in R using the *pratchet* function implemented in the R package *phangorn*^118^. From the alignment we computed a distance matrix based on raw substitution counts from which, we carried out a branch shortening analysis^8^ in order to estimate the relative ages of the samples. The ages were estimated using the radiocarbon dated sample, GoyetQ56-1 with the age of 42,540 years^16^, as a reference for calibration. To compute the age estimates, we used the inferred substitution rate for the coding region for modern humans at 1.57×10^-8^ substitutions/bp/year (95% confidence interval (CI): 1.17×10^-8^-1.98×10^-8^ substitutions/bp/year^119^.

We obtained a similar phylogeny as observed in the ML analysis (Fig. S10). We find Thorin in a clade with Forbes’s Quarry and Stajnia S5000, Thorin shares an additional substitution with Forbes’ Quarry (FQ) in comparison to Stajnia S5000. From the relative age estimates, Thorin has a similar age as FQ and is slightly younger than Stajnia S5000 (Fig. S12). However, the age estimated on FQ should be treated with caution, since only ~80% of the mt genome is available for analysis^49^. The obtained age for Thorin (95% confidence interval: ~80-100 ka (1,000 y ago)) is significantly older than the age of the stratigraphic layer from which it was excavated. A similar age discrepancy was found in the analysis of Stajnia S5000 and Chagyrskaya 8, which have both been molecularly dated to be older than what their respective radiometric contexts suggest^15,46^.

**Molecular tip dating and divergence date estimates from Neanderthal mtDNAs**

In addition to the parsimony analysis, we carried out a bayesian phylogenetic analysis using BEAST v. 2.6.3^45^. We used 87 mitochondrial sequences including 24 Neanderthals^16,11,33,110^, 1 Denisovan^120^, 9 ancient humans^105,106^, and 53 present-day humans^121^. We aligned the sequences with *mafft* and constrained our analysis to the non-D-loop region (rCRS coordinates: 577-16023 bp) in order to avoid the hypervariable regions of the genome. We first performed two sets of analysis, one with an estimated substitution rate and another with a fixed substitution rate at 1.57×10^-8^substitution/bp/year assuming a similar mitochondrial substitution rate as for modern humans.

We calibrated the tree using sample ages obtained from radiocarbon and molecular clock dating (Table S11). For the dated samples, we used the point estimate as the initial value and set a uniform prior distribution spanning the corresponding 95% confidence interval. For the undated Neanderthals we used an initial value of 50,000 years ago and a uniform prior ranging from 30,000 to 300,000 years ago. We further assigned a uniform prior for the time to the most recent ancestor (TMRCA) of all Neanderthals going from 100,000 to infinity and similarly assigned a uniform prior for the TMRCA of modern humans spanning from 50,000 years ago to infinity^37^. We set the age of present-day humans to 0 with no prior distribution.

In order to determine the optimal substitution model supporting our data, we used the BEAST package *bModelTest*^121^. For our data the best fitting model was Tamura-Nei 1993 (TN93)^146^ including rate heterogeneity and invariable sites. To determine the best fitting combination of a clock (strict clock or relaxed clock log normal) and a tree model (Coalescent Bayesian Skyline (CBS) or Coalescent Constant Population (CCP)), we performed a marginal like estimation (MLE) analysis for each model combination using stepping-stone (SS) sampling from the BEAST2 *model-selection* package with alpha set to 0.3 and a preBurnin percentage of 10%. For each clock/tree model combination, we ran 100 steps with a chain length of 15,000,000 in order to ensure convergence of the majority of the steps. Following ref. 147, we found that the models with CBS were strongly favored over CCP (fixed substitution rate: log_10_ BF > 30, estimated substitution rate: log_10_ BF > 26). We do not observe a notable difference between the strict and relaxed clock log normal models (fixed substitution rate: log_10_ BF > 0.17, estimated substitution rate: log_10_ BF > 0.4), however we chose the relaxed clock model as it is a slightly better fit and allows for varying substitution rates between branches.

We then performed four independent MCMC runs with 75,000,000 iterations and 225,000,000 iterations for the analysis with fixed and estimated substitution rates, respectively. We used a burn-in of 10%, while sampling parameters and trees for every 10,000 iterations. The independent runs were then combined respectively using *LogCombiner* in order to generate the final maximum clade credibility tree with *TreeAnnotator*.

When using the fixed substitution rate, we obtained an age of 106,170 years (95% HPD: 63,494 - 149,600) for Thorin placing it in the age range between the sequenced late and early Neanderthal. With the estimated substitution rate, we obtained a similar age although slightly younger, 100,160 years ago (95% HPD: 58,704 – 141,600; Fig. S12, Table S12). We estimated the mean substitution rate to 1.76×10^-8^. We estimated the divergence between Neanderthal and modern human lineages to ~390 ka with a fixed substitution rate and ~350 ka with an estimated substitution rate. The split date between “Thorin”-clade and the rest of the late Neanderthals was estimated to ~160 ka and ~150 ka with fixed and estimated substitution rate, respectively (Table S13, Fig. S12).

The age we obtained for Thorin in this analysis largely aligns with the resulting age estimate from the parsimony analysis. However, the molecular clock dating analysis does not agree with the ^14^C, U-series, and OSL ages obtained from the sediment layer Thorin was excavated from, nor with its isotopic indicators. As previously mentioned, there are two published Neanderthals, Chagyrskaya 8 and Stajnia S5000, for which a similar age discrepancy has been observed. A plausible genetic explanation behind this could be that the Neanderthals in this particular clade have a slightly slower substitution rate than the other late Neanderthals. To test this, we carried out another BEAST analysis using the same sequence alignment but adding Thorin as an additional calibration point with a mean age of 50 ky (95% CI: 45–55 ky) while estimating the substitution rate. We did this to test how the extra calibration point would affect the overall ages for the samples in the “Thorin”-clade as well as the corresponding divergence dates.

For the analysis we used TN93 as a substitution model allowing for rate heterogeneity and invariable sites. We used the relaxed clock model in order to obtain rate estimates for each branch of the tree and combined it with the CBS tree model, which was strongly favored over CCP (log_10_ BF > 26). We ran four independent runs of each 100,000,000 iterations of which 10 % was used as burn-in. Both tree and parameters were sampled for every 5,000 iterations. The runs were subsequently combined using *LogCombiner*, from which we generated the maximum clade credibility tree using *TreeAnnotator*.

With Thorin as an additional calibration point, we obtained a mean substitution rate of 1.82×10^-8^ substitutions/bp/year. For the late Neanderthals, we did not observe a significant change in their estimated ages, as they diverged from the “Thorin”-clade relatively early. However, we did find the ages for the early Neanderthals including Chagyrskaya 8 and Stajnia S5000 to be closer to their archaeological context ages. Similarly, we also estimated Mezmaiskaya 1 to be ~74 ka, which is also closer to its previous estimate of 60-70 ka (ref. 48; Table S12, Fig. S12). As a consequence, we find the Neanderthal-modern human split to be 20-50 ka later than our previous estimate (mean: 331 ka, 95% HPD: 253-417 ka), the Hohlenstein-Stadel divergence from the branch of the Altai Neanderthal to be ~215 ka (95% HPD: 163-274 ka) and the divergence between the “Thorin”-clade and the rest of the late Neanderthals to ~123 ka (95% HPD: 97-152 ka; Table S13, Fig. S12). Overall, the range of substitution rates increased slightly when using Thorin as a calibration as expected, but was still within a relatively narrow range (Fig. S13). Hence we conclude that a slower rate in the "Thorin"-clade would form a plausible explanation for the discrepancies observed between the molecular and contextual ages of the Neanderthal samples in that clade (Fig. S13).

**Supplementary Note 4 - Population genetic analyses**

**Procrustes analysis**

We used procrustes analysis^148^ to place the low-coverage Neanderthals within the context of a principal component analysis (PCA) of the three high coverage samples Altai Neanderthal, Vindija 33.19, and Denisova 3. We find that the projected samples form a cline along PC2 spanning from a central position between the two Neanderthals towards Vindija 33.19 (Fig. S14). The extremes of the cline are marked by the early European Neanderthals Hohlenstein-Stadel and Scladina towards the Altai Neanderthal, and the late European Neanderthals with the low coverage Vindija individual at the extreme towards Vindija 33.19, consistent with previous findings^16^. In comparison, Thorin projects closer to the Altai Neanderthal end of the cline than the late European Neanderthals and both Mezmaiskaya samples, in a similar position to the Forbes’ Quarry Neanderthal from Gibraltar (Fig. S14).

**D statistics**

In order to assess the relationship between the archaic individuals, we carried out different configurations of D-statistics in the form of *D (pop1, pop2; pop3, outgroup)*, to test whether two populations *pop1* and *pop2* form a clade with respect to a population *pop3*. We first tested for differences in levels of allele sharing between Neanderthals and present-day human populations by using *D* (Vindija 33.19/Altai Neanderthal, low; modern, Mbuti), where ‘low’ represents all the low coverage Neanderthals including Thorin, while ‘modern’ represents a number of contemporary and ancient individuals. The expectation for this statistic is D=0 if the pair of Neanderthals forms a clade with respect to the pair of modern human populations. Significant departures from D=0 indicate increased allele sharing between a modern human / Neanderthal pair, which can result either due to a true positive signal of gene flow, or alternatively a false positive signal because of contamination of the ancient sample with modern human DNA. To distinguish these possible scenarios, we carried out all D-tests by using either all fragments or restricting to only deaminated reads in the low coverage Neanderthal samples. To assess the possibility of reference bias we also add a configuration with the human reference genome as *pop3*. A significant departure from the tested null-hypothesis will indicate significantly more reference bias in the Thorin data compared to the other Neanderthals.

When using deaminated fragments only, we find that all low coverage Neanderthals and Vindija 33.19 are equally related to modern humans (D=0), consistent with previous results^16^ (Fig. S15). Hence, we do not find evidence of subsequent interbreeding between the Thorin-lineage and modern humans. Using all sample reads, we observe highly significant D-statistics for non-African modern human populations when paired with Neanderthal samples with high estimated contamination rates (Hohlenstein-Stadel, Scladina, Spy 94a, Mezmaiskaya 1), suggesting the bulk of their contamination derives from non-African modern sources. No differences between the two sets of statistics are observed in Neanderthals with low contamination on the other hand, including Thorin (Fig. S15).

We carried out a similar D-statistics using the Altai Neanderthal. D-statistics involving non-African modern human populations are shifted from D=0 also for deaminated reads for all European Neanderthals after 100 ka (excluding Hohlenstein-Stadel and Scladina, both of which show large standard errors due to limited amount of data). This is consistent with the previously described closer genetic similarity between the European Neanderthals and the Neanderthal that introgressed with modern humans. Since the Altai Neanderthal is only distantly related to the human-introgressing Neanderthal, we find a slightly higher allele sharing between the tested Neanderthals and the non-African modern human populations (Fig. S16). Results for Thorin are similar to the other European Neanderthals for both statistics, from which we can derive that the introgressing population diverged prior to the common ancestor of Thorin and the other European Neanderthals.

In the D-statistics discussed above, we also test for evidence of a higher level of reference bias in Thorin compared to the high coverage genome of Vindija 33.19. In neither tests and regardless of using all reads or only deaminated reads we do not find evidence of such. We also carried out a similar test with the configuration *D (Thorin, Neanderthal X; Human reference genome, Mbuti)* and found no evidence of significantly higher reference bias in Thorin compared to the other tested low coverage Neanderthals (Fig. S17). Likewise we test for potential capture bias in our data with the D test *D(capture, shotgun; reference, Mbuti)*, testing our capture reads used throughout this study and a set of reads generated without capture for significant allele sharing with the reference genome. We obtain a Z-score = -0.2 using all reads and rule out the possibility of capture bias in our analysis (Fig. S17).

We carried out the following D-test *D (Thorin, Neanderthal X; Han / French, Mbuti)* to further investigate the possibility of additional allele sharing between Thorin and modern humans (represented by Han and French populations) compared to the other low coverage Neanderthals. As we do not find significant deviations from the null-hypothesis, D≈0, we are ruling out the possibility of additional gene flow between the Thorin lineage and modern humans, consistent with previous D-statistics (Fig. S17).

We next quantified the relative amount of allele sharing of Thorin and other Neanderthals with the two high coverage samples Vindija 33.19 and Altai Neanderthal, using D-statistics of the form *D* (Altai Neanderthal, Vindija 33.19; Neanderthal X, Mbuti; Fig. 5). We find that all tested Neanderthals share significantly more alleles with Vindija 33.19 than with the Altai Neanderthals. Furthermore, the relative amount of excess sharing differs between samples, ranging from the lowest for the two early European Neanderthals Hohlenstein-Stadel and Scladina to the highest for the late European Neanderthal samples (Fig. 5). Thorin shares relatively fewer alleles with Vindija than all other Neanderthal samples after 80 ka, mirroring the results from the PCA analysis (Fig. S14). Using D-statistics of the form *D* (Vindija 33.19, Thorin; Neanderthal X, Mbuti; Figs. 5 & S18), and *D* (Vindija 33.19, Neanderthal X; Thorin, Mbuti) (Fig. S18), we confirm that Thorin is consistent with forming an outgroup with respect to other Neanderthal samples after 80 ka, including the ~70 ka Mezmaiskaya 1 individual from the Caucasus as well as the ~80 ka Chagyrskaya 8 individual from Siberia. Interestingly, the Gibraltar Neanderthal individual from Forbes’ Quarry was found significantly closer to Thorin than to Vindija 33.19. We caution that the signal was not significant when restricting to deaminated reads for Thorin, albeit this also resulted in a substantial reduction in the amount of data and large standard errors due to most libraries of Thorin being USER-treated (Fig. S18).

Finally, we tested for signature of gene flow from a “deep” lineage into late European Neanderthals using D-statistics of the form D (Vindija 33.19, Neanderthal X; Mezmaiskaya 2, Mbuti), D (Mezmaiskaya 2, Neanderthal X; Vindija 33.19, Mbuti), and D (Vindija 33.19, Mezmaiskaya 2; Neanderthal X, Mbuti). In the absence of such gene flow, the expectation for the first configuration is D=0 (i.e. Mezmaiskaya 2 forms an outgroup to other late European Neanderthals), whereas it is D<0 for the second configuration (i.e. Vindija 33.19 shares more alleles with late European Neanderthals than with Mezmaiskaya 2), while the expectation for the latter configuration is D>0 likewise due to allele sharing between Vindija 33.19 and late European Neanderthals. We find that the late Neanderthals Spy 94a, Goyet Q56-1, and the other Vindija individual are consistent with this expectation (Extended Data Figs. 8-9). For Les Cottés-Z4 1514 on the other hand, we reject D (Vindija 33.19, Neanderthal X; Mezmaiskaya 2, Mbuti) = 0 (Z = 8.0), indicating gene flow from a deep lineage forming an outgroup to Mezmaiskaya 2 and the other late Neanderthals (Extended Data Figs. 5 & 9).

**Treemix**

To infer a tree topology using the allele frequencies, we carried out a treemix analysis^136^. Due to the differences in the level of contamination, we restricted the analysis to deaminated fragments for samples with significant signs of modern contamination (Table S14). We, furthermore, use two high coverage Mbuti individuals from Simons Genome Diversity Panel^137^ (sample IDs: HGDP00456, SS6004471) as a root population. The analysis was carried out grouping SNPs into blocks of 300 bp (-k 300) to account for linkage disequilibrium without the default sample size correction (-noss).

From the model with no migration event, we find that Thorin is an outgroup to the rest of the late European Neanderthals as well as Mezmaiskaya 1 (Fig. S19). However, the relatively high residual values between some of the Neanderthals indicate that the initial model fails to fully resolve the relationships between the tested samples. First, we find a poor fit for Chagyrskaya 8 with respect to both the Altai Neanderthal and Vindija 33.19 indicating that Chagyrskaya is not modeled with a sufficient amount of shared drift to the two in this particular model. We observe a similar pattern between Mezmaiskaya 2 and Vindija 33.19 as well as with Les Cottés with respect to Thorin and Goyet Q56-1. On the other hand, we find that the model includes too much shared drift for Les Cottés Z4-1514 with respect to Vindija 33.19 and Mezmaiskaya 2. We therefore carried out another treemix analysis allowing for a migration event (Fig. S19). With this option treemix infers an admixture event between the Altai Neanderthal and Chagyrskaya resolving the poor fit between the two samples from the former model. Adding an additional migration event (m=2), we infer an admixture event between Vindija 33.19 and Goyet Q56-1 (Fig. S19). However, the poor fit between Les Cottés Z4-1514 with respect to Thorin persists.

**Runs of homozygosity**

We called runs of homozygosity (ROH) employing spatial smoothing by fused lasso regression of read allele frequencies along individual Neanderthal genomes. In this approach, the vector of minor read allele frequencies is used as a predictor variable in a regression that aims to estimate a piecewise constant vector of coefficients by penalizing differences in consecutive coefficients. In the context of calling ROHs, the goal of this approach is to find long stretches with coefficients estimated to b=0, corresponding to regions without evidence for the presence of two alleles at the SNPs considered. Sparsity in the coefficients and hence tolerance to false positive heterozygous SNPs due to sequencing error or ancient DNA damage can be adjusted through the regularization parameter l.

To test the feasibility of this approach to detect ROHs in low-coverage Neanderthal genomes, we called ROHs on simulated datasets of high-coverage genomes randomly down-sampled to lower coverage (2X and 1X average genomic coverage). We find that lower values of the regularization parameter (λ = 0.25 or λ = 0.5) result in cumulative ROH lengths that are highly correlated between subsampled and full genomes (Fig. S20). When using higher regularization (i.e. stronger emphasis on smoothing; λ = 0.75 or λ = 1), shorter segments tend to be missed in the subsampled genomes, resulting in a reduction of cumulative ROH length when using lower minimum ROH length (Fig. S20). Conversely, cumulative ROH length at higher minimum length cutoffs are overestimated, due to merging of nearby shorter fragments (Fig. S20, S21). Both of these effects are particularly pronounced in the more challenging lowest coverage scenario (1X). Based on these results, we used a regularization parameter of l = 0.5 in the final analyses.

**Demographic modeling**

The SFS-based method *momi2* was used to carry out demographic modeling from allele count data. Unlike other SFS-based methods, *momi2* allows to include low coverage samples represented by pseudohaploid genotypes in the inference, by using an SFS with entries conditional on being polymorphic in a set of high individuals only^50,128^. We therefore used an SFS with ascertainment restricted to the four high-quality archaic samples with diploid genotypes (Altai Neanderthal, Chagyrskaya 8, Vindija 33.19 and Denisovan) throughout.

We first fit a backbone demography including only the high-coverage individuals, incorporating previously inferred demographic events. Specifically, the model includes the following:

- A shared archaic lineage ancestral to Denisovans and Neanderthals diverging from modern humans

- Early Siberian Neanderthals (represented by the Altai Neanderthal) diverging earliest within the Neanderthal clade

- Introgression of a “super-archaic” ghost lineage into the Denisovan

- Gene flow from the Altai Neanderthal into the Denisovan

We assumed a shared ancestral Neanderthal population size for the internal edges within the Neanderthal clade, and individual size parameters for each Neanderthal sample at the leaf. Previous studies have demonstrated low effective population sizes and evidence for recent inbreeding in Neanderthal populations^8,15^, we therefore parameterized population sizes at the leaf as exponentially decaying from their divergence to the sample age. We performed ten independent replicate fits from random parameter starting values for this model to ensure convergence to a single optimum. We then added the two low coverage samples Thorin and Mezmaiskaya 1 to this initial model, as independent divergences from the Vindija 33.19 lineage and allowing it to occur at any time after its divergence from the Altai Neanderthal lineage. This new model was re-optimized in ten replicates, using the best fit parameter estimates from the initial model as starting values, and random initial values for the newly added demographic parameters. The resulting best-fit model yielded estimates consistent with previous results for the published Neanderthal samples^12,15^ (Fig. 6, Table S15).

To further investigate the relationship of the Forbes’ Quarry Neanderthal with Thorin, we added its data to the previously obtained best-fit model and re-optimized using the same procedure as above. We considered two different scenarios for the divergence of FQ. In the first scenario, the FQ lineage was allowed to diverge from the Vindija lineage at any time between its initial divergence from the ancestor with the Altai Neanderthal. The best-fitting model for this scenario resulted in a divergence of FQ at ~106 ka, slightly earlier than the divergence of the Thorin lineage (Fig. S23, left). In the second scenario, FQ was modeled to diverge from the Thorin lineage. This scenario resulted in a best fit of FQ diverging at ~81 ka from the Thorin lineage (Fig. S23, right). The final Log-likelihood of this model improved by 11 units compared to the independent FQ divergence (-2,149,760 for Thorin divergence; -2,149,771 for independent divergence), indicating that the FQ lineage is more closely related to Thorin than the other European Neanderthals.

We also performed additional models to investigate the evidence for admixture with a deep Neanderthal lineage in the ~43 ka Les Cottés Z4-1514 sample from France. We first fit a new model by adding the genomes of the late Neanderthals from Mezmaiskaya 2 and Les Cottés Z4-1514 to the best-fit model with Thorin described above. Based on the evidence from the D-statistic results, we modeled the Les Cottés Z4-1514 lineage as a mixture of a late European Vindija-like lineage (constrained to diverge after Mezmaiskaya 2), and an early European lineage (constrained to diverge prior to Mezmaiskaya 2). The resulting model fit the data well (all |Z| < 3 from *f*_4_* statistics) and resulted in a ~28% contribution to Les Cottés Z4-1514 from an early European lineage estimated to diverge prior to Mezmaiskaya 1 and Chagyrskaya 8, but after Thorin from the ancestral European Neanderthal stem (Fig. S24, left). To test whether this model fits significantly better than a model without admixture, we fit a nested model with the same configuration but the admixture proportion constrained to zero. The resulting model fit was significantly worse (Log-likelihood of full model -2,258,876, nested model with admixture proportion constrained to zero -2,258,948), and resulted in three outliers in the expected *f*_4_* statistics ((Fig. S24, middle), strongly suggesting the presence of “deep” European Neanderthal ancestry in Les Cottés Z4-1514. To further investigate whether this contribution was related to Thorin, we fit another model where the deep lineage was allowed to only diverge from the Thorin lineage. The best-fitting result showed a near-simultaneous divergence of the deep lineage immediately after the divergence of the Thorin lineage, and a substantially poorer model fit (Log-likelihood -2,258,887). Our results therefore suggest the presence of ancestry related to another deep European Neanderthal lineage in the genome of Les Cottés Z4-1514 which is distinct from the Thorin lineage.

**Supplementary Note 5 - Paleoproteomic analyses**

The diagnostic recognition of undisputable hominin clades is a key point in Paleolithic archaeology because during this time various distinct lineages, such as Anatomically Modern Humans (AMH), Neanderthals, or Denisovans, coexisted throughout Eurasia leading to difficulties in determining the spatio-temporal distribution of these populations, their eventual interactions or their technical and cultural abilities without clearly diagnostic hominin remains or specific biomolecular information. Fortunately, advances in ‘omic’ technology has revolutionized our understanding of human evolution, largely through the developments in genome sequencing capabilities^149^, providing us with mitochondrial and nuclear genomes not only of ancient anatomically modern humans^129,150,151^ but now also several extinct human lineages such as Neanderthals^8,11^ and Denisovans^114^ , the significance of which has been reviewed elsewhere^152^.

Biomolecular methods explore various paths for the recognition of these hominin populations and are of crucial importance when diagnostic hominin remains are lacking. These methods can help reach hominin attributions and understand some crucial scientific debates like the archaic hominin extinctions and their eventual relations at a period when AMH and Neanderthals may have, or not, directly interacted in Europe^3,4^. However, when it is available, paleogenetic research is usually limited to inferences from highly fragmentary DNA sequence data, often limited to mtDNA but with progressively more nuclear genomes being published. Nevertheless, introgressions between all west Eurasian Pleistocene populations - AMH, Neanderthals and Denisovans make their interpretations much more complex than expected^153,154^.

Other molecular methods have then been explored. The younger field of proteomics, or ‘paleoproteomics’ as it is being referred to in the study of ancient material, is then becoming an ever more popular tool for helping our understanding of the human past. This was initially through a relatively simplistic form of protein fingerprinting, containing enough information for species identification purposes, and more recently it has involved a shift to more sophisticated attempts at ‘sequencing’ rather than fingerprinting. This shift can result in improved taxonomic resolution and even offers the potential for recovering phylogenetic information of interest to the evolution of the long extinct beyond that of ancient DNA^155^. For example, the suggestion that Neanderthals made the Châtelperronian was supported for more than a century by most of the scientific community, with apparent cultural links and continuity between local Neanderthal industries and the beginning of the Upper Paleolithic^156-159^. This theory appeared to also be supported by the Arcy-sur-Cure (French Burgundy) Neanderthal teeth found in the 1960’s^160^ and then by the discovery of a partial Neanderthal skeleton in 1979 in the Châtelperronian “EJOP sup layer” of La Roche-à-Pierrot in Saint-Césaire (French Charente-Maritime)^161^. More recently however controversial issues emerged about these Neanderthal associations with the Châtelperronian culture^162^ and the attribution itself of the Saint-Césaire Neanderthal skeleton has now been considered undemonstrated at best^163^. The only remaining connection between Neanderthals and the Châtelperronian now appears to be solely based on the evidence from the Grotte du Renne site in Arcy-sur-Cure, a site however excavated more than half a century ago. However, the stratigraphic integrity of the Arcy site has itself been thoroughly debated due to radiocarbon results rendering the integrity of the Châtelperronian layers clearly disputable^164,165^. In that context, morphological information of the Arcy hominin remains^166^ should be used with some caution in terms of the scientific determination of the hominins who made the Châtelperronian culture. To overcome these limitations of the Arcy data, biomolecular analyses were carried out on directly-dated hominin material concluding that Neanderthals were the makers of the Châtelperronian in Arcy, with both DNA and protein ‘sequencing’^38^. However, the authors made it clear that the ancient DNA retrieved could have derived from Neanderthals or anatomically modern humans (“both specimens support the modern human state in more than 70% of their sequences” indicating that DNA from both were present), relying on a dubious measure of deamination to increase their speculations of it deriving from Neanderthal remains. They did however greatly strengthen their identifications of the specimens as deriving from Neanderthals using proteomics, with an unsubstantiated approach relying on semi-tryptic peptide matches on a newly proposed peptide biomarker^38^. More recently, such ‘paleoproteomic’ data were also used in support of the identifications of Denisovans in Tibet, employing the same/homologous peptide markers to those in the Arcy studies^167^, greatly expanding their geographic range. However, these apparent benefits in the level of information recovered with this ‘sequencing’ technology also come with severe caveats that need to be appreciated to temper our understanding of the limitations of ‘paleoproteomics’ and its influence in our interpretations of the human past.

The aims of part of this study were to investigate the limitations of paleoproteomics, in particular the extent to which proteomic approaches could generate misleading information. To demonstrate these issues, we can carry out semi-tryptic searches of the proteomes from Holocene AMH as well as presumed Middle Paleolithic Neanderthal remains to evaluate the error rate in identification.

Proteomes were recovered from the Holocene human specimens in three fractions per sample. All were treated with 0.6 M hydrochloric acid overnight (~18 hours), centrifuged at 12,400 rpm, and half of the supernatant ultrafiltered (10 kDa) into 50 mM ammonium bicarbonate (ABC) prior to reduction, alkylation and digestion (SOL fraction). The other half was precipitated in acetone at -20℃ overnight, centrifuged and resuspended in ABC for digestion as above (PREC). Then the acid-insoluble residue was incubated with 6 M guanidine hydrochloride (GuHCl) overnight prior to being ultrafiltered into ABC as above. In each case, reduction was carried out with 100 mM dithiothreitol (DTT) in 50 mM ABC (4.2 µL in 100 µL sample) at 60℃ for 10 minutes, allowed to cool, and acetylated with 100 mM iodoacetamide (8.4 µL in 100 µL sample) in the dark at room temperature for 45 minutes. The samples were further quenched with the same amount of DTT prior to digestion with 2 µg sequencing grade trypsin (Promega, UK) overnight (~18 hours) at 3℃. Sample digests (including one blank including filtered HCl, GuHCl and ABC) were then purified with C18 Solid Phase Extraction clean-up and dried to completion in a centrifugal evaporator prior to resuspension with 5% ACN + 0.1% formic acid (FA) and then analyzed using LC-MS/MS (Waters nanoAcquity UPLC system coupled to a Thermo Scientific Orbitrap Elite MS) at the Biological Mass Spectrometry Core Research Facility (University of Manchester) similar to methods described in ref. 168. In brief, samples were concentrated on a 20 mm x 180 μm pre-column prior to being separated on a 1.7 μM Waters nanoAcquity Ethylene Bridged Hybrid (BEH) C18 analytical column of (75 mm × 250 μm i.d.) and fractionation was achieved using a gradient beginning at 99% buffer A/1% buffer B and finishing at 75% buffer A/25% buffer B, whereby buffer A = 0.1% FA in H_2_O and buffer B = 0.1% FA in ACN. For this study, resulting MS/MS datafiles (.mgf) were searched against a local database made from the published sequences from refs. 38 and 167 using MASCOT v2.5.1. Datafiles were searched with semi-trypsin as the selected enzyme and using the following criteria: up to two missed cleavages, peptide tolerance of ± 5 ppm, MS/MS fragment ion mass value tolerance of 0.5 Da, a fixed carbamidomethyl modification of cysteine (mass shift = +57.02 Da), variable deamidation of asparagine (N) and glutamine (Q) modifications (mass shift = +0.98 Da; to allow for common diagenetic alterations), and variable oxidation of methionine (M), and hydroxylation of proline (P) and lysine (K) modifications (mass shift = +15.99 Da; equivalent mass to the process of hydroxylation).

***Potential False Positives at the Species Level***

In our analyses of three methods (acid-soluble fraction precipitated in acetone, acid soluble fraction ultrafiltered into ammonium bicarbonate, and incubation of acid-insoluble proteins in GuHCl as a denaturing buffer) from subsamples of three Holocene AMH skeletons (Supplementary Tables S16-S24), our results found that for at least two of the three anatomically modern humans analyzed, two of the three fractions gave a higher scoring protein match to the archaic COLX sequence using semi-tryptic searches than to the expected AMH sequence (Supplementary Table S25; raw data can be found at https://figshare.com/articles/dataset/Paleoprotomics_data_from_Neanderthals_Denisovans_and_anatomically_modern_humans/16676839/1). Our analyses of the Thorin Neanderthal specimens (Supplementary Tables S26-S35) also gave mixed results. Although our example spectrum does not yield a complete b or y ion series, those that are present around the amino acid substitution site (reflecting from both ends of the peptide) could readily be used to infer an archaic peptide despite knowing AMH as the source without setting in place standard criteria for confident semi-tryptic peptide matches. By contrast to archaeological samples, the blank run (Supplementary Table S36) produced an order of magnitude fewer peptide matches (e.g., ~4,000 vs >40,000) and Mascot scores typically of ~5-10, whereas ~20-100 was typically observed for our ancient samples (Supplementary Tables S16-S35).

Supplementary Tables

**Table S1. Radiocarbon determinations of Thorin obtained using the Oxford HYP compound-specific protocol**^44^. %C is the value of carbon on combustion. CN atomic ratios for HYP ought to be 5.0. See text for details.

| OxA | Used AG collagen (mg) | %C | δ^13^C (per mille) | δ^15^N (per mille) | CN atomic ratio | fM | ± |
| --- | --- | --- | --- | --- | --- | --- | --- |
| 37787 | 49.18 | 40.2 | -24.6 | 12.3 | 5.1 | 0.00193 | 0.00104 |
| 38388 | 25.9 | 37.6 | -23.3 | 13.8 | 5.0 | -0.0001 | 0.00123 |
| 38389 | 24.31 | 36.7 | -23.5 | 13.8 | 5.1 | -0.00052 | 0.00135 |

**Table S2. Summary of U-series ages values calculated by averaging each raster.** Values in red indicate data likely impacted by diagenetic processes, such as detrital thorium and/or uranium leaching depending on rasters. All ages are reported with a 2-sigma uncertainty.

|  | **R32/38** | **2SE** | **R34/38 (corr)** | **2SE** | **R30/38** | **2SE** | **U/Th** | **Age** | **2SE** |
| --- | --- | --- | --- | --- | --- | --- | --- | --- | --- |
| **Bison_1** | 0.0001 | 0.0003 | 1.0300 | 0.0230 | 0.2290 | 0.0410 | 102400 | **27.4** | **5.4** |
| **Bison_2** | -0.0000 | 0.0002 | 1.0380 | 0.0140 | 0.2580 | 0.0150 | -8708 | **31.1** | **2.0** |
| **Bison_3** | 0.0002 | 0.0002 | 1.0490 | 0.0230 | 0.2890 | 0.0290 | 21225 | **35.0** | **4.0** |
| **Bison_4** | 0.0005 | 0.0007 | 1.0430 | 0.0170 | 0.2840 | 0.0210 | 11100 | **34.6** | **2.9** |
| **Bison_5** | -0.0002 | 0.0002 | 1.0370 | 0.0150 | 0.1400 | 0.0210 | -22000 | **15.8** | **2.5** |
| **Bison_6** | 0.0001 | 0.0009 | 1.0320 | 0.0200 | 0.1350 | 0.0210 | -4160 | **15.3** | **2.5** |
| **Bison_7** | -0.0007 | 0.0003 | 1.0200 | 0.0190 | 0.0710 | 0.0180 | -1202 | **7.9** | **2.0** |
| **Bison_8** | 0.0004 | 0.0001 | 1.0369 | 0.0077 | 0.1980 | 0.0160 | 2100 | **23.1** | **2.0** |
| **Bison_9** | 0.0003 | 0.0001 | 1.0390 | 0.0100 | 0.2180 | 0.0230 | 2234 | **25.6** | **3.0** |
| **Bison_10** | 0.0004 | 0.0001 | 1.0353 | 0.0091 | 0.2070 | 0.0120 | 1893 | **24.3** | **1.5** |
| **Bison_11** | 0.0004 | 0.0001 | 1.0308 | 0.0067 | 0.2062 | 0.0091 | 1825 | **24.3** | **1.2** |
| **Bison_12** | 0.0008 | 0.0002 | 1.0266 | 0.0074 | 0.1170 | 0.0140 | 794 | **13.2** | **1.6** |
| **Bison_13** | 0.0006 | 0.0001 | 1.0247 | 0.0081 | 0.1330 | 0.0150 | 1036 | **15.2** | **1.8** |
| **Bison_14** | 0.0006 | 0.0001 | 1.0305 | 0.0091 | 0.1420 | 0.0150 | 1176 | **16.2** | **1.8** |
| **Thorin_1** | 0.0100 | 0.0037 | 1.0030 | 0.0590 | **0.0880** | 0.0630 | 510 |  |  |
| **Thorin_2** | -0.0040 | 0.0018 | 1.0820 | 0.0430 | **0.0890** | 0.0550 | -1043 |  |  |
| **Thorin_3** | -0.0029 | 0.0019 | 1.0380 | 0.0360 | 0.1790 | 0.0470 | 456 | **20.6** | **5.8** |
| **Thorin_4** | -0.0021 | 0.0016 | **0.9980** | 0.0510 | 0.1390 | 0.0640 | 535 | **16.4** | **7.9** |
| **Thorin_5** | **0.0420** | 0.0730 | 1.0400 | **0.2600** | 0.1000 | **1.6000** | **288** |  |  |
| **Thorin_6** | 0.0015 | 0.0043 | 1.0170 | 0.0760 | **0.0100** | **0.1500** | 968 |  |  |
| **Thorin_7** | 0.0056 | 0.0067 | 1.0150 | 0.0650 | **0.0150** | 0.0610 | 1229 |  |  |
| **Thorin_8** | **0.0530** | 0.0730 | 1.0700 | 0.0280 | 0.4000 | 0.0600 | **392** | **50.8** | **9.3** |
| **Thorin_9** | 0.0020 | 0.0002 | 1.0400 | 0.1100 | 0.3200 | 0.0490 | 400 | **40.0** | **7.7** |
| **Thorin_10** | 0.0032 | 0.0020 | 1.0630 | 0.0460 | 0.2370 | 0.0750 | 1567 | **27.4** | **9.5** |
| **Thorin_11** | 0.0010 | 0.0008 | 1.0340 | 0.0410 | 0.1110 | 0.0300 | 1182 | **12.4** | **3.4** |
| **Thorin_12** | 0.0007 | 0.0008 | 1.0950 | 0.0560 | 0.3620 | 0.0270 | 808 | **43.5** | **4.1** |
| **Thorin_13** | 0.0009 | 0.0004 | 1.0620 | 0.0300 | 0.2090 | 0.0400 | 833 | **23.8** | **4.9** |
| **Thorin_14** | 0.0010 | 0.0008 | 1.0440 | 0.0250 | 0.2190 | 0.0220 | 1989 | **25.6** | **2.8** |
| **Thorin_15** | 0.0004 | 0.0003 | 1.0950 | 0.0240 | 0.1180 | 0.0390 | 1023 | **12.4** | **4.3** |
| **Thorin_16** | 0.0007 | 0.0003 | **0.9910** | 0.0270 | 0.1420 | 0.0400 | 1859 | **16.9** | **5.0** |
| **Thorin_17** | 0.0003 | 0.0004 | **0.9720** | 0.0270 | 0.1040 | 0.0310 | 2957 | **12.3** | **3.8** |
| **Thorin_18** | 0.0001 | 0.0004 | **0.9750** | 0.0340 | **-0.0210** | 0.0250 | -3588 |  |  |
| **Thorin_19** | -0.0004 | 0.0005 | **0.9200** | 0.0290 | **-0.0590** | 0.0260 | -3455 |  |  |
| **Thorin_20** | -0.0003 | 0.0005 | **0.9480** | 0.0340 | **-0.0550** | 0.0250 | 55000 |  |  |

**Table S3. Summary of the US-ESR parameters and results for Thorin and bison fauna of Level B2.**

| **SAMPLE** | **Thorin** (GM-BP-SCU1) | **Bison** ( GM-BP-SCU2) |
| --- | --- | --- |
| **ENAMEL** | | |
| Dose (Gy)^a^ | 64.1±4.5  (66.3 +2.3/-6.7)* | 67.0±5.6  (68.8 +3.8/-7.4)* |
| U (ppm)^b^ | 2.9±0.5 | 4.4±0.6 |
| 234U/238U ^b^ | 1.2610±0.0992 | 1.1344±0.1002 |
| 230Th/234U ^b^ | 0.3337±0.1631 | 0.2755±0.1350 |
| Thickness (m) | 850±205 | 1022±298 |
| Water (%) | 3±1 | 3±1 |
| **DENTINE** | | |
| U (ppm) ^b^ | 3.6±0.5 | 9.46±0.4 |
| 234U/238U ^b^ | 1.0393±0.0044 | 1.0348±0.0076 |
| 230Th/234U ^b^ | 0.1948±0.07 | 0.2313±0.0344 |
| Water (%) | 5±3 | 5±3 |
| **SEDIMENT** | | |
| U (ppm) | 1.9±0. 1 | |
| Th (ppm) | 5.4±0.3 | |
| K (%) | 0.73±0.04 | |
| Water (%) | 12±5 | |
| **EXTERNAL DOSE RATE SEDIMENT** | | |
| Beta Dose (μGy a^-1^) | 119±16 | 101±17 |
| Gamma Dose (μGy a^-1^) | 500±70 | 500±70 |
| Cosmic (μGy a^-1^) | 166±14 | 166±14 |
| **COMBINED US-ESR AGE** | | |
| Internal dose rate  (μGy a^-1^) | 596±385 | 564±302 |
| Beta Dose dentine (μGy a^-1^) | 4±4 | 45±24 |
| P enamel ^c^ | -0.91±0.26 | -061±0.26 |
| P dentine ^c^ | -0.04±0.5 | -0.32±0.32 |
| Total Dose rate (μGy a^-1^) | 1381±392 | 1376±312 |
| EU ESR age (ka) ^d^ | 47±5 | 42±5 |
| LU ESR age (ka) ^d^ | 64±7 | 59±7 |
| CS-US age (ka) ^d^ | 56±41 | 56±39 |
| US-ESR age (ka) ^c*^ | 48 +5/-13 | 49 +5/-10 |

^a^Dose equivalent (D_e_) obtained using McDose 2.0

^b^Uranium concentration values were obtained by LA-MC-ICPMS (dentine values are averaged over the entire tooth).

^c^Ages were calculated using the program from ref. 169, with updated values from ref. 170.

^d^Ages were calculated using DATA program from ref. 171*.*

*The dose equivalent (D_e_) was modified to suit the US-ESR model, as described in the supplementary material text.

**Table S4. Library preparation and sequencing summary.**

| extract | USER ? | Library ID | PCR cycles (N) | shotgun or WGE | ng for WGE | post-WGE PCR cycles | Library | sequencing ID | sequencing run type |
| --- | --- | --- | --- | --- | --- | --- | --- | --- | --- |
| E1 | No USER | E1L1P1 | 6 | shotgun | - | - | Mandrin_1600_E1_3_TTAGGC | ESW_QFVP_Slimak_1600_E1_3 | HiSeq2500, 100SR |
|  |  |  |  |  |  |  | Mandrin_1600_E1_3_TTAGGC | ESW_SKHT_Slimak_1600_E1_3 | HiSeq2500, 80PE , 100PE |
|  |  | E1L1P2 | 10 | shotgun | - | - | Mandrin_1600_E1L1P2_7_CAGATC | ESW_FK7N_E1L1P2_7 | HiSeq2500, 80PE |
|  |  |  |  |  |  |  | Mandrin_1600_E1L1P2_7_CAGATC | ESW_JRYJ_E1L1P2_7 | HiSeq2500, 80PE |
|  |  |  |  |  |  |  | Mandrin_1600_E1L1P2_7_CAGATC | ESW_LYTK_E1L1P2_7 | HiSeq2500, 80PE |
|  |  |  |  | WGE | 70 | 13 | Mandrin_1600_E1L1P2_7_CAGATC | ESW_DCJY_E1L1P2_7_cap | HiSeq2500, 80PE |
|  |  |  |  |  |  |  | Mandrin_1600_E1L1P2_7_CAGATC | ESW_4PDU_E1L1P2_7_cap | HiSeq2500, 80PE |
|  |  |  |  |  |  |  | Mandrin_1600_E1L1P2_7_CAGATC | ESW_VLES_E1L1P2_7_cap | HiSeq2500, 80PE |
|  | USER | E1UL2P1 | 10 | shotgun | - | - | Mandrin_1600_E1UL2P1_10_TAGCTT_USER | ESW_FK7N_E1UL2P1_10 | HiSeq2500, 80PE |
|  |  |  |  |  |  |  | Mandrin_1600_E1UL2P1_10_TAGCTT_USER | ESW_PXQA_E1UL2P1_10 | HiSeq2500, 80PE |
|  |  |  |  |  |  |  | Mandrin_1600_E1UL2P1_10_TAGCTT_USER | ESW_JRYJ_E1UL2P1_10 | HiSeq2500, 80PE |
|  |  |  |  |  |  |  | Mandrin_1600_E1UL2P1_10_TAGCTT_USER | ESW_LYTK_E1UL2P1_10 | HiSeq2500, 80PE |
|  |  |  |  | WGE | 88 | 13 | Mandrin_1600_E1UL2P1_10_TAGCTT_USER | ESW_DCJY_E1UL2P1_10_cap | HiSeq2500, 80PE |
|  |  |  |  |  |  |  | Mandrin_1600_E1UL2P1_10_TAGCTT_USER | ESW_RQNQ_E1UL2P1_10_cap | HiSeq2500, 80PE |
|  |  |  |  |  |  |  | Mandrin_1600_E1UL2P1_10_TAGCTT_USER | ESW_4PDU_E1UL2P1_10_cap | HiSeq2500, 80PE |
|  |  |  |  |  |  |  | Mandrin_1600_E1UL2P1_10_TAGCTT_USER | ESW_VLES_E1UL2P1_10_cap | HiSeq2500, 80PE |
|  |  | E1UL2P2 | 10 | shotgun | - | - | Mandrin_1600_E1UL2P2_36_TGCAGG_USER | ESW_FK7N_E1UL2P2_36 | HiSeq2500, 80PE |
|  |  |  |  |  |  |  | Mandrin_1600_E1UL2P2_36_TGCAGG_USER | ESW_JRYJ_E1UL2P2_36 | HiSeq2500, 80PE |
|  |  |  |  |  |  |  | Mandrin_1600_E1UL2P2_36_TGCAGG_USER | ESW_LYTK_E1UL2P2_36 | HiSeq2500, 80PE |
|  |  |  |  | WGE | 185 | 12 | Mandrin_1600_E1UL2P2_36_TGCAGG_USER | ESW_DCJY_E1UL2P2_36_cap | HiSeq2500, 80PE |
|  |  |  |  |  |  |  | Mandrin_1600_E1UL2P2_36_TGCAGG_USER | ESW_4PDU_E1UL2P2_36_cap | HiSeq2500, 80PE |
|  |  |  |  |  |  |  | Mandrin_1600_E1UL2P2_36_TGCAGG_USER | ESW_VLES_E1UL2P2_36_cap | HiSeq2500, 80PE |
|  |  | E1UL3P1 | 10 | shotgun | - | - | Mandrin_1600_E1UL3P1_13_CTATCA_USER | ESW_PXQA_E1UL3P1_13 | HiSeq2500, 80PE |
|  |  |  |  |  |  |  | Mandrin_1600_E1UL3P1_13_CTATCA_USER | ESW_JRYJ_E1UL3P1_13 | HiSeq2500, 80PE |
|  |  |  |  | WGE | 104 | 12 | Mandrin_1600_E1UL3P1_13_CTATCA_USER | ESW_RQNQ_E1UL3P1_13_cap | HiSeq2500, 80PE |
|  |  |  |  |  |  |  | Mandrin_1600_E1UL3P1_13_CTATCA_USER | ESW_4PDU_E1UL3P1_13_cap | HiSeq2500, 80PE |
|  |  | E1UL3P2 | 10 | shotgun | - | - | Mandrin_1600_E1UL3P2_37_ACTGCC_USER | ESW_FK7N_E1UL3P2_37 | HiSeq2500, 80PE |
|  |  |  |  |  |  |  | Mandrin_1600_E1UL3P2_37_ACTGCC_USER | ESW_LYTK_E1UL3P2_37 | HiSeq2500, 80PE |
|  |  |  |  | WGE | 185 | 8 | Mandrin_1600_E1UL3P2_37_ACTGCC_USER | ESW_DCJY_E1UL3P2_37_cap | HiSeq2500, 80PE |
|  |  |  |  |  |  |  | Mandrin_1600_E1UL3P2_37_ACTGCC_USER | ESW_VLES_E1UL3P2_37_cap | HiSeq2500, 80PE |
| E2 | No USER | E2L1P1 | 14 | shotgun | - | - | Mandrin_1600_E2_12_CTTGTA | ESW_QFVP_Slimak_1600_E2_12 | HiSeq2500, 100SR |
|  |  |  |  |  |  |  | Mandrin_1600_E2_12_CTTGTA | ESW_SKHT_Slimak_1600_E2_12 | HiSeq2500, 80PE , 100PE |
|  |  | E2L1P2 | 10+8 | shotgun | - | - | Mandrin_1600_E2L1P2_8_ACTTGA | ESW_FK7N_E2L1P2_8 | HiSeq2500, 80PE |
|  |  |  |  |  |  |  | Mandrin_1600_E2L1P2_8_ACTTGA | ESW_JRYJ_E2L1P2_8 | HiSeq2500, 80PE |
|  |  |  |  | WGE | 111 | 12 | Mandrin_1600_E2L1P2_8_ACTTGA | ESW_DCJY_E2L1P2_8_cap | HiSeq2500, 80PE |
|  |  |  |  |  |  |  | Mandrin_1600_E2L1P2_8_ACTTGA | ESW_4PDU_E2L1P2_8_cap | HiSeq2500, 80PE |
|  |  | E2L2P1 | 12+10 | shotgun | - | - | Mandrin_1600_E2L2P1_39_CGACCT | ESW_FK7N_E2L2P1_39 | HiSeq2500, 80PE |
|  |  |  |  | WGE | 325 | 8 | Mandrin_1600_E2L2P1_39_CGACCT | ESW_DCJY_E2L2P1_39_cap | HiSeq2500, 80PE |
|  |  | E2L2P2 | 10+10 | shotgun | - | - | Mandrin_1600_E2L2P2_48_ACAGTC | ESW_PXQA_E2L2P2_48 | HiSeq2500, 80PE |
|  |  |  |  |  |  |  | Mandrin_1600_E2L2P2_48_ACAGTC | ESW_JRYJ_E2L2P2_48 | HiSeq2500, 80PE |
|  |  |  |  | WGE | 357 | 7 | Mandrin_1600_E2L2P2_48_ACAGTC | ESW_RQNQ_E2L2P2_48_cap | HiSeq2500, 80PE |
|  |  |  |  |  |  |  | Mandrin_1600_E2L2P2_48_ACAGTC | ESW_4PDU_E2L2P2_48_cap | HiSeq2500, 80PE |
|  | USER | E2UL2P1 | 10+8 | shotgun | - | - | Mandrin_1600_E2UL2P1_41_TGTCTG_USER | ESW_FK7N_E2UL2P1_41 | HiSeq2500, 80PE |
|  |  |  |  |  |  |  | Mandrin_1600_E2UL2P1_41_TGTCTG_USER | ESW_JRYJ_E2UL2P1_41 | HiSeq2500, 80PE |
|  |  |  |  |  |  |  | Mandrin_1600_E2UL2P1_41_TGTCTG_USER | ESW_LYTK_E2UL2P1_41 | HiSeq2500, 80PE |
|  |  |  |  | WGE | 650 | 11 | Mandrin_1600_E2UL2P1_41_TGTCTG_USER | ESW_DCJY_E2UL2P1_41_cap | HiSeq2500, 80PE |
|  |  |  |  |  |  |  | Mandrin_1600_E2UL2P1_41_TGTCTG_USER | ESW_4PDU_E2UL2P1_41_cap | HiSeq2500, 80PE |
|  |  |  |  |  |  |  | Mandrin_1600_E2UL2P1_41_TGTCTG_USER | ESW_VLES_E2UL2P1_41_cap | HiSeq2500, 80PE |
|  |  | E2UL2P2 | 8+8 | shotgun | - | - | Mandrin_1600_E2UL2P2_50_TTGAAC_USER | ESW_FK7N_E2UL2P2_50 | HiSeq2500, 80PE |
|  |  |  |  |  |  |  | Mandrin_1600_E2UL2P2_50_TTGAAC_USER | ESW_JRYJ_E2UL2P2_50 | HiSeq2500, 80PE |
|  |  |  |  |  |  |  | Mandrin_1600_E2UL2P2_50_TTGAAC_USER | ESW_LYTK_E2UL2P2_50 | HiSeq2500, 80PE |
|  |  |  |  | WGE | 615 | 8 | Mandrin_1600_E2UL2P2_50_TTGAAC_USER | ESW_DCJY_E2UL2P2_50_cap | HiSeq2500, 80PE |
|  |  |  |  |  |  |  | Mandrin_1600_E2UL2P2_50_TTGAAC_USER | ESW_4PDU_E2UL2P2_50_cap | HiSeq2500, 80PE |
|  |  |  |  |  |  |  | Mandrin_1600_E2UL2P2_50_TTGAAC_USER | ESW_VLES_E2UL2P2_50_cap | HiSeq2500, 80PE |
|  |  | E2UL3P1 | 10+8 | shotgun | - | - | Mandrin_1600_E2UL3P1_42_ACGTGC_USER | ESW_LYTK_E2UL3P1_42 | HiSeq2500, 80PE |
|  |  |  |  |  |  |  | Mandrin_1600_E2UL3P1_42_ACGTGC_USER | ESW_PXQA_E2UL3P1_42 | HiSeq2500, 80PE |
|  |  |  |  | WGE | 243 | 8 | Mandrin_1600_E2UL3P1_42_ACGTGC_USER | ESW_VLES_E2UL3P1_42_cap | HiSeq2500, 80PE |
|  |  |  |  |  |  |  | Mandrin_1600_E2UL3P1_42_ACGTGC_USER | ESW_RQNQ_E2UL3P1_42_cap | HiSeq2500, 80PE |
|  |  | E2UL3P2 | 9+8 | shotgun | - | - | Mandrin_1600_E2UL3P2_17_CGCTAT_USER | ESW_PXQA_E2UL3P2_17 | HiSeq2500, 80PE |
|  |  |  |  |  |  |  | Mandrin_1600_E2UL3P2_17_CGCTAT_USER | ESW_JRYJ_E2UL3P2_17 | HiSeq2500, 80PE |
|  |  |  |  | WGE | 500 | 8 | Mandrin_1600_E2UL3P2_17_CGCTAT_USER | ESW_RQNQ_E2UL3P2_17_cap | HiSeq2500, 80PE |
|  |  |  |  |  |  |  | Mandrin_1600_E2UL3P2_17_CGCTAT_USER | ESW_4PDU_E2UL3P2_17_cap | HiSeq2500, 80PE |
| E3 | No USER | E3L1P1 | 14 | shotgun | - | - | Mandrin_1600_E3_17_CGCTAT | ESW_QFVP_Slimak_1600_E3_17 | HiSeq2500, 100SR |
|  |  |  |  |  |  |  | Mandrin_1600_E3_17_CGCTAT | ESW_SKHT_Slimak_1600_E3_17 | HiSeq2500, 80PE , 100PE |
|  |  | E3L1P2 | 10+10 | shotgun | - | - | Mandrin_1600_E3L1P2_15_TGATCG | ESW_JRYJ_E3L1P2_15 | HiSeq2500, 80PE |
|  |  |  |  | WGE | 93 | 11 | Mandrin_1600_E3L1P2_15_TGATCG | ESW_4PDU_E3L1P2_15_cap | HiSeq2500, 80PE |
|  |  | E3L2P1 | 12+10 | shotgun | - | - | Mandrin_1600_E3L2P1_40_TGACGT | ESW_PXQA_E3L2P1_40 | HiSeq2500, 80PE |
|  |  |  |  |  |  |  | Mandrin_1600_E3L2P1_40_TGACGT | ESW_JRYJ_E3L2P1_40 | HiSeq2500, 80PE |
|  |  |  |  | WGE | 162 | 8 | Mandrin_1600_E3L2P1_40_TGACGT | ESW_RQNQ_E3L2P1_40_cap | HiSeq2500, 80PE |
|  |  |  |  |  |  |  | Mandrin_1600_E3L2P1_40_TGACGT | ESW_4PDU_E3L2P1_40_cap | HiSeq2500, 80PE |
|  |  | E3L2P2 | 12+10 | shotgun | - | - | Mandrin_1600_E3L2P2_49_CATCGT | ESW_FK7N_E3L2P2_49 | HiSeq2500, 80PE |
|  |  |  |  |  |  |  | Mandrin_1600_E3L2P2_49_CATCGT | ESW_JRYJ_E3L2P2_49 | HiSeq2500, 80PE |
|  |  |  |  |  |  |  | Mandrin_1600_E3L2P2_49_CATCGT | ESW_LYTK_E3L2P2_49 | HiSeq2500, 80PE |
|  |  |  |  | WGE | 175 | 9 | Mandrin_1600_E3L2P2_49_CATCGT | ESW_DCJY_E3L2P2_49_cap | HiSeq2500, 80PE |
|  |  |  |  |  |  |  | Mandrin_1600_E3L2P2_49_CATCGT | ESW_4PDU_E3L2P2_49_cap | HiSeq2500, 80PE |
|  |  |  |  |  |  |  | Mandrin_1600_E3L2P2_49_CATCGT | ESW_VLES_E3L2P2_49_cap | HiSeq2500, 80PE |
|  | USER | E3UL2P1 | 10+10 | shotgun | - | - | Mandrin_1600_E3UL2P1_43_TGATCC_USER | ESW_FK7N_E3UL2P1_43 | HiSeq2500, 80PE |
|  |  |  |  | WGE | 360 | 8 | Mandrin_1600_E3UL2P1_43_TGATCC_USER | ESW_DCJY_E3UL2P1_43_cap | HiSeq2500, 80PE |
|  |  | E3UL2P2 | 10+9 | shotgun | - | - | Mandrin_1600_E3UL2P2_18_TGAACA_USER | ESW_PXQA_E3UL2P2_18 | HiSeq2500, 80PE |
|  |  |  |  |  |  |  | Mandrin_1600_E3UL2P2_18_TGAACA_USER | ESW_JRYJ_E3UL2P2_18 | HiSeq2500, 80PE |
|  |  |  |  | WGE | 560 | 8 | Mandrin_1600_E3UL2P2_18_TGAACA_USER | ESW_RQNQ_E3UL2P2_18_cap | HiSeq2500, 80PE |
|  |  |  |  |  |  |  | Mandrin_1600_E3UL2P2_18_TGAACA_USER | ESW_4PDU_E3UL2P2_18_cap | HiSeq2500, 80PE |
|  |  | E3UL3P1 | 10+10 | shotgun | - | - | Mandrin_1600_E3UL3P1_44_CTCTAG_USER | ESW_FK7N_E3UL3P1_44 | HiSeq2500, 80PE |
|  |  |  |  |  |  |  | Mandrin_1600_E3UL3P1_44_CTCTAG_USER | ESW_LYTK_E3UL3P1_44 | HiSeq2500, 80PE |
|  |  |  |  | WGE | 455 | 8 | Mandrin_1600_E3UL3P1_44_CTCTAG_USER | ESW_DCJY_E3UL3P1_44_cap | HiSeq2500, 80PE |
|  |  |  |  |  |  |  | Mandrin_1600_E3UL3P1_44_CTCTAG_USER | ESW_VLES_E3UL3P1_44_cap | HiSeq2500, 80PE |
|  |  | E3UL3P2 | 10+8 | shotgun | - | - | Mandrin_1600_E3UL3P2_19_GTATCT_USER | ESW_FK7N_E3UL3P2_19 | HiSeq2500, 80PE |
|  |  |  |  |  |  |  | Mandrin_1600_E3UL3P2_19_GTATCT_USER | ESW_LYTK_E3UL3P2_19 | HiSeq2500, 80PE |
|  |  |  |  | WGE | 610 | 8 | Mandrin_1600_E3UL3P2_19_GTATCT_USER | ESW_DCJY_E3UL3P2_19_cap | HiSeq2500, 80PE |
|  |  |  |  |  |  |  | Mandrin_1600_E3UL3P2_19_GTATCT_USER | ESW_VLES_E3UL3P2_19_cap | HiSeq2500, 80PE |

**Table S5. Sequencing summary statistics.**

| **Library** | **# SE or PE** | **#Read or Read pairs** | **#Collapsed** | **%Clonality(mtDNA)** | **%HQUniqEndogenous(mtDNA)** | **Coverage(mtDNA)** | **Length(mtDNA)** | **%Clonality(nucDNA)** | **%HQUniqEndogenous(nucDNA)** | **Coverage**  **(nucDNA)** | **Length**  **(nuDNA)** |
| --- | --- | --- | --- | --- | --- | --- | --- | --- | --- | --- | --- |
| Mandrin_1600_E1L1P2_7_CAGATC | PE | 262534839 | 221604041 | 94.7519 | 0.0117 | 90.3280 | 50.9742 | 89.2897 | 2.5277 | 0.1542 | 75.110 |
| Mandrin_1600_E1UL2P1_10_TAGCTT_USER | PE | 319353856 | 255195158 | 95.2777 | 0.0093 | 75.9908 | 47.3897 | 90.5122 | 1.8678 | 0.1234 | 71.401 |
| Mandrin_1600_E1UL2P2_36_TGCAGG_USER | PE | 205432671 | 163789514 | 86.7192 | 0.0224 | 118.4697 | 47.8241 | 72.7481 | 4.9472 | 0.2130 | 72.473 |
| Mandrin_1600_E1UL3P1_13_CTATCA_USER | PE | 107506617 | 86771259 | 85.8666 | 0.0307 | 87.2629 | 47.8187 | 76.6296 | 5.1787 | 0.1193 | 72.327 |
| Mandrin_1600_E1UL3P2_37_ACTGCC_USER | PE | 164437290 | 134062850 | 86.5682 | 0.0287 | 123.0666 | 47.4018 | 71.6642 | 5.8891 | 0.2066 | 72.416 |
| Mandrin_1600_E1_3_TTAGGC | PE & SE | 647454523 | 551300936 | 9.4452 | 0.0009 | 13.0692 | 44.4539 | 9.1564 | 1.6481 | 0.1587 | 54.055 |
| Mandrin_1600_E2L1P2_8_ACTTGA | PE | 168547552 | 132175268 | 99.4823 | 0.0016 | 6.0668 | 44.6968 | 93.0949 | 1.1059 | 0.0252 | 49.888 |
| Mandrin_1600_E2L2P1_39_CGACCT | PE | 189003580 | 159103410 | 97.9574 | 0.0042 | 20.9779 | 44.8265 | 74.4747 | 2.8533 | 0.0896 | 53.064 |
| Mandrin_1600_E2L2P2_48_ACAGTC | PE | 319480334 | 269223510 | 98.6365 | 0.0027 | 23.6374 | 46.1489 | 80.6545 | 2.2415 | 0.1186 | 52.631 |
| Mandrin_1600_E2UL2P1_41_TGTCTG_USER | PE | 263682361 | 211439613 | 91.4752 | 0.0096 | 59.5107 | 42.7701 | 52.5479 | 3.1094 | 0.1219 | 50.537 |
| Mandrin_1600_E2UL2P2_50_TTGAAC_USER | PE | 244341910 | 202019419 | 93.5431 | 0.0112 | 64.8563 | 43.1379 | 61.9893 | 3.5573 | 0.1331 | 51.785 |
| Mandrin_1600_E2UL3P1_42_ACGTGC_USER | PE | 223790514 | 185966443 | 94.0984 | 0.0153 | 80.3455 | 42.9090 | 76.4073 | 2.9039 | 0.1005 | 52.871 |
| Mandrin_1600_E2UL3P2_17_CGCTAT_USER | PE | 208205671 | 170018451 | 91.9168 | 0.0165 | 84.2148 | 43.0276 | 57.6366 | 4.4598 | 0.1506 | 53.218 |
| Mandrin_1600_E2_12_CTTGTA | PE & SE | 898739326 | 703966635 | 88.6684 | 0.0003 | 4.7751 | 42.0711 | 88.2102 | 0.3390 | 0.0355 | 46.037 |
| Mandrin_1600_E3L1P2_15_TGATCG | PE | 63943719 | 53362939 | 99.5011 | 0.0025 | 4.3494 | 47.8429 | 92.4246 | 1.4725 | 0.0151 | 53.299 |
| Mandrin_1600_E3L2P1_40_TGACGT | PE | 336933066 | 262787435 | 99.3795 | 0.0011 | 11.9406 | 48.0490 | 90.7157 | 0.8417 | 0.0540 | 54.379 |
| Mandrin_1600_E3L2P2_49_CATCGT | PE | 466943536 | 369479937 | 99.4949 | 0.0009 | 14.4007 | 50.3130 | 92.9552 | 0.6815 | 0.0616 | 55.336 |
| Mandrin_1600_E3UL2P1_43_TGATCC_USER | PE | 117695601 | 92670434 | 90.8606 | 0.0145 | 44.5003 | 43.9287 | 51.5701 | 3.2197 | 0.0652 | 54.042 |
| Mandrin_1600_E3UL2P2_18_TGAACA_USER | PE | 249083130 | 201255722 | 96.4769 | 0.0070 | 46.5842 | 44.3048 | 68.8415 | 2.6336 | 0.1136 | 53.418 |
| Mandrin_1600_E3UL3P1_44_CTCTAG_USER | PE | 261543603 | 208436072 | 96.8987 | 0.0072 | 49.1452 | 43.7312 | 70.0276 | 2.6386 | 0.1162 | 52.636 |
| Mandrin_1600_E3UL3P2_19_GTATCT_USER | PE | 138746544 | 135673712 | 94.7869 | 0.0121 | 43.6771 | 44.1452 | 62.7581 | 3.5430 | 0.0846 | 54.506 |
| Mandrin_1600_E3_17_CGCTAT | PE & SE | 44899510 | 38195346 | 58.2189 | 0.0026 | 2.7062 | 44.7420 | 58.7172 | 1.9883 | 0.0121 | 49.283 |

PE: sequencing using Paired-End mode, SE: sequencing using Single-Read mode

Clonality refers to the fraction of aligned reads identified as PCR or optical duplicates.

Endogenous DNA content and mitochondrial and nuclear genome coverage are provided following duplicate removal.

**Table S6. Samples used for Y chromosome analysis.**

| Sample ID | Population ID | Country | Region | Coverage | Reference |
| --- | --- | --- | --- | --- | --- |
| UstIshim | UstIshim | Russia | Siberia | 21X | 129 |
| GRC13292545 | Mbo | Cameroon | WestAfrica | 8X | 172 |
| GRC13292546 | Mbo | Cameroon | WestAfrica | 11X | 172 |
| Mota | Mota | Ethiopia | Africa | 1.5X | 173 |
| SS6004473 | Ju_hoan_North | Namibia | SouthAfrica | 12X | 137 |
| SS6004471 | Mbuti | Congo | CentralAfrica | 11X | 137 |
| SS6004475 | Yoruba | Nigeria | WestAfrica | 11X | 137 |
| SS6004470 | Mandenka | Senegal | WestAfrica | 12X | 137 |
| SS6004480 | Dinka | Sudan | EastAfrica | 11X | 137 |
| SS6004474 | Sardinian | Italy (Sardinia) | SouthEurope | 10X | 137 |
| SS6004468 | French | France | WestEurope | 12X | 137 |
| SS6004467 | Dai | China | EastAsia | 11X | 137 |
| SS6004469 | Han | China | EastAsia | 11X | 137 |
| SS6004476 | Karitiana | Brazil | SouthAmerica | 10X | 137 |
| SS6004477 | Australian | Australia | NorthEastAustralia | 12X | 137 |
| SS6004472 | Papuan | PapuaNewGuinea | Oceania | 13X | 137 |
| SIII | SIII | Russia | Europe | 1.5X | 9 |
| Spy94a | Spy94a | Belgium | Europe | 0.1X | 16 |
| Mezmaiskaya2 | Mezmaiskaya2 | Russia | Europe | 1.8X | 16 |
| Yana | Yana | Russia | Siberia | 3X | 104 |
| KolymaRiver | Kolyma_River | Russia | Siberia | 1.8X | 104 |
| Denisova4 | Denisova 4 | Russia | Siberia | 0.2X | 123 |
| Denisova8 | Denisova 8 | Russia | Siberia | 0.4X | 123 |
| Thorin | Thorin | France | Europe | 0.2X | This study |

**Table S7.**

| Sample ID | Population ID | Country | Region | cov_group | Reference |
| --- | --- | --- | --- | --- | --- |
| vindija2010. | Vindija2010 | Croatia | Europe | low | 11 |
| Denisova | Denisova | Russia | EastEurope | high | 114 |
| NE1 | NE1 | Hungary | CentralEurope | high | 88 |
| Loschbour | Loschbour | Luxembourg | WestEurope | high | 127 |
| LBK | Stuttgart | Germany | WestEurope | high | 127 |
| Ust_Ishim | UstIshim | Russia | Siberia | high | 129 |
| Clovis | Anzick-1 | USA | NorthAmerica | high | 174 |
| AltaiNeandertal | AltaiNea | Russia | EastEurope | high | 8 |
| mezmaiskaya1 | Mezmaiskaya 1 | Russia | Caucasus | low | 8 |
| Bichon | Bichon | Switzerland | WestEurope | high | 126 |
| KK1 | KK1 | Georgia | WestEurope | high | 126 |
| HGDP01029 | Ju_hoan_North | Namibia | SouthAfrica | high | 137 |
| SS6004473 | Ju_hoan_North | Namibia | SouthAfrica | high | 137 |
| HGDP01284 | Mandenka | Senegal | WestAfrica | high | 137 |
| SS6004470 | Mandenka | Senegal | WestAfrica | high | 137 |
| SS6004475 | Yoruba | Nigeria | WestAfrica | high | 137 |
| HGDP00927 | Yoruba | Nigeria | WestAfrica | high | 137 |
| HGDP00456 | Mbuti | Congo | CentralAfrica | high | 137 |
| SS6004471 | Mbuti | Congo | CentralAfrica | high | 137 |
| DNK02 | Dinka | Sudan | EastAfrica | high | 137 |
| SS6004480 | Dinka | Sudan | EastAfrica | high | 137 |
| HGDP00665 | Sardinian | Italy | SouthEurope | high | 137 |
| SS6004474 | Sardinian | Italy | SouthEurope | high | 137 |
| HGDP00521 | French | France | WestEurope | high | 137 |
| SS6004468 | French | France | WestEurope | high | 137 |
| HGDP01307 | Dai | China | EastAsia | high | 137 |
| SS6004467 | Dai | China | EastAsia | high | 137 |
| HGDP00778 | Han | China | EastAsia | high | 137 |
| SS6004469 | Han | China | EastAsia | high | 137 |
| SS6004479 | Mixe | Mexico | CentralAmerica | high | 137 |
| LP6005443-DNA_E11 | Mixe | Mexico | CentralAmerica | high | 137 |
| LP6005443-DNA_F11 | Mixe | Mexico | CentralAmerica | high | 137 |
| HGDP00998 | Karitiana | Brazil | SouthAmerica | high | 137 |
| SS6004476 | Karitiana | Brazil | SouthAmerica | high | 137 |
| LP6005441-DNA_A12 | Surui | Brazil | SouthAmerica | high | 137 |
| LP6005441-DNA_B12 | Surui | Brazil | SouthAmerica | high | 137 |
| SS6004477 | Australian | Australia | Australia | high | 137 |
| SS6004478 | Australian | Australia | Australia | high | 137 |
| HGDP00542 | Papuan | PNG | Oceania | high | 137 |
| SS6004472 | Papuan | PNG | Oceania | high | 137 |
| GB20 | Mota | Ethiopia | EastAfrica | high | 137 |
| WC1 | WC1 | Iran | WestAsia | high | 124 |
| Bar8 | Bar8 | Turkey | WestAsia | high | 125 |
| SIII | SIII | Russia | WestAsia | high | 9 |
| Vindija33.19 | Vindija33.19 | Croatia | Europe | high | 12 |
| Yamnaya | Yamnaya | Caucasus | EastEurope | high | 128 |
| BOT2016 | BOT2016 | Kazakhstan | EastEurope | high | 128 |
| EBA2 | EBA2 | Kazakhstan | EastEurope | high | 128 |
| USR1 | USR1 | USA | NorthAmerica | high | 130 |
| goyetQ56-1 | Goyet Q56-1 | Belgium | Europe | low | 16 |
| spy94a | Spy 94a | Belgium | Europe | low | 16 |
| lescottes-Z4-1514 | Les Cottés-Z4-1514 | France | Europe | low | 16 |
| mezmaiskaya2 | Mezmaiskaya 2 | Russia | Caucasus | low | 16 |
| hybrid | Denisova 11 | Russia | Siberia | low | 131 |
| NEO240 | NEO240 | Russia | EastAsia | high | 104 |
| Yana | Yana | Russia | Siberia | high | 104 |
| Yana_old2 | Yana2 | Russia | Siberia | high | 104 |
| Kolyma_River | Kolyma_River | Russia | Siberia | high | 104 |
| hohlenstein-stadel | Hohlenstein-Stadel | Germany | Europe | low | 13 |
| scladina | Scladina | Belgium | Europe | low | 13 |
| FQ | FQ | Gibraltar | Europe | low | 49 |
| Chagyrskaya-Phalanx | Chagyrskaya 8 | Russia | Siberia | high | 15 |
| Stajnia_S5000 | Stajnia S5000 | Poland | Europe | low | 46 |
| Estatuas_nonHST-like_merged | Estatuas non-HST-like (merged) | Spain | Europe | low | 14 |
| Estatuas_pit1Layer2 | Estatuas p1L2 (non-HST-like) | Spain | Europe | low | 14 |
| Estatuas_pit1Layer3 | Estatuas p1L3 (non-HST-like) | Spain | Europe | low | 14 |
| Estatuas_pit1Layer4 | Estatuas p1L4 (HST-like) | Spain | Europe | low | 14 |
| Estatuas_pit2Layer2 | Estatuas p2L2 (non-HST-like) | Spain | Europe | low | 14 |
| Thorin | Thorin | France | Europe | low | This study |

**Table S8. C to T and G to A substitution rates.**
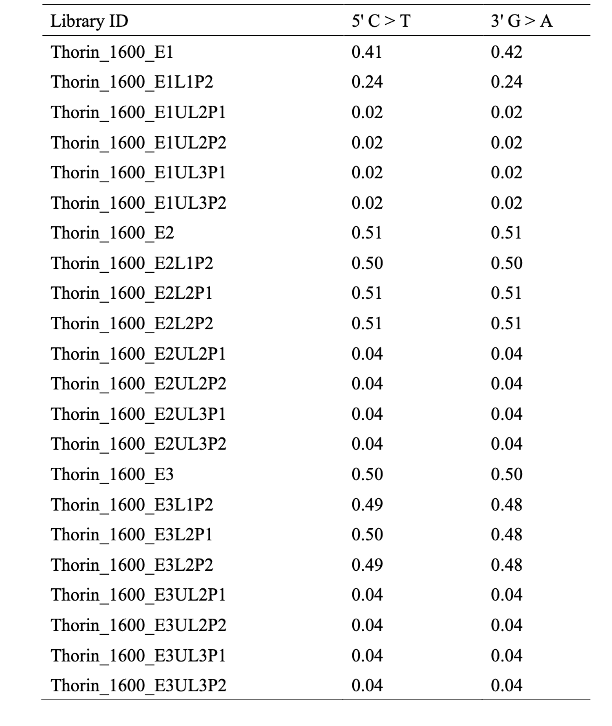

**Table S9. Proportion of authentic mitochondrial DNA.**
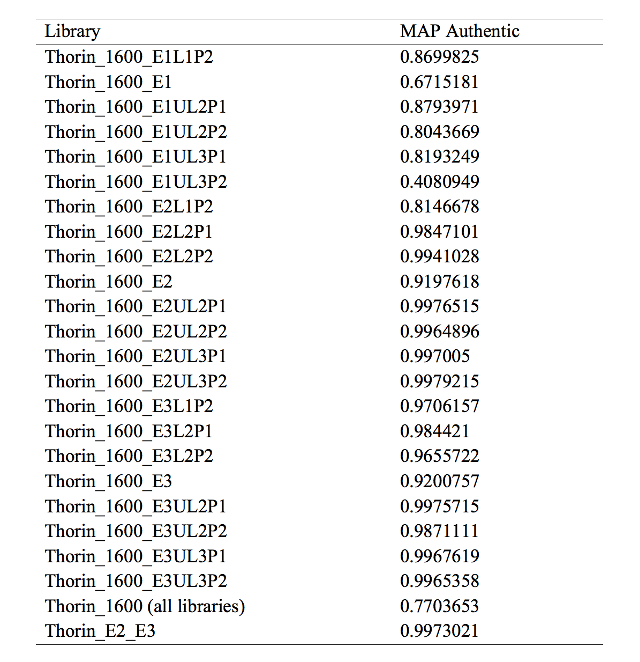

**Table S10. Maximum-Likelihood estimations of X chromosomal contamination.**

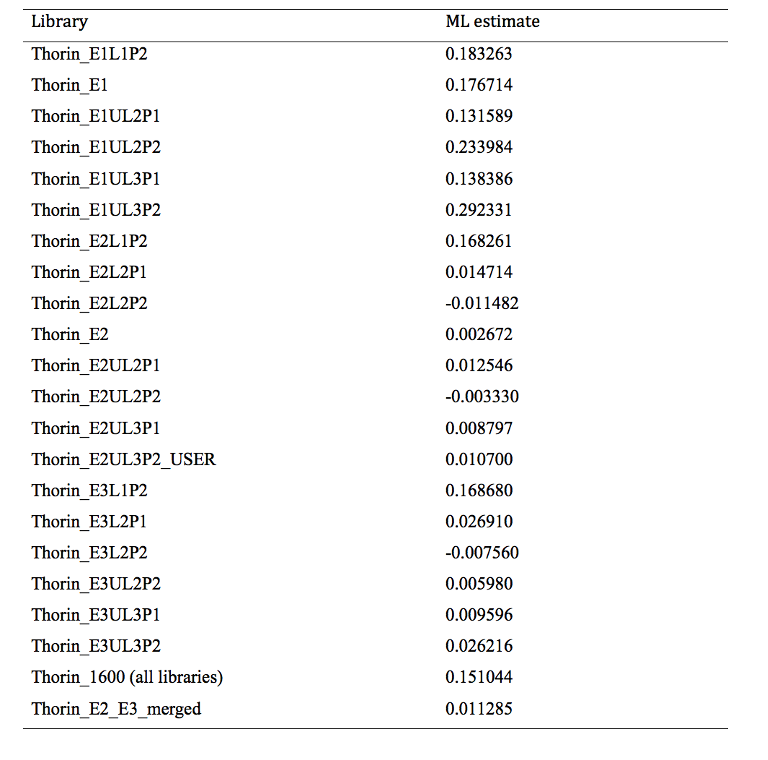

**Table S11. List of samples with radiocarbon dates used for calibration.**
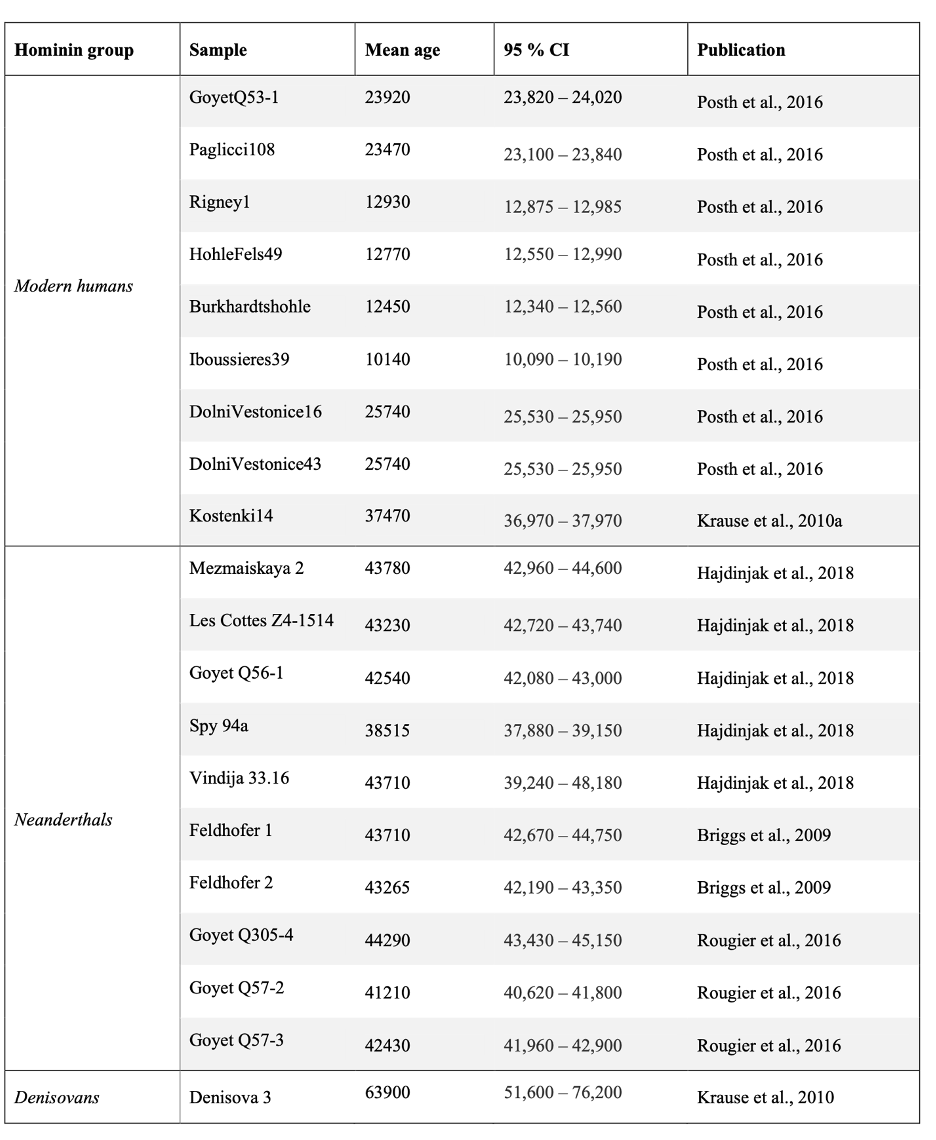

**Table S12. Age estimates from BEAST analysis.**

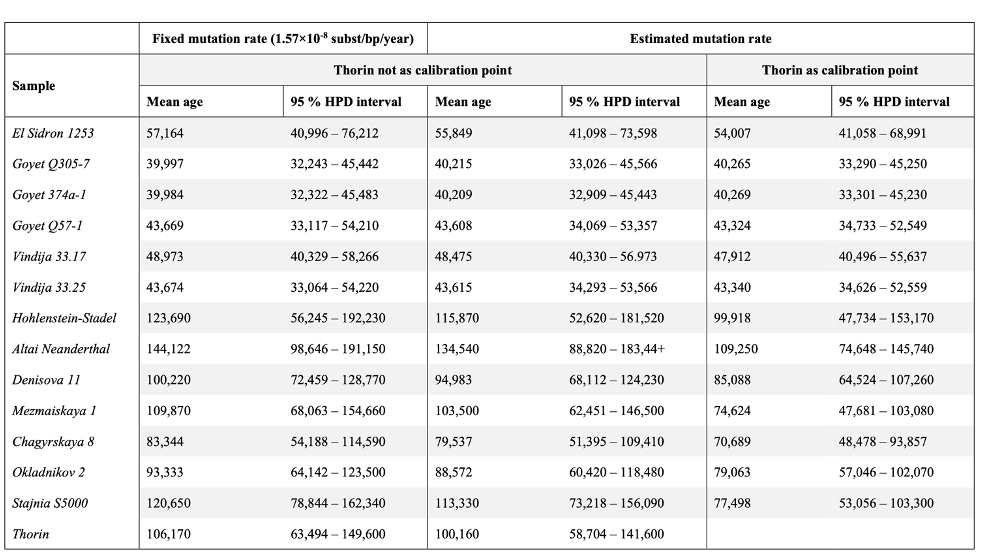

**Table S13. Divergence date estimates with comparison to previous studies.**

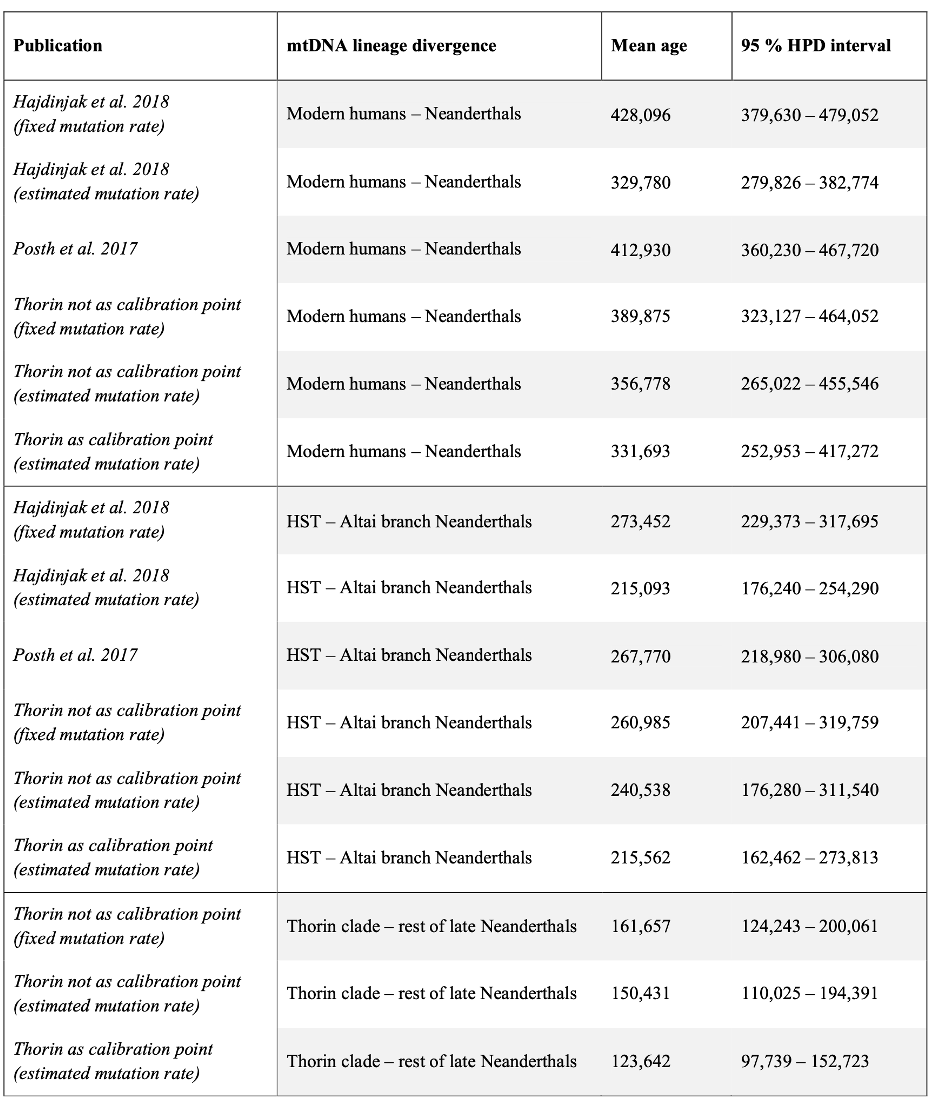

**Table S14. Samples included in treemix analysis.**
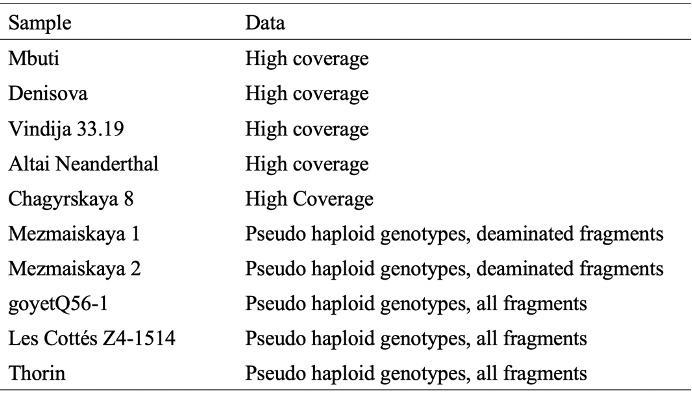

**Table S15. Demographic modeling parameter estimates**

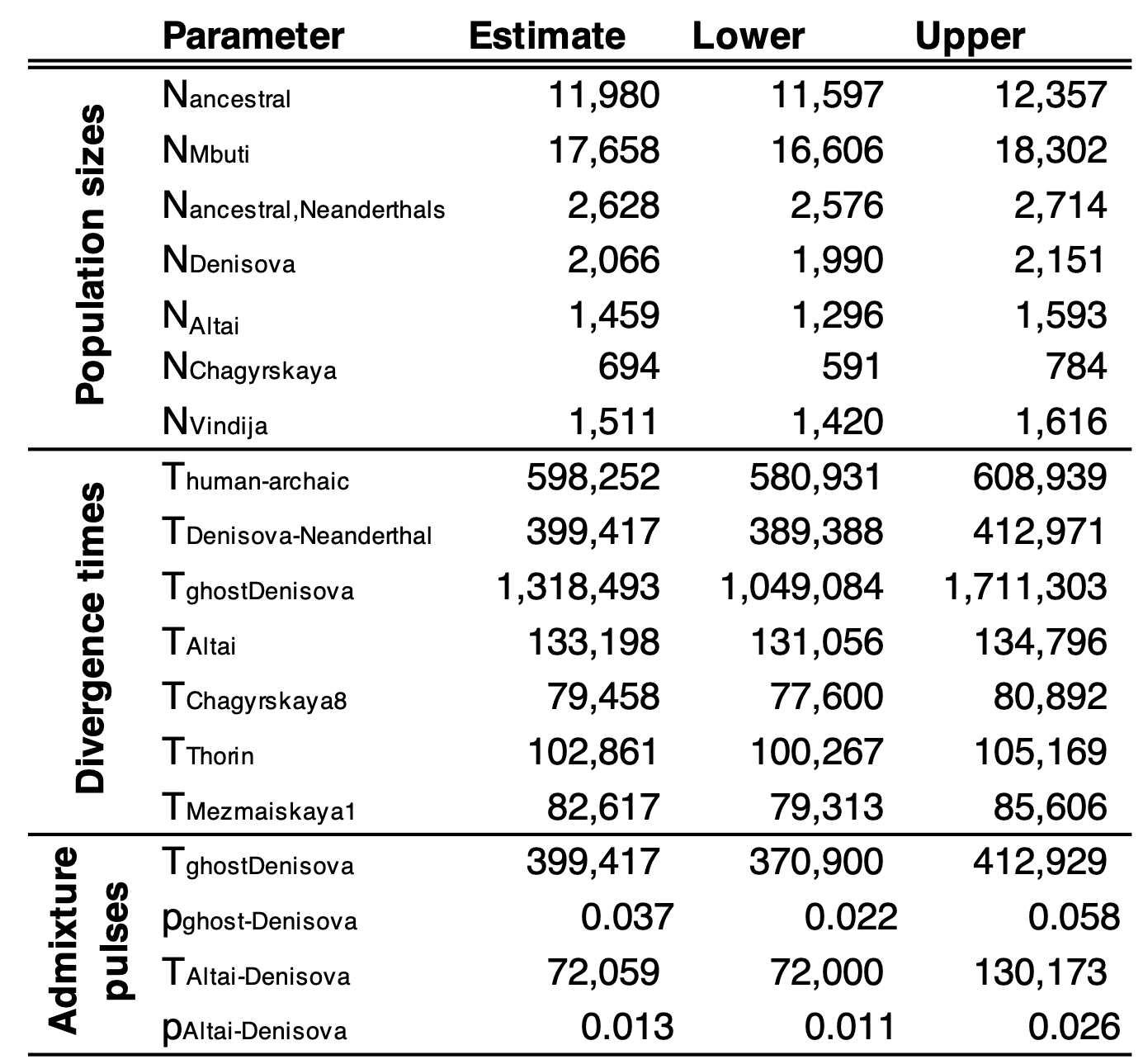

**For Tables S16 through S36, please see separate .CSV files in online Supplemental Information**

Supplementary Figures

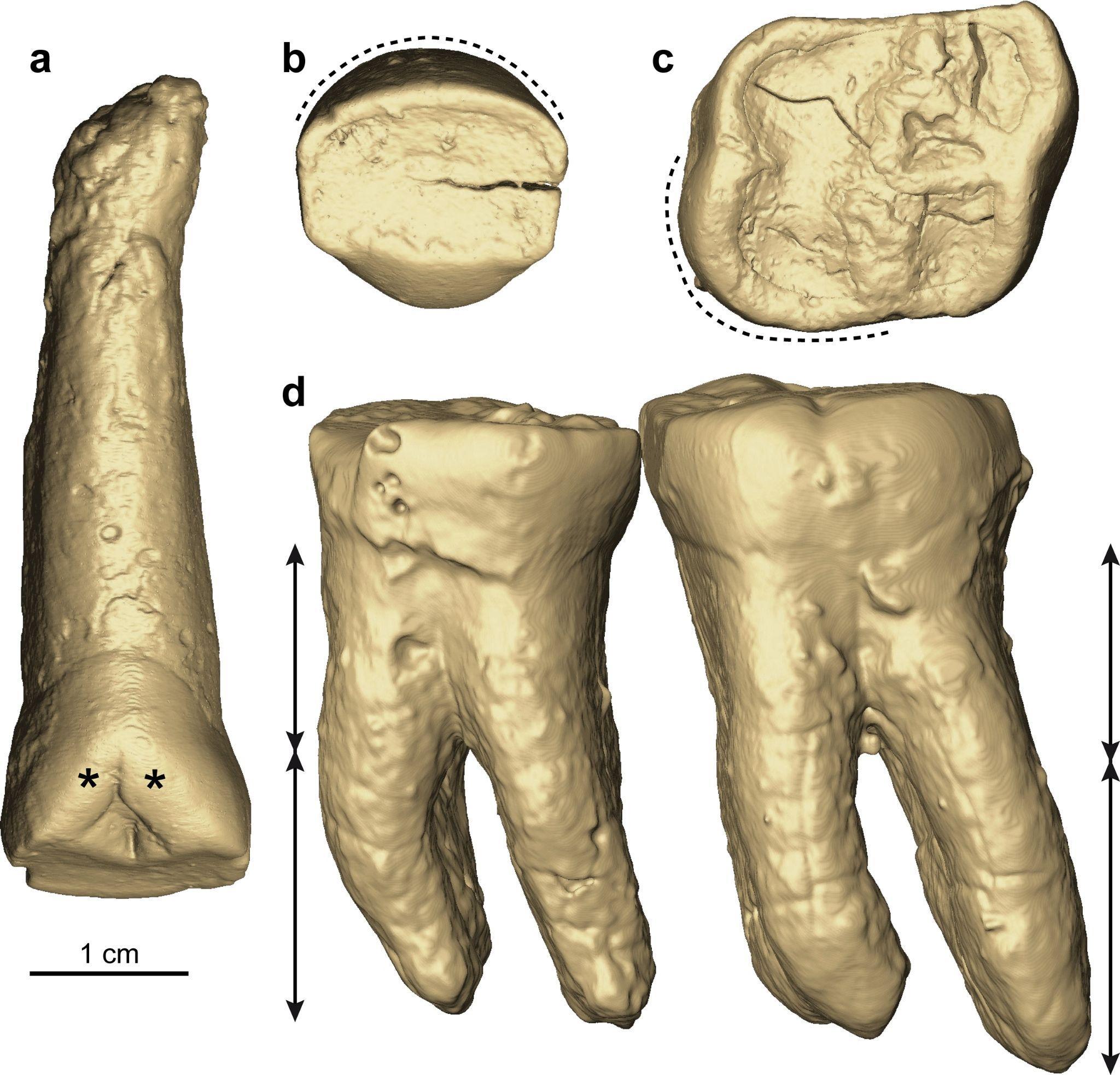

**Fig. S1. Virtual renderings of five permanent teeth from the left side of the jaw of the individual Thorin (a–d). a**, maxillary central incisor in lingual view showing developed shoveling despite the incisal wear (indicated by two asterisks), the long root and thick cement deposit (notably at the rot apex). **b**, maxillary lateral incisor illustrating the mark labial curvature (dotted line). **c**, maxillary first molar in occlusal view displaying the large hypocone (dotted line). **d**, mandibular first and second molars in buccal view showing the high root stem with respect to the branches (the arrows show the low root bifurcation).

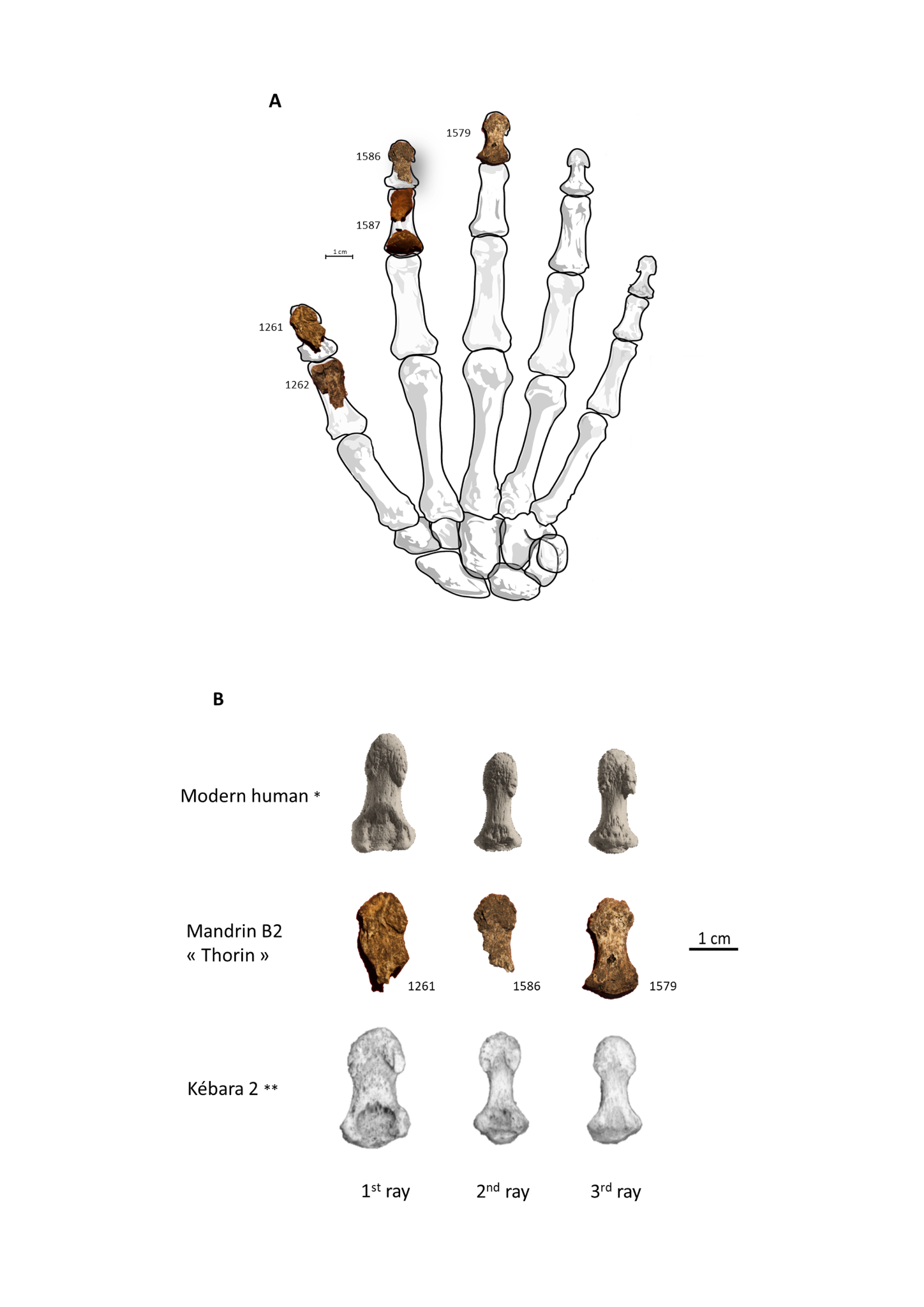

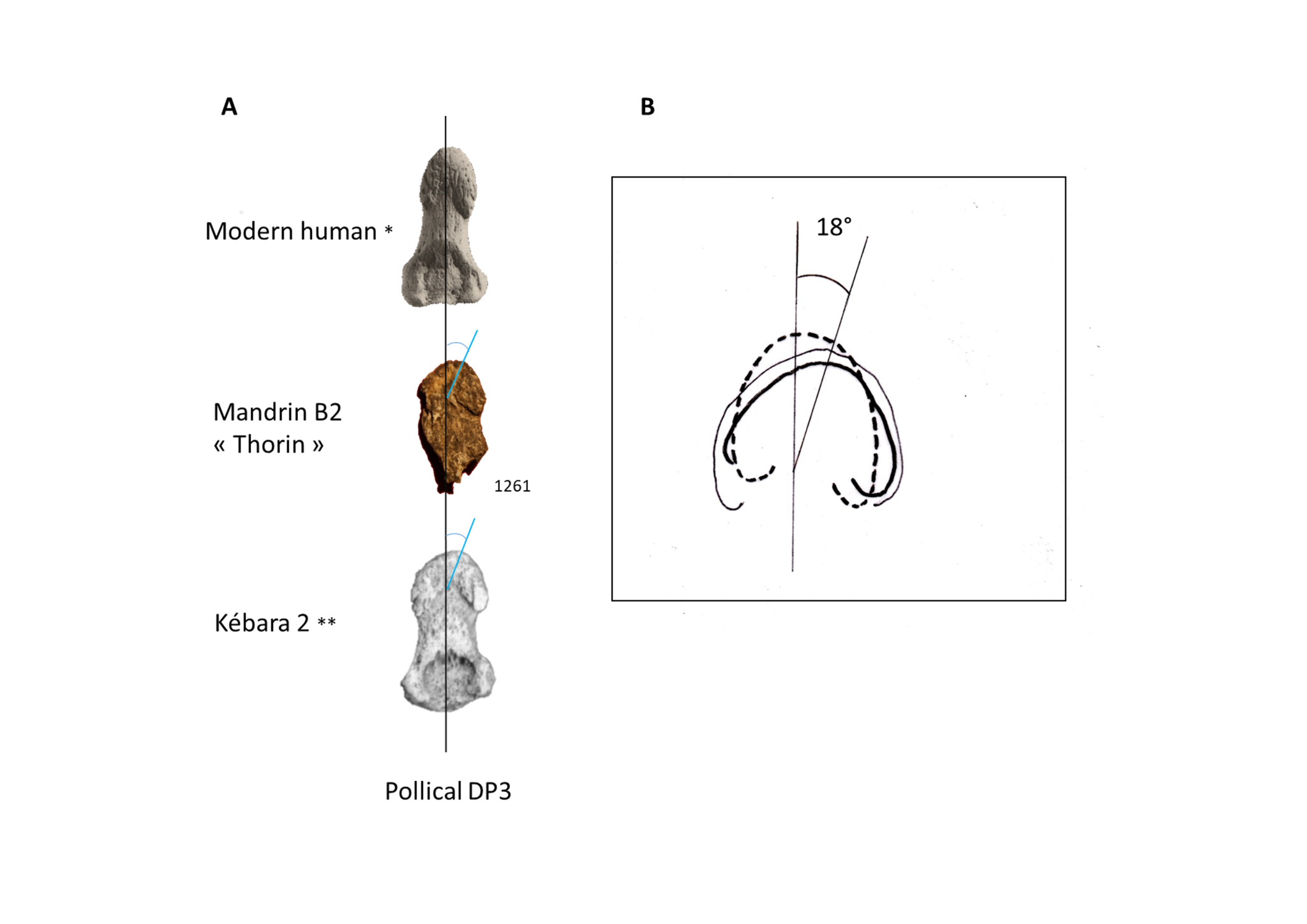

**Fig. S2.** Right. A: Setting of the skeletal elements of Thorin's left hand (palmar view). B: comparison of the left DP1, DP2, DP3 of Thorin’s hand with those of Kebara 2 Neanderthal and contemporary human. *From ref. 175, modified; **From ref. 140 , modified. Left A: Neanderthal morphological characters of Thorin’s DP3 (comparison with Neanderthal Kebara DP3 and modern human DP3). B: comparative drawing of the pollical DP3 (bold line: Thorin; thin line: Kebara 2; dotted line: modern man) showing the ulnar deviation (angle of about 18°) and the rounded morphology of the distal tuberosity of Neanderthals thumb distal phalanx.

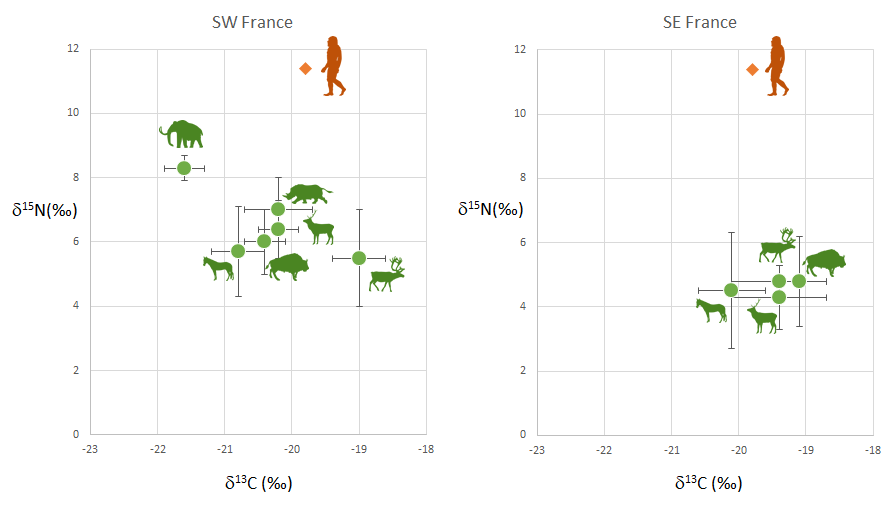

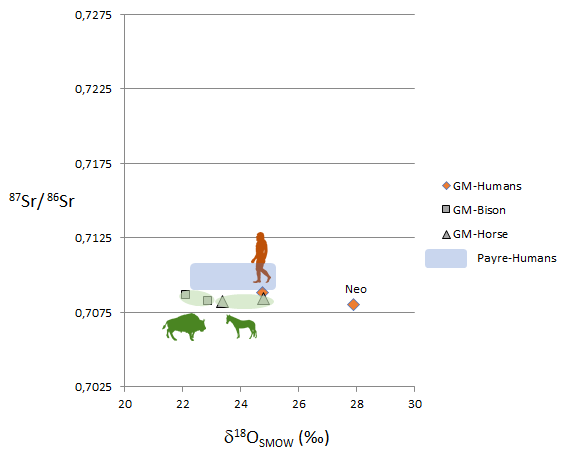

**Fig. S3 |** (S1 & S2) **Top**. *δ*^13^C and *δ*^15^N values of (Left) the late Neanderthal and penecontemporaneous fauna from St. Césaire in Southwestern France, compared with (Right) Thorin and penecontemporaneous fauna in Southeastern France. **Bottom.** ^87^Sr/^86^Sr isotopic ratios against *d*^18^O for Thorin, a Neolithic human, and horse and bison from Mandrin, and Neanderthal ranges from the nearby site of Payre.

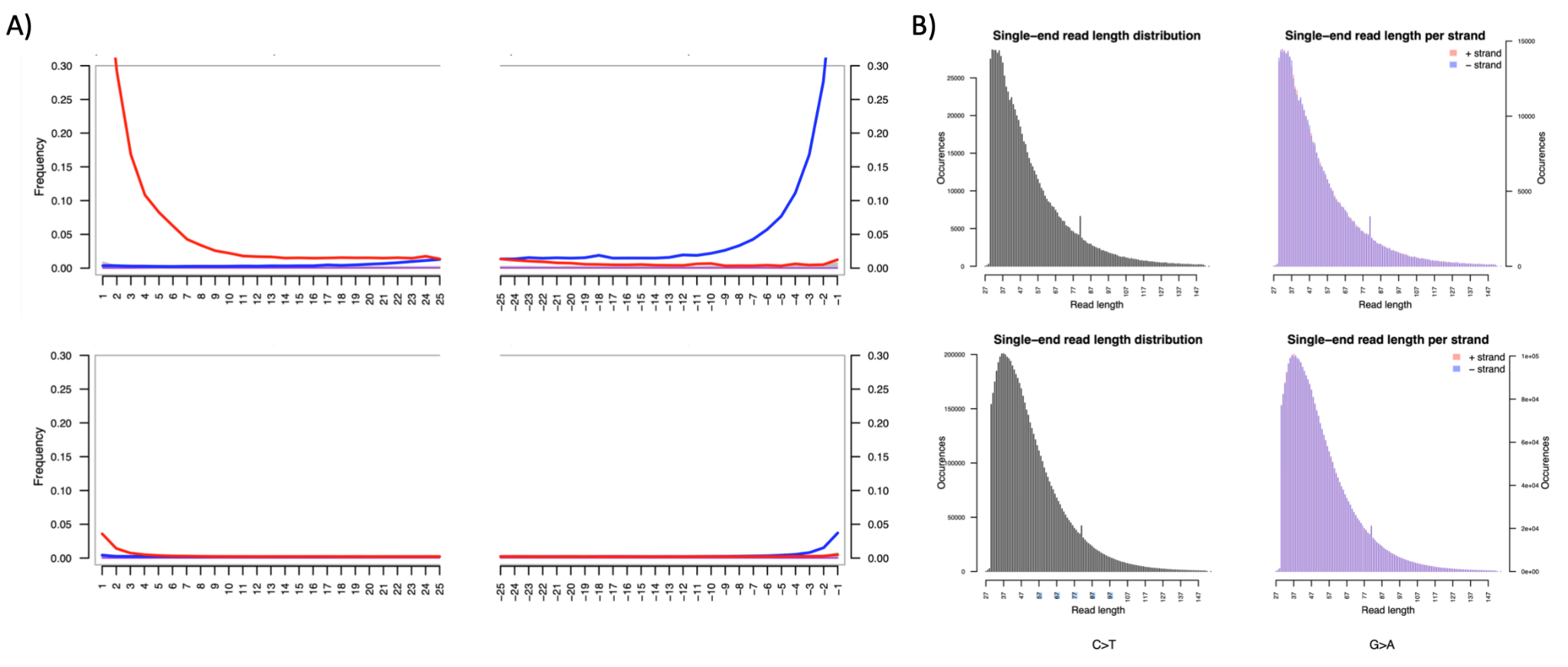

**Fig. S4. Cytosine deamination pattern and fragment length distribution.**

(A) We show the frequency of C to T (left) and G to A (right) substitutions along the strand for both a non-USER treated (upper panel) and a USER treated library (lower panel). We find that the C to T and G to A substitutions are successfully removed by the USER treatment. (B) Fragment size distributions for a non-USER treated (upper panel) and a USER treated library (lower panel).

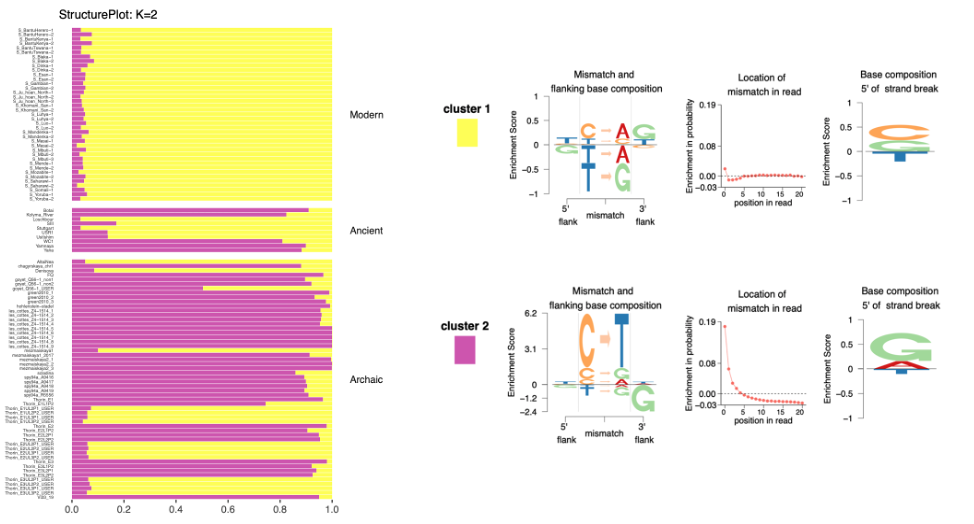

**Fig. S5. Structure plot and mismatch profiles for *K*=2.**

In A) we show the results of aRchaic using two clusters. We used modern and ancient samples as reference in order to assess the damage patterns in the archaic samples/libraries. The test using two clusters distinguishes between modern and ancient non-USER treated samples. The modern samples show a clear membership in cluster 1, while non-USER treated libraries are assigned to cluster 2. Since the distinct deamination patterns in the fragment termini are removed in USER-treated libraries, these display high membership in cluster 1 as well. B) Displays the mismatch profiles of the two clusters.

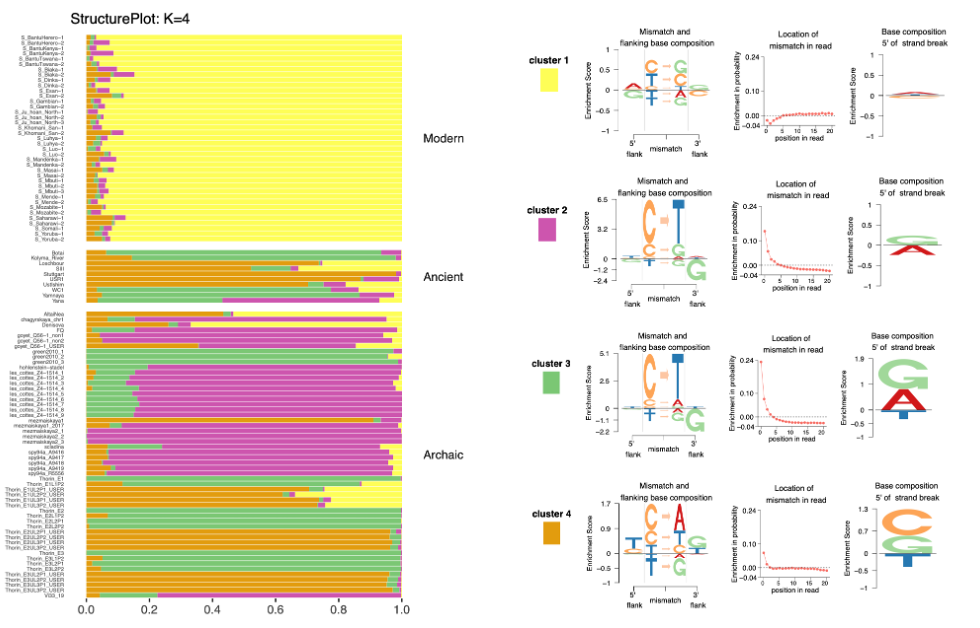

**Fig. S6. Structure plot and mismatch profiles for *K*=4.**

In A) we show the structure plot of the same set of samples shown in Fig. S4. For this test, we assess whether it is possible to distinguish between USER and non-USER treated samples and libraries. The modern samples show a high level of membership in cluster 1, while non-USER treated ancient and archaic samples are assigned to cluster 3. USER-treated samples and libraries show elevated membership in cluster 4. B) Mismatch profiles of the four clusters.

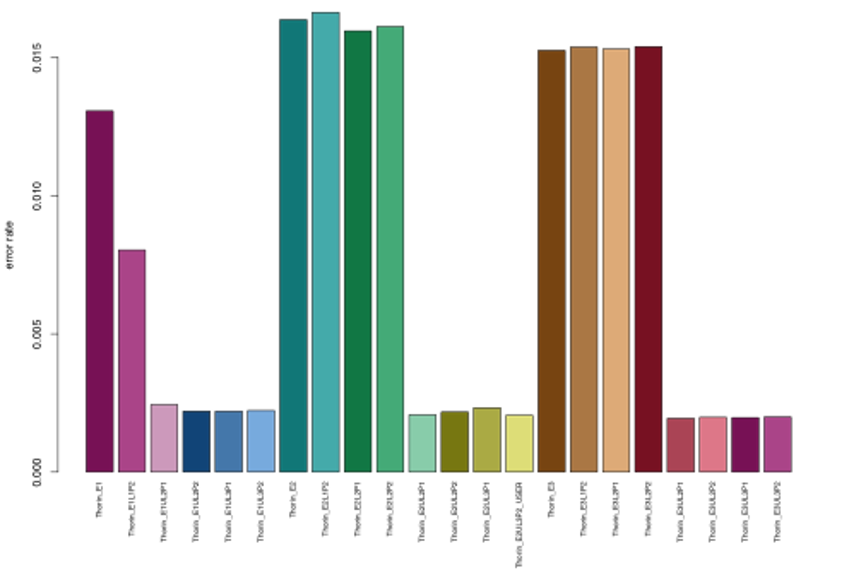

**Fig. S7. Overall error rate for both USER-treated and non-treated libraries.**

Error rates for each of the 22 libraries generated from the Thorin sample.

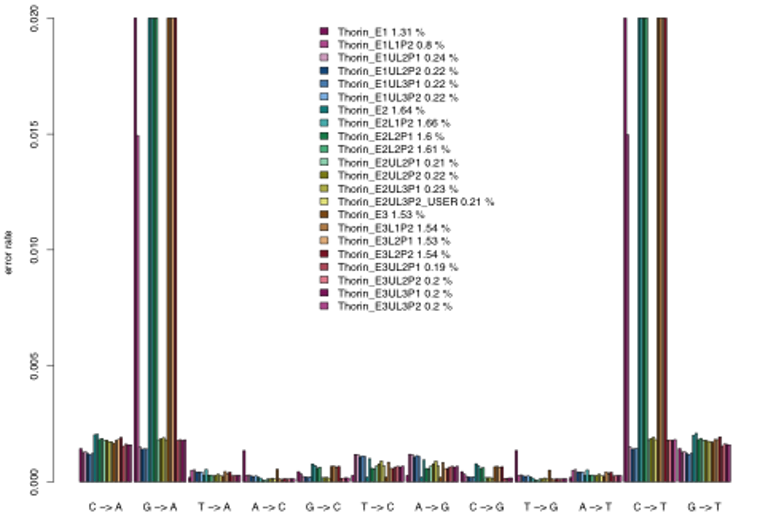

**Fig. S8. Type specific error rates per library.**

The graph shows the rates of which each of the 12 possible substitutions are observed in the data. The percentages assigned next to each library denotes the overall error rate.

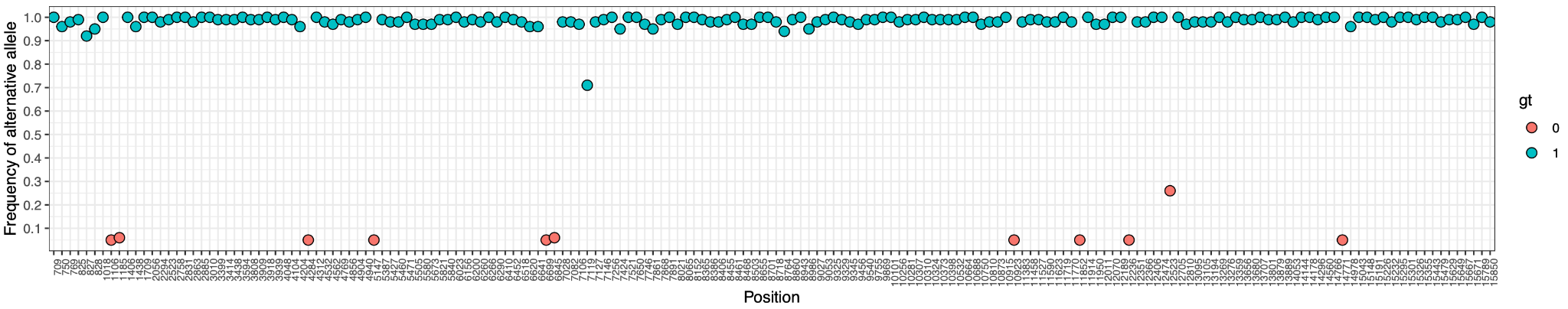

**Fig. S9. Identification of ambiguous sites in the mitochondrial genome.**

Plot shows the frequencies of alternative alleles across the mitochondrial genome. The x-axis indicates genomic position, while the y-axis indicates the fraction of reads carrying the alternative allele at the specific position. Colors indicate the called genotype at each position. Only sites where at least 5% of the reads carried the alternative allele are displayed.

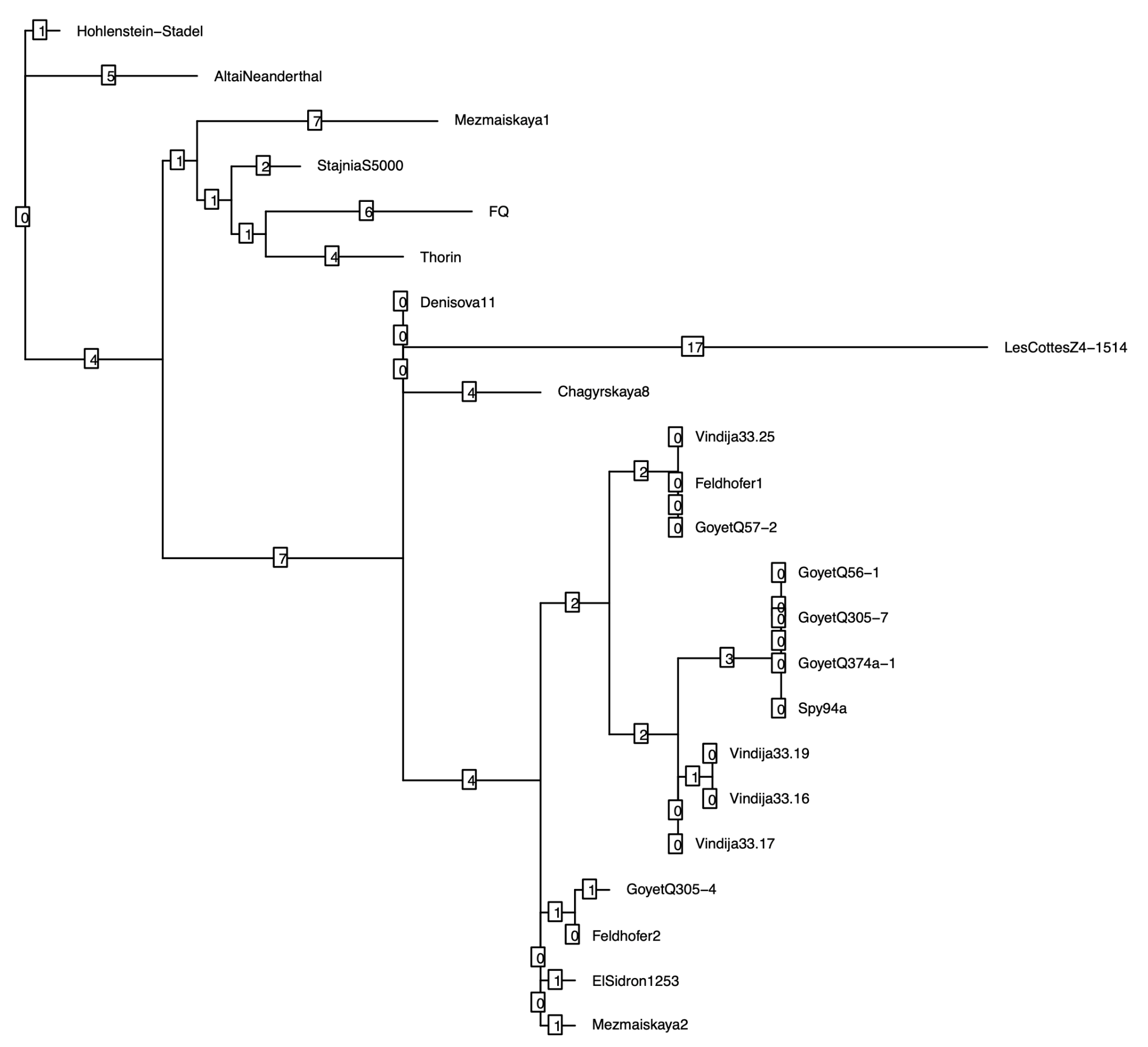

**Fig. S10. Maximum parsimony analysis.**

Phylogenetic tree from maximum parsimony analysis using 23 Neanderthals and Denisova 3 as an outgroup (not included in tree). The substitution counts are denoted in the squares of each branch.

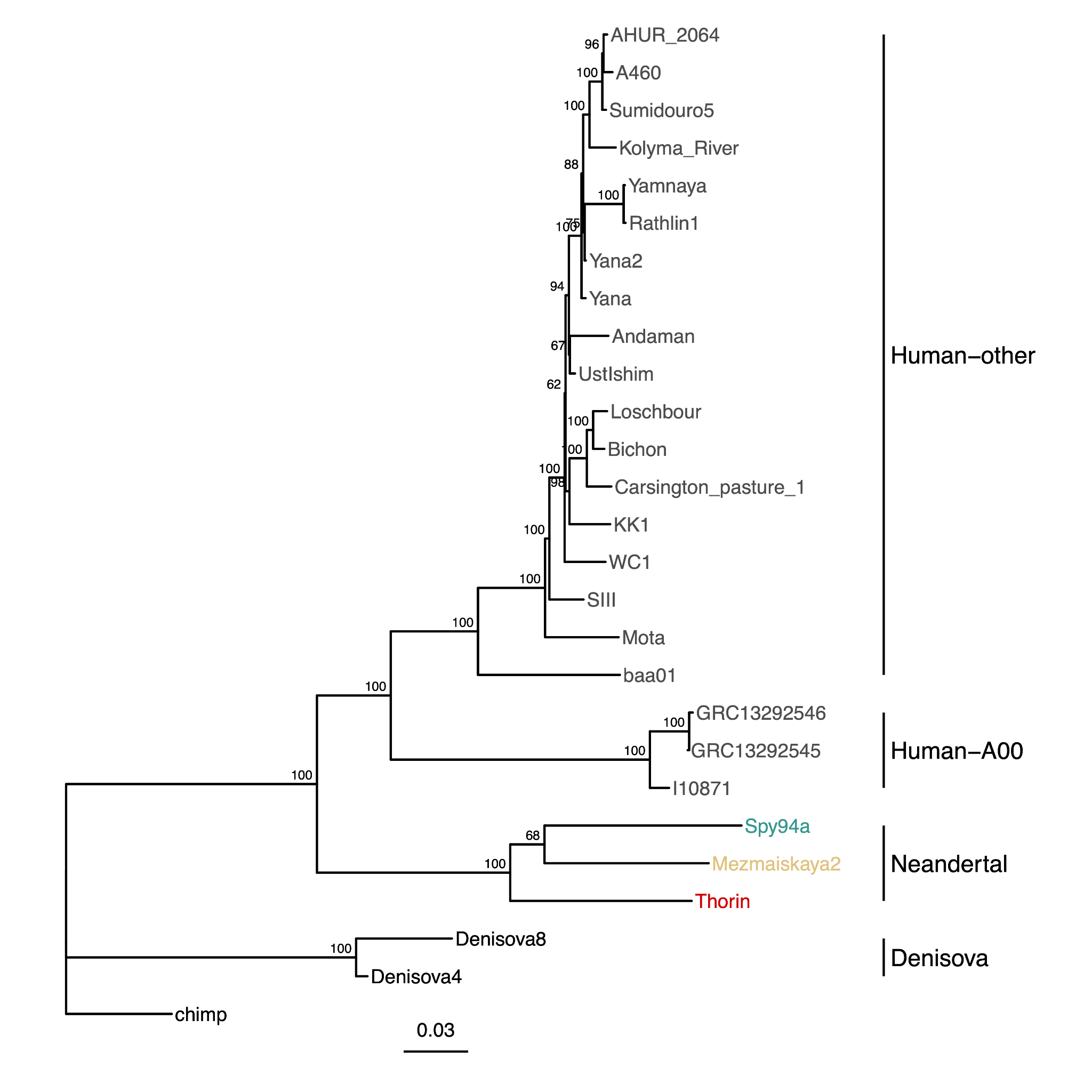

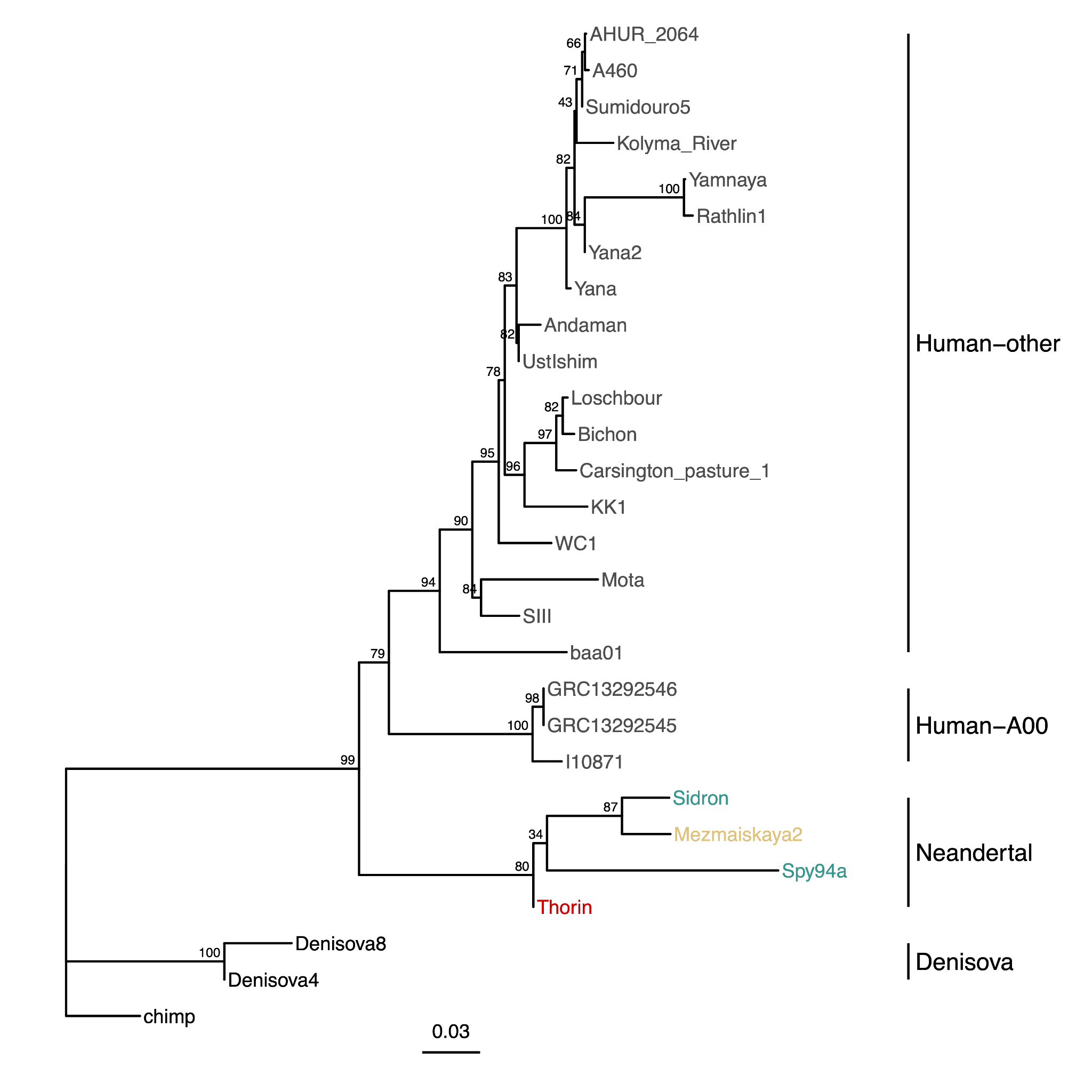

**Fig. S11.** Neighbor-joining tree of archaic and modern human Y chromosome sequences. Numbers at branches represent bootstrap support based on 100 replicates. **Top,** based on transversion SNP genotypes within the 6.9 Mb region targeted by the capture probes used to generate data for the Spy94a and Mezmaiskaya2 samples. **Bottom,** based on transversion SNP genotypes within the 560 kb region targeted by the capture probes used to generate data for the El Sidrón 12532 sample.

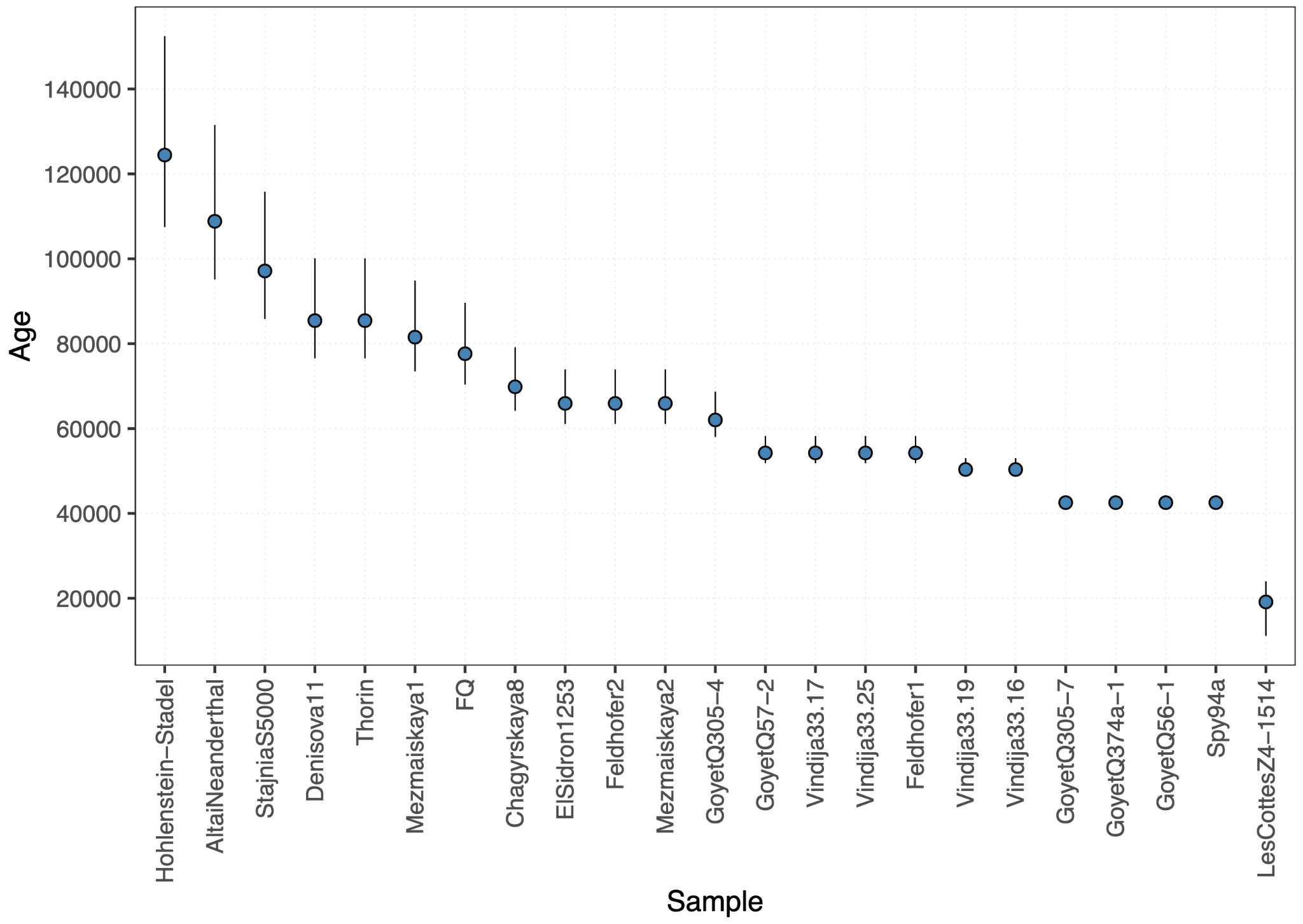

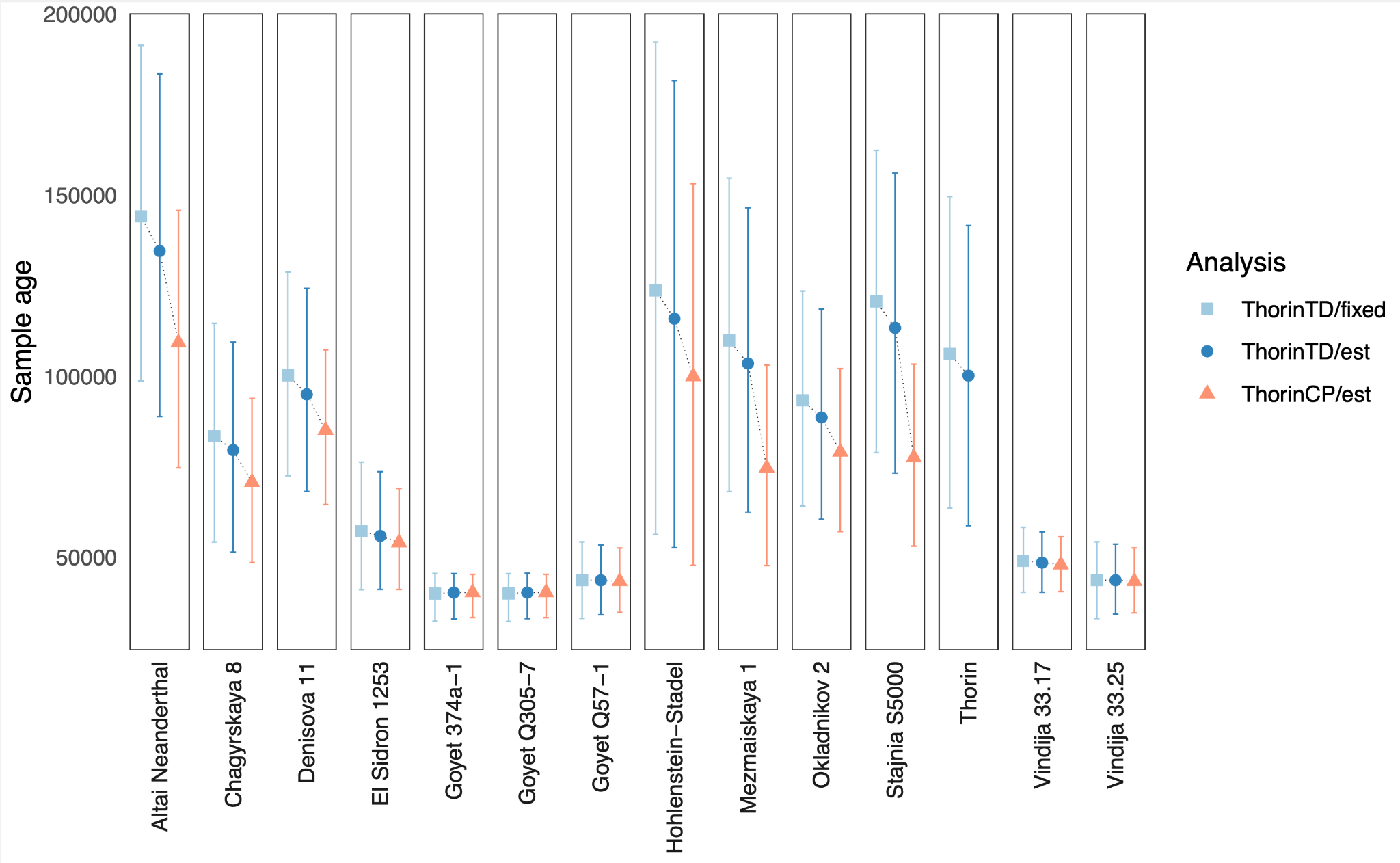

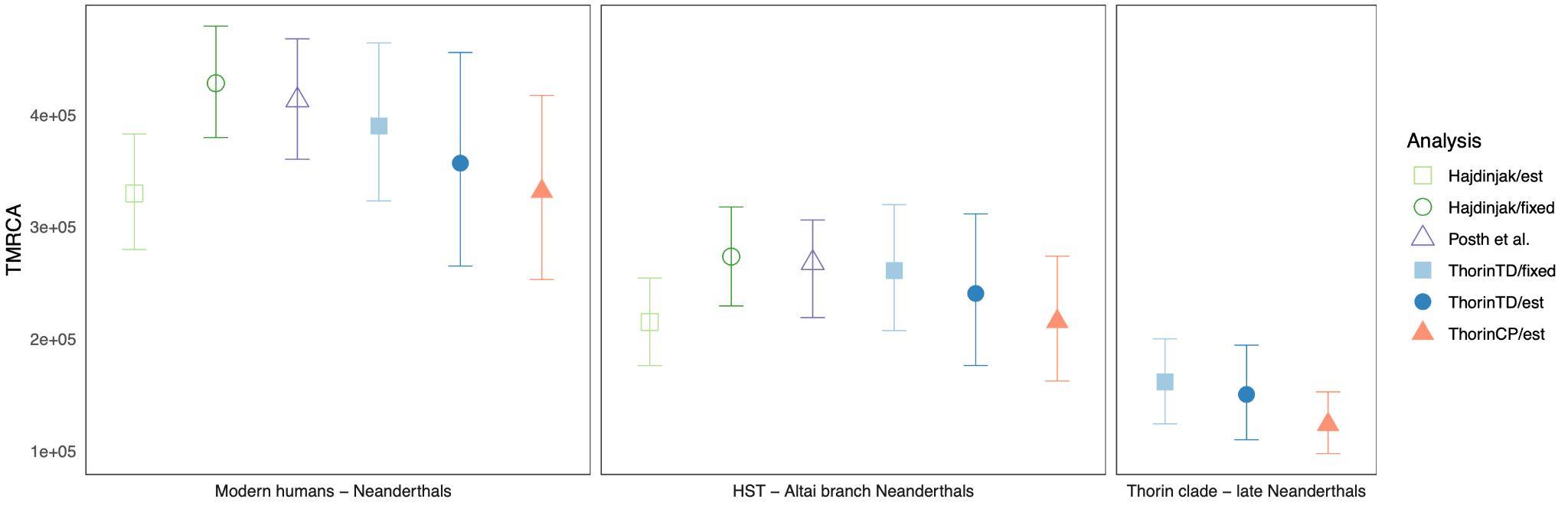

**Fig. S12.** Points indicate the mean ages, while error bars indicate the 95% confidence intervals (or HPD) for each sample. **Top, Relative age estimates obtained by branch shortening.** Age estimates obtained by branch shortening using GoyetQ56-1 as a calibration point. **Middle Comparison of age estimates obtained from BEAST analysis.** Blue square = Thorin Tip Dated (i.e., analysis where Thorin was not used as calibration point) with fixed substitution rate, blue circle = Thorin Tip Dated with estimated substitution rate, red triangle = Thorin Calibration Point (analysis using Thorin as an additional calibration point) with estimated substitution rate. **Bottom, Comparison of TMRCAs from this study with previous studies**^16 ,107^ . ThorinTD = Thorin Tip Dated (i.e., analysis where Thorin was not used as calibration point), ThorinCP = Thorin Calibration Point (analysis using Thorin as additional calibration point). Est = estimated substitution rate, fixed = fixed substitution rate.

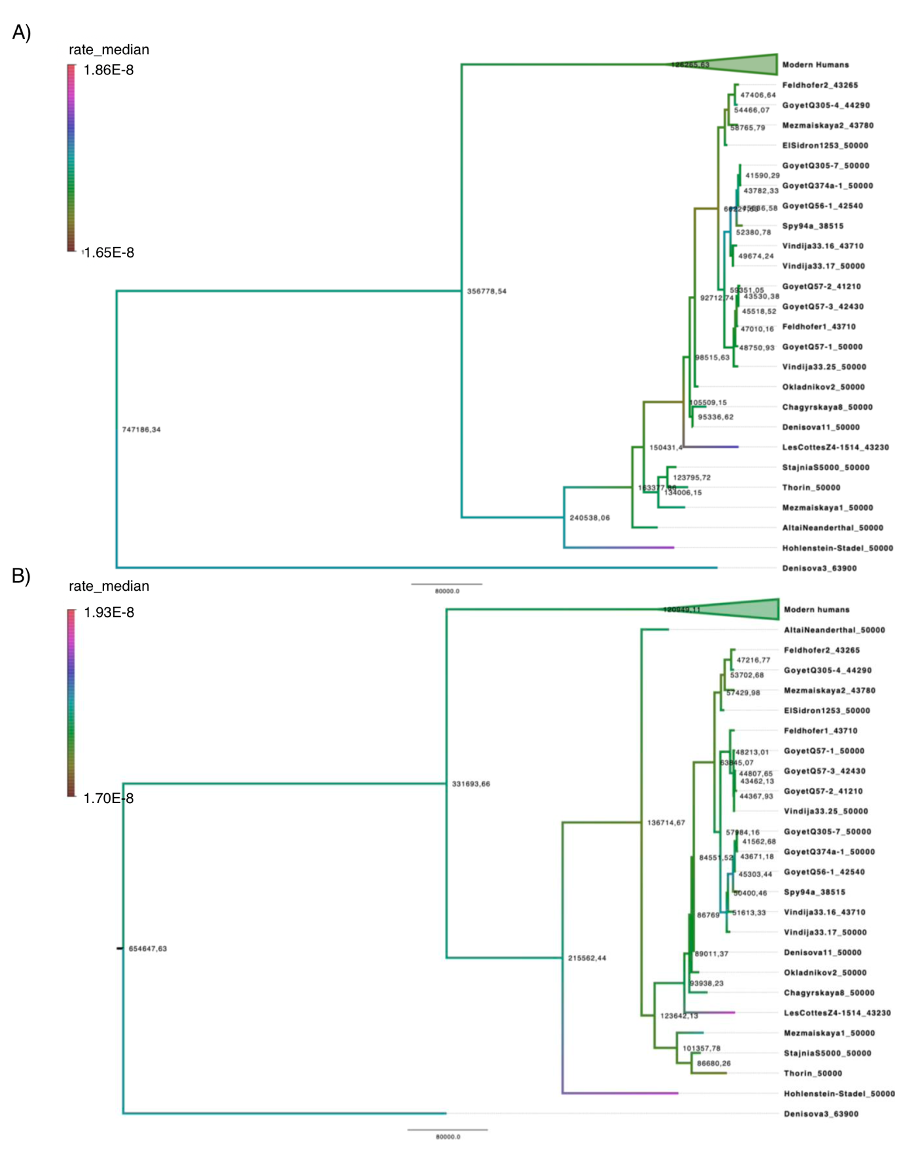

**Fig. S13. Phylogenetic trees obtained with BEAST.** A) Tree phylogeny from analysis, where Thorin was tip dated. B) Tree phylogeny from analysis with Thorin included as additional calibration point. Branches are colored by the median substitution rate. Node labels indicate mean divergence dates.

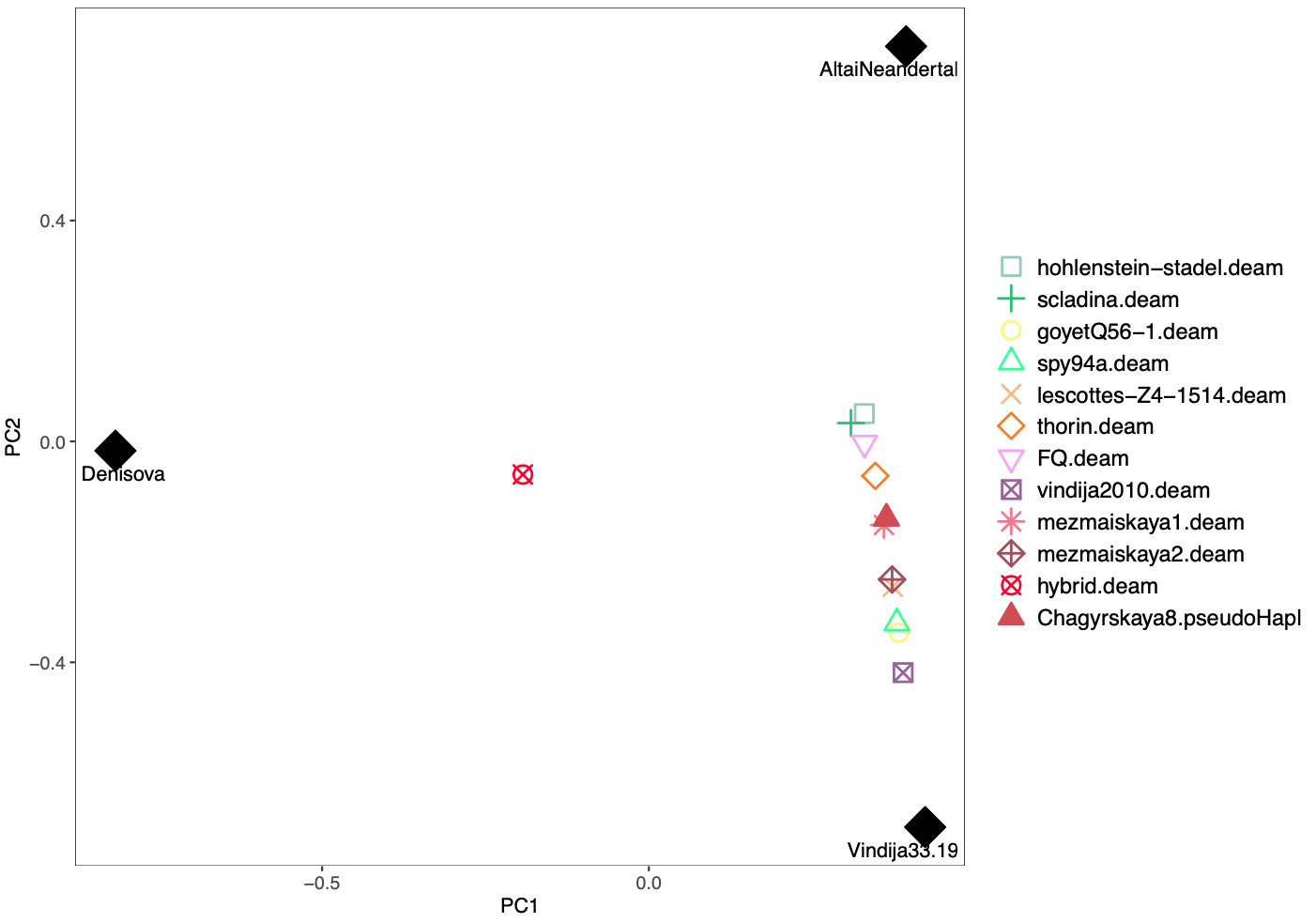

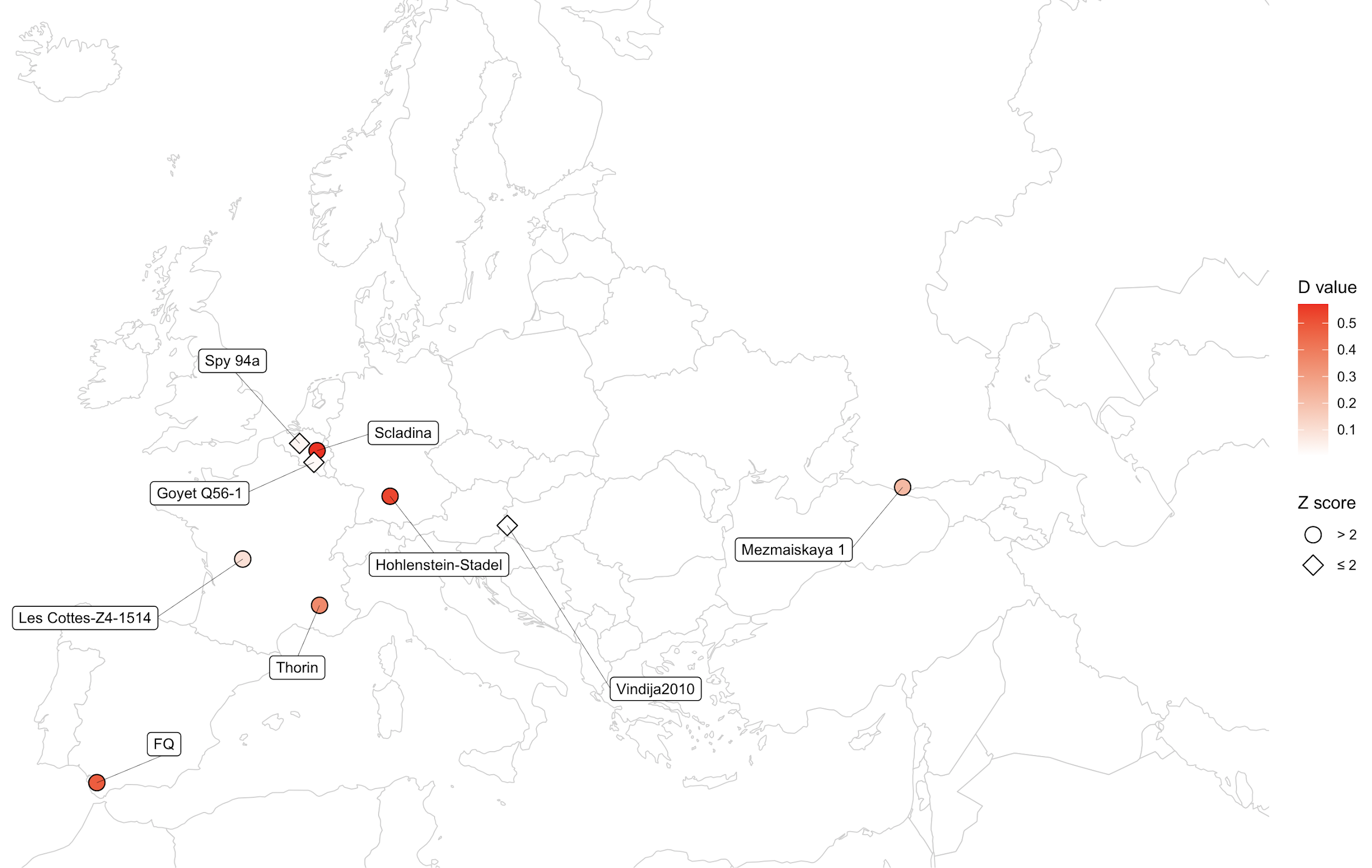

**Fig. S14. Top, Procrustes analysis of low coverage Neanderthals.** Procrustes analysis based on coordinates obtained from PCA. The filled points indicate high coverage samples, while black filled shapes indicate the samples used for the reference shape in the procrustes transformation. The analyzed samples are individually projected onto the three reference samples and visualized together. **Bottom,** Map showing geographic location and D statistics in the form of D (Vindija 33.19, Neanderthal X; Mezmaiskaya 2, Mbuti) for the Thorin Neanderthal and other Neanderthal samples used in this study. Circle: Z-score above 2 meaning that the tested tree topology is rejected, rhombus: Z-score equal to or under 2 meaning that the tested tree topology is accepted.

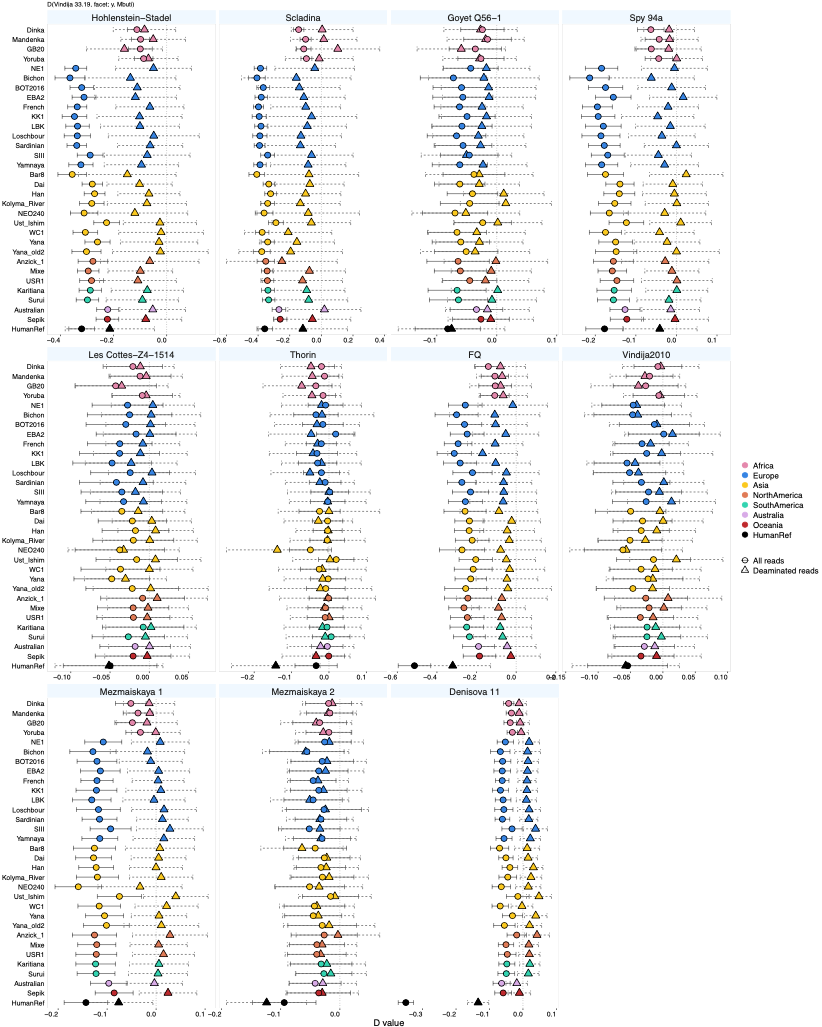

**Fig. S15.** D-statistics in the form of D (Vindija 33.19, Neanderthal X; modern human, Mbuti) to test for tree-like relationship between Vindija 33.19 and Neanderthal X with respect to modern humans. The estimated D values are visualized along the X-axis, while the tested contemporary populations are shown along the Y-axis. Circle represents analysis using all fragments parsing initial filtering, while triangle represents results using only deaminated reads. Error bars indicate 3 standard error, while the number beside each point indicates the Z-score. D-statistics obtained with all fragments are shifted from the D values obtained using only deaminated reads for most Neanderthals. The observed shift is an indicator of contamination from a non-African source. We do not observe a notable shift in Thorin suggesting limited amounts of contamination in our data. Based on the results from the damaged reads Thorin does not show evidence of additional allele sharing with Modern Humans.

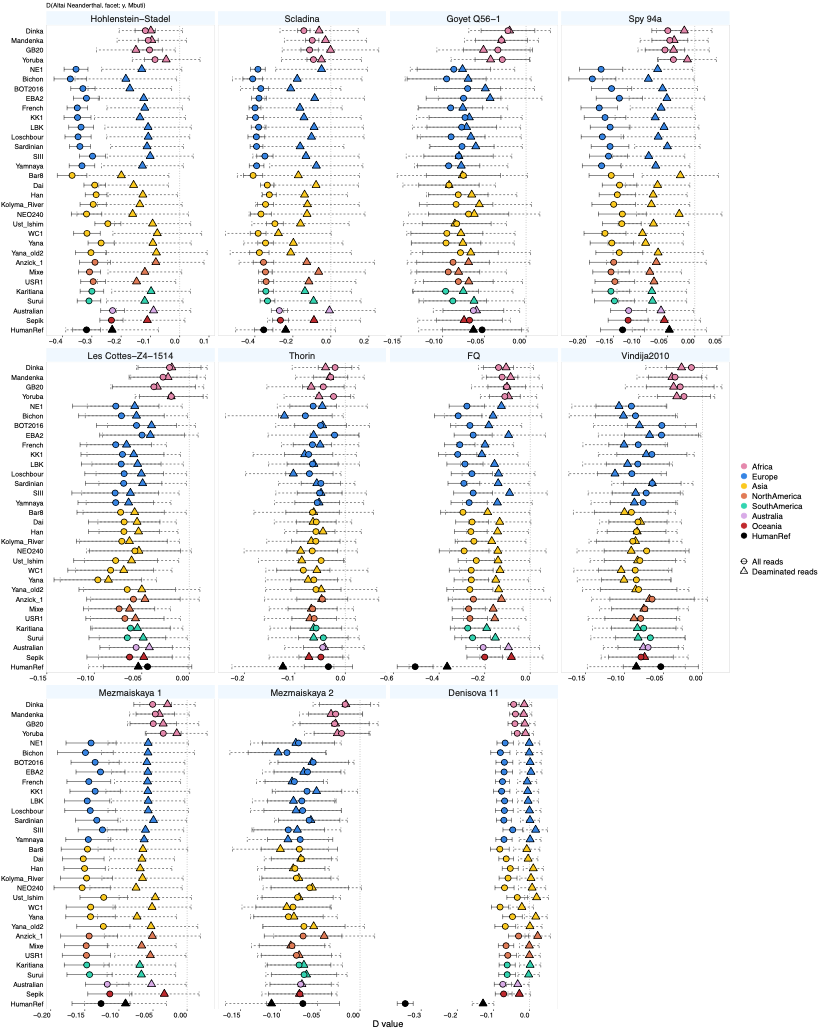

**Fig. S16.** D-statistics in the form of D (Altai Neanderthal X; modern human, Mbuti) to test for tree-like relationship between the Altai Neanderthal and Neanderthal X with respect to modern humans. The estimated D values are visualized along the X-axis, while the tested contemporary populations are shown along the Y-axis. Circle represents analysis using all fragments parsing initial filtering, while triangle represents results using only deaminated reads. Error bars indicate 3 standard error, while the number beside each point indicates the Z-score. D-statistics obtained with all fragments are shifted from the D values obtained using only deaminated reads for most Neanderthals. The observed shift is an indicator of contamination from a non-African source. D-values obtained from damaged reads are shifted from D=0 for non-African populations due to excess allele sharing between Neanderthal X and the human-introgressing Neanderthal ancestry carried by Modern Humans.

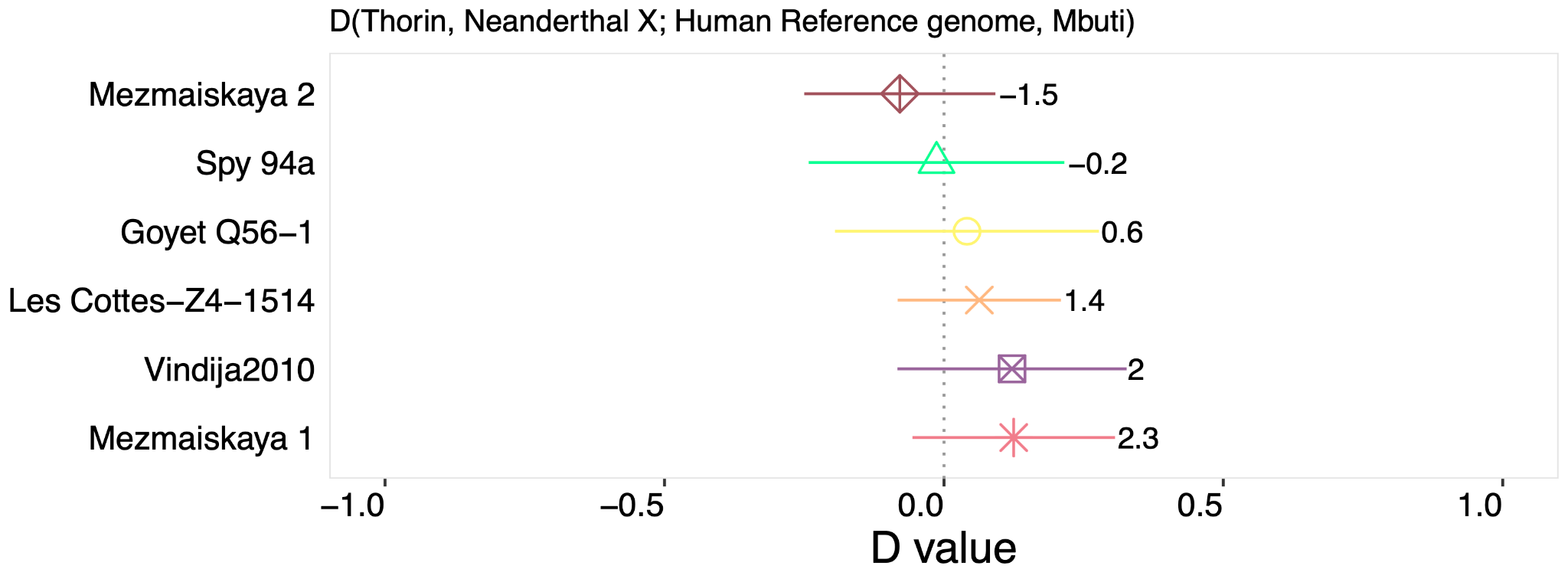

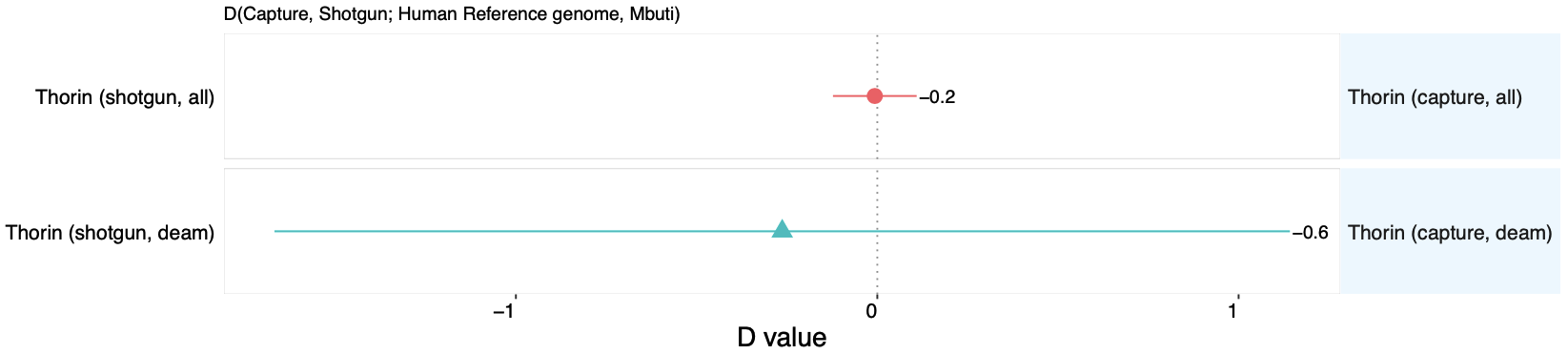

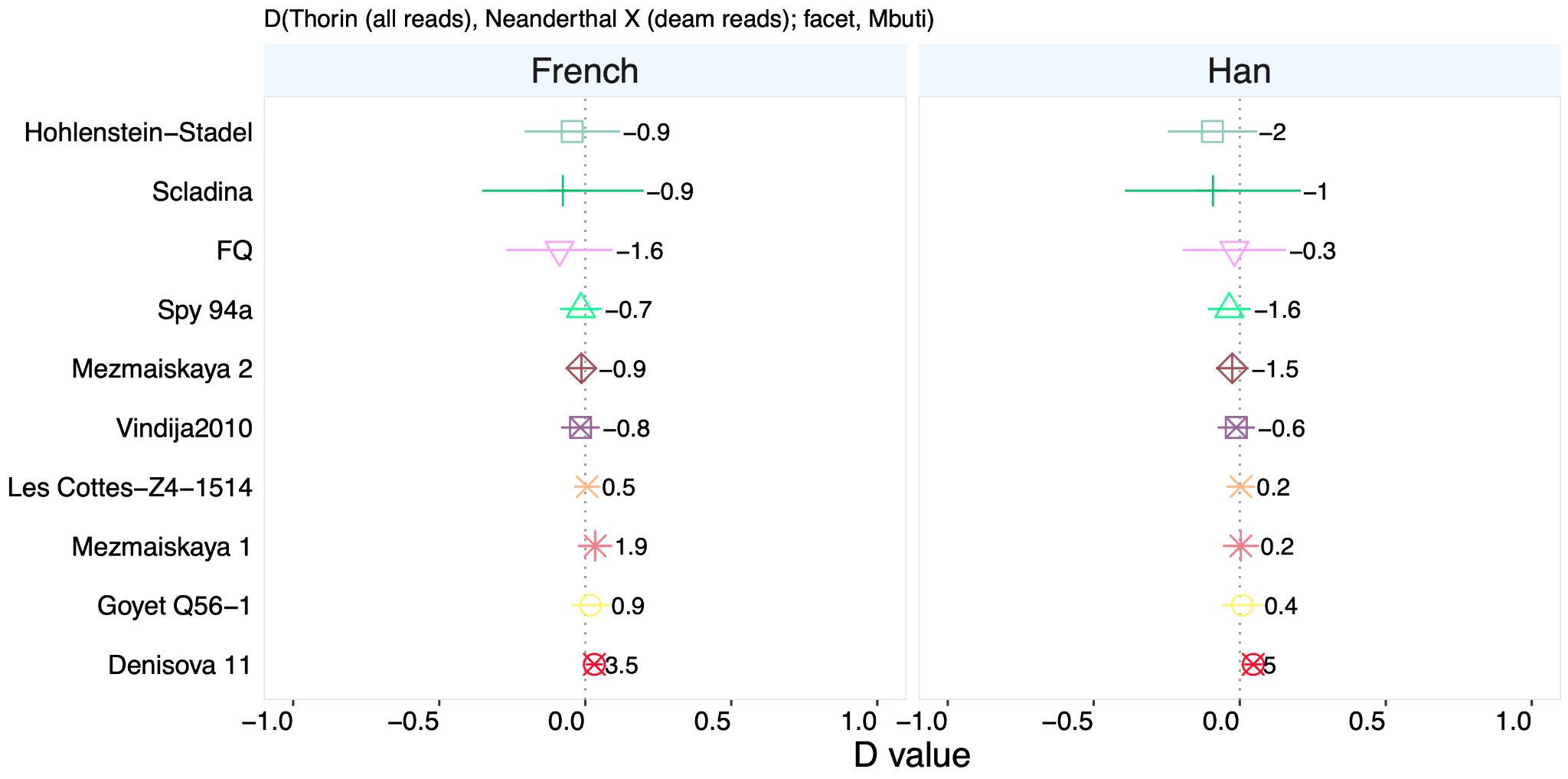

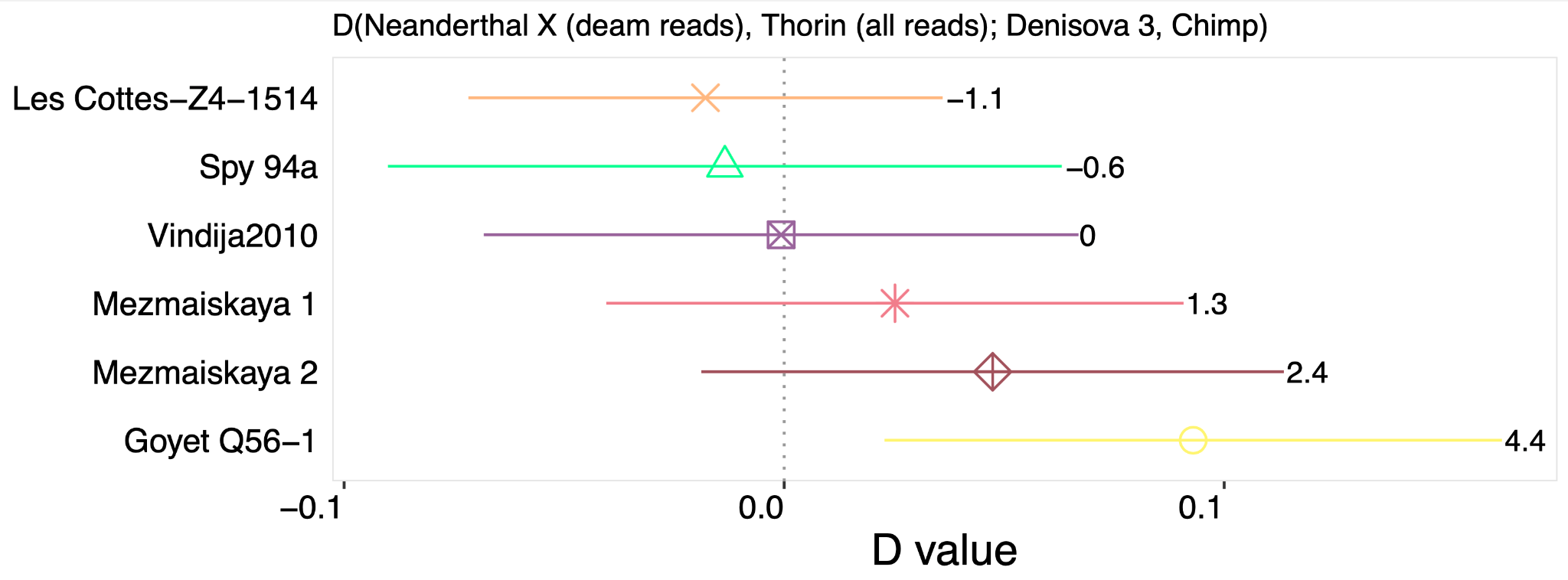

**Fig. S17. D-statistics in the form of D.** D-values are plotted along X-axis, and tested Neanderthals are shown along Y-axis. **Top,** D (Thorin, Neanderthal X; Human Reference Genome, Mbuti) using damaged reads to test for excess allele sharing between Thorin and the reference genome with respect to Neanderthal X. In cases of more reference bias in Thorin than Neanderthal X, we would expect to observe a significant departure from D = 0, i.e. Z > 3. We do not observe a significant signal when comparing Thorin to previously published low coverage Neanderthals, thus we exclude the possibility of reference bias influencing our downstream analysis**. Middle,** D (Thorin (all reads), Neanderthal X (damaged reads), French/Han, Mbuti) testing for recent gene flow between Thorin and Modern Humans. We do not find Thorin carrying additional alleles shared with contemporary populations, confirming that Thorin and the other tested Neanderthal are forming a clade with respect to the human-introgressing Neanderthal. **Bottom,** D (Neanderthal X (damaged reads), Thorin (all reads); Denisova 3, Chimp) to test for excess allele sharing between Thorin and Denisova 3 compared to other low coverage Neanderthals. Thorin is forming a clade with each of the tested Neanderthals with respect to Denisova 3 confirming that Thorin is a Neanderthal with no additional Denisovan ancestry.

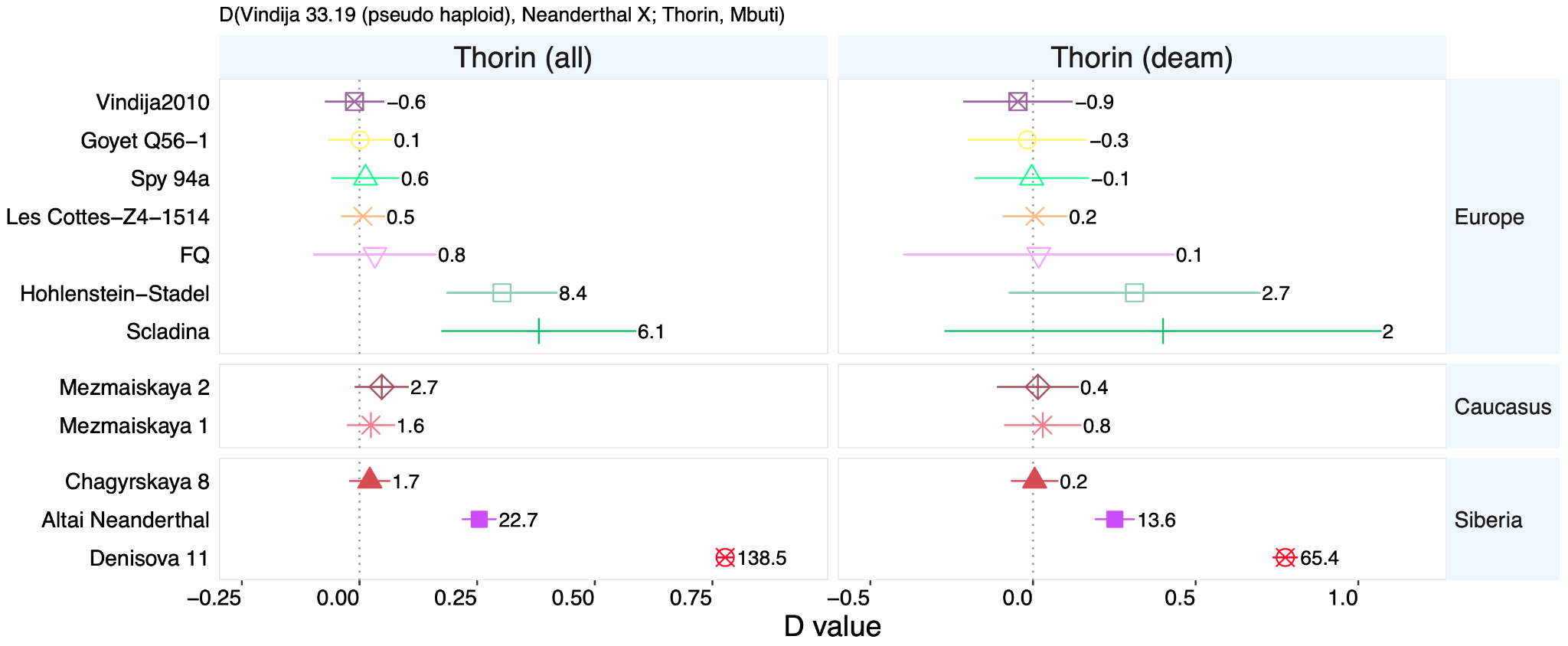

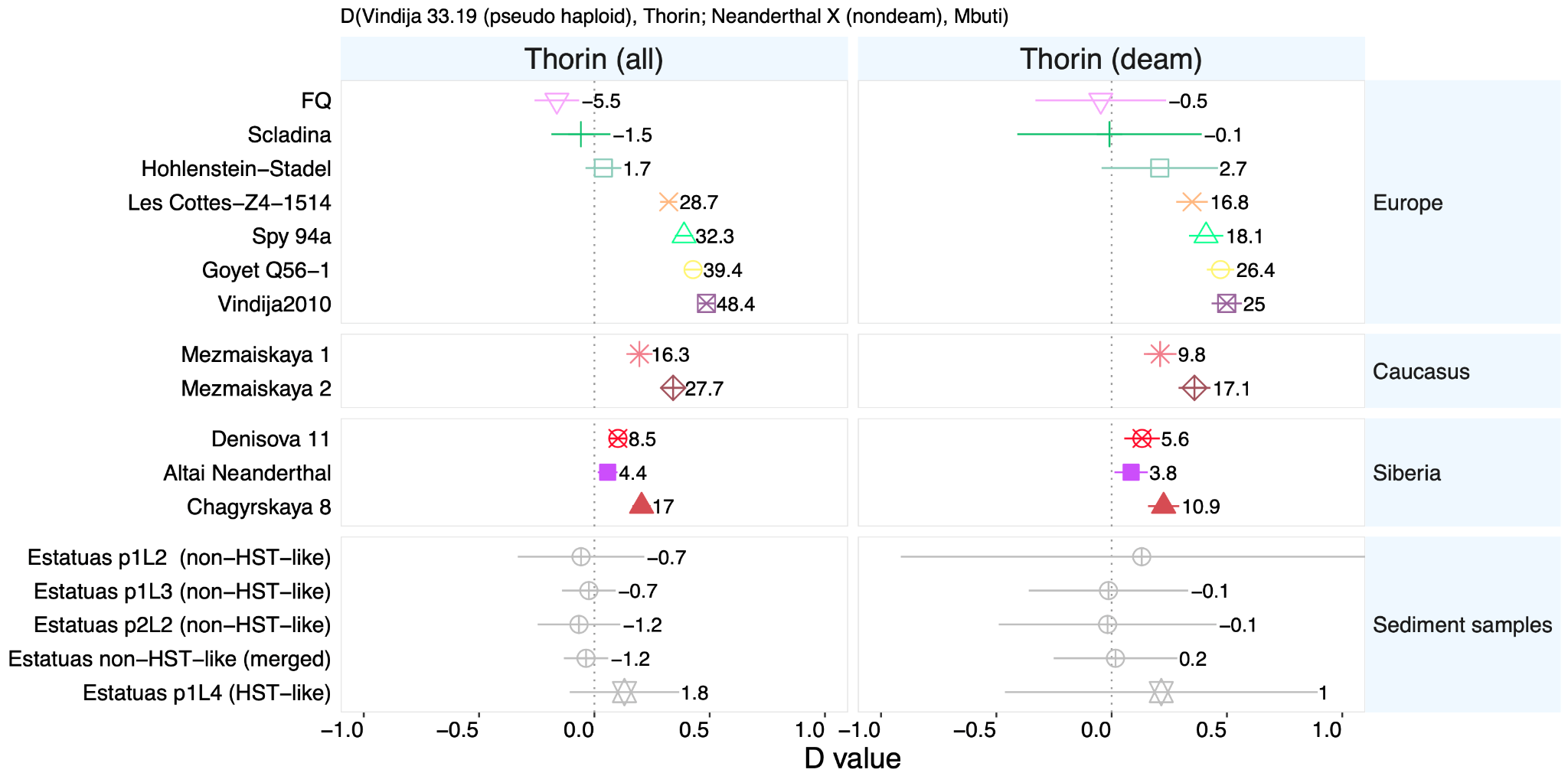

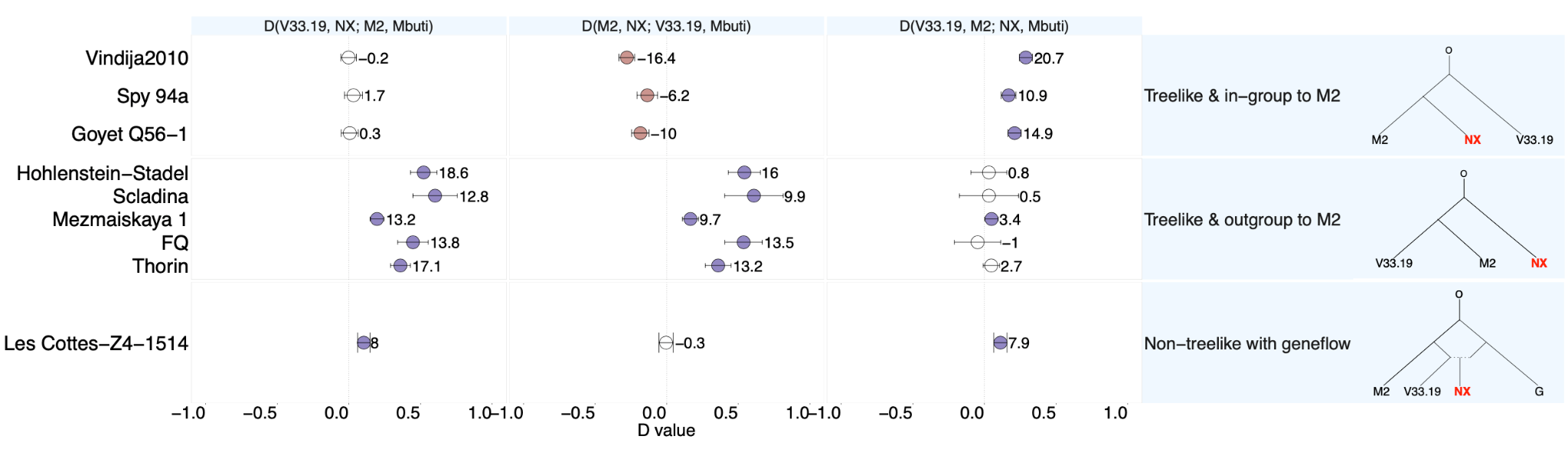

**Fig. S18. D-statistics in the form of D.** The estimated D values are visualized along the X-axis, while the test populations are shown along the Y-axis. We only use the deaminated reads for the in-group Neanderthal to avoid bias. Error bars indicate 3 standard errors, while the number beside each point indicates the Z-score. Only results using deaminated reads for Neanderthals X are shown. **Top, D (Vindija 33.19, Neanderthal X; Thorin, Mbuti)**. Thorin is an outgroup to Neanderthal samples more recent than 80 ka. **Middle, Shared drift between Forbes Quarry and Thorin.** D (Vindija 33.19, Thorin; Neanderthal X, Mbuti), for which we used pseudo haploid genotype calling for both Vindija 33.19 and Thorin in order to avoid bias. **Bottom, Evidence of gene flow in Les Cottés-Z4 1514.** D statistics in the form of D (Vindija 33.19, Neanderthal X; Mezmaiskaya 2, Mbuti).

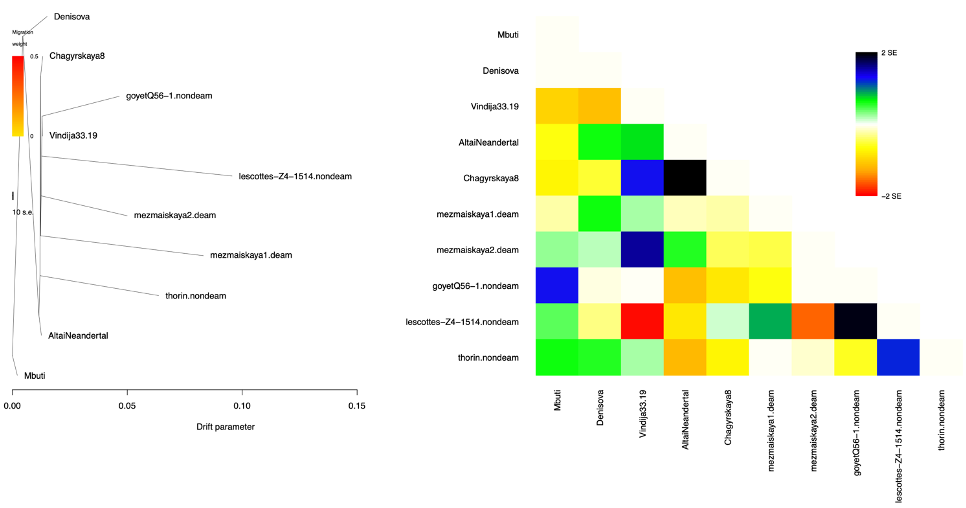

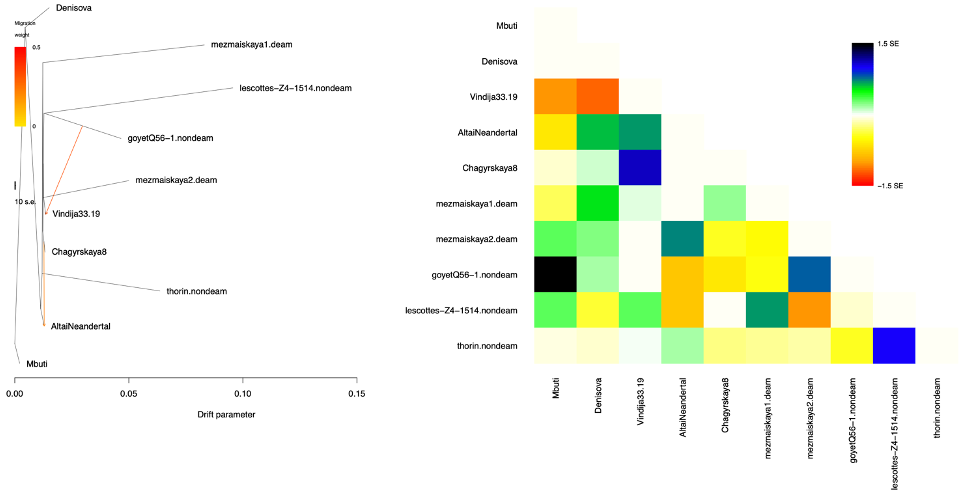

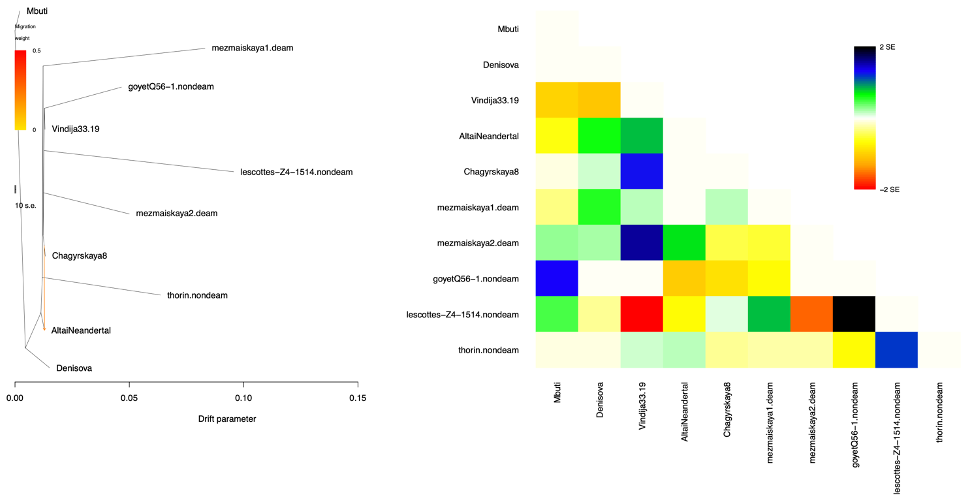

**Fig. S19. Treemix.** Left: Inferred treemix topology, with drift parameter along the X-axis. Right: Heat plot of residual values corresponding to the tree topology. **Top,** with no migration. **Middle,** with 1 migration event**. Bottom,** with 2 migration events.

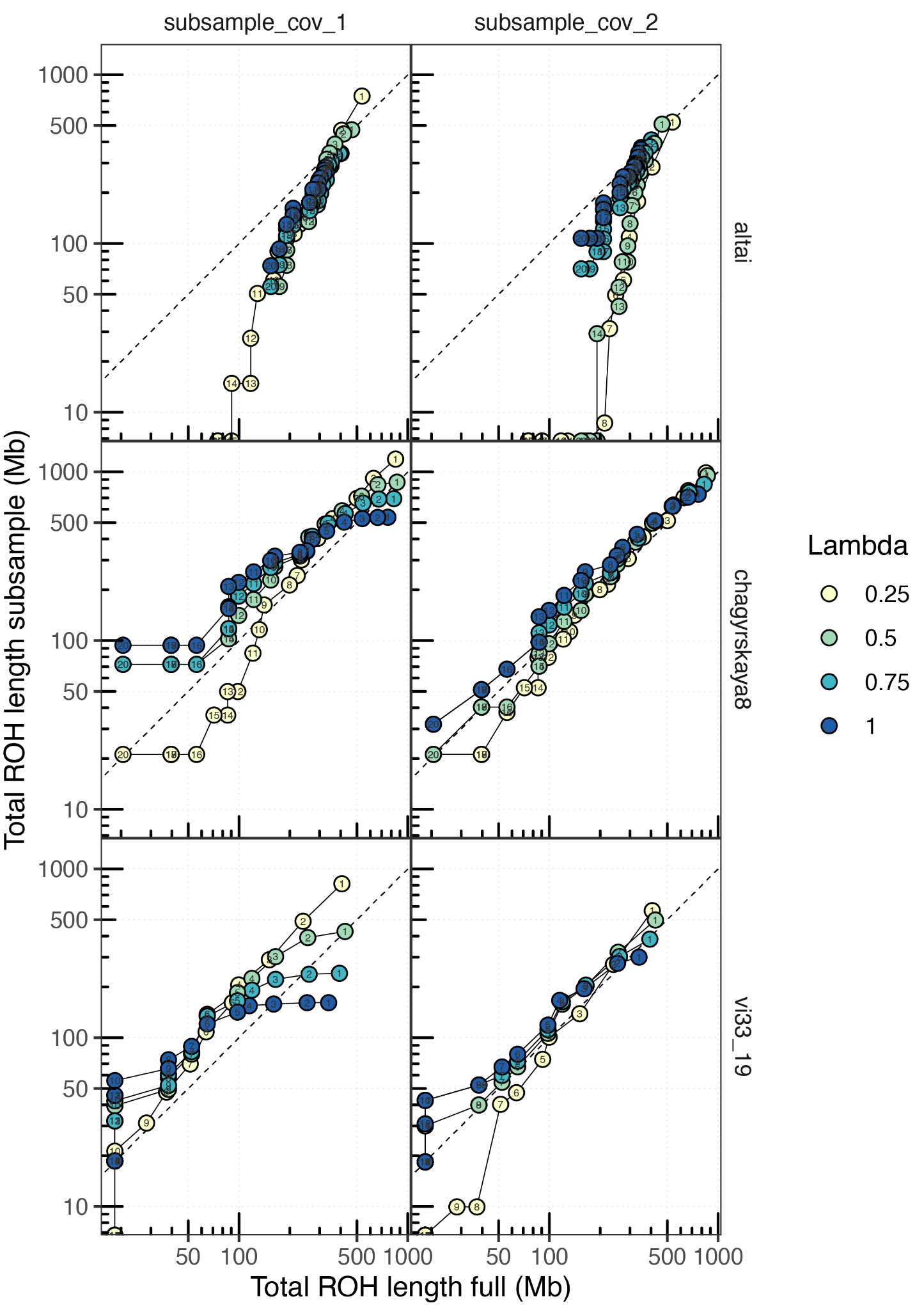

**Fig. S20 ROH calling on subsampled genomes.** Panels compare cumulative ROH lengths between subsampled and full genomes, for three high coverage Neanderthals (rows) and two different coverage values (columns). Each panel shows cumulative ROH lengths for four different values of regularization parameter λ (indicated by symbol color), with symbol numbers indicating minimum ROH length cutoff used.

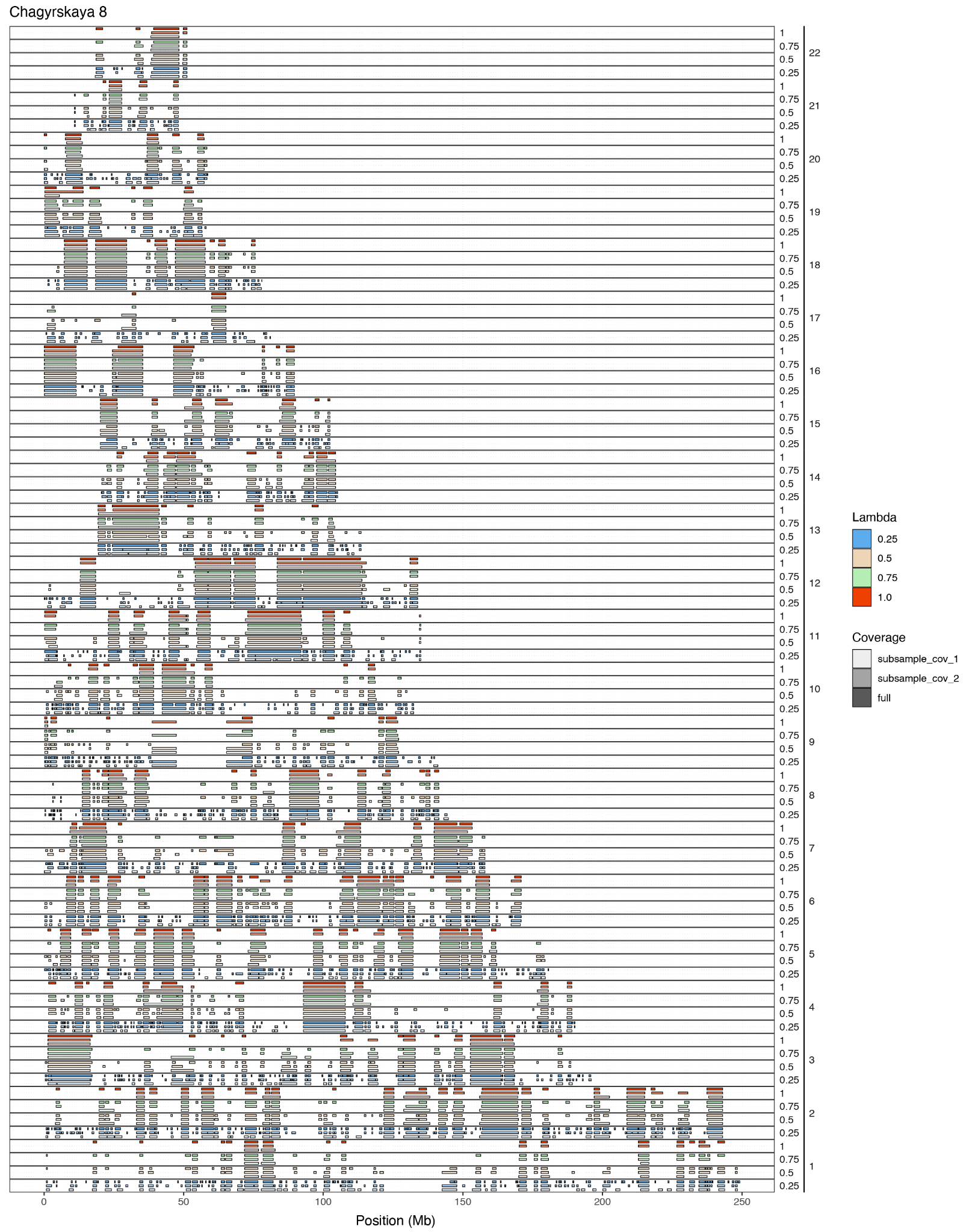

**Fig. S21 Example ROH calls in the Chagyrskaya 8 genome.** Colored segments indicate ROHs called for each chromosome, using different values of regularization parameter λ. Full and subsampled datasets are indicated using segment color transparency.

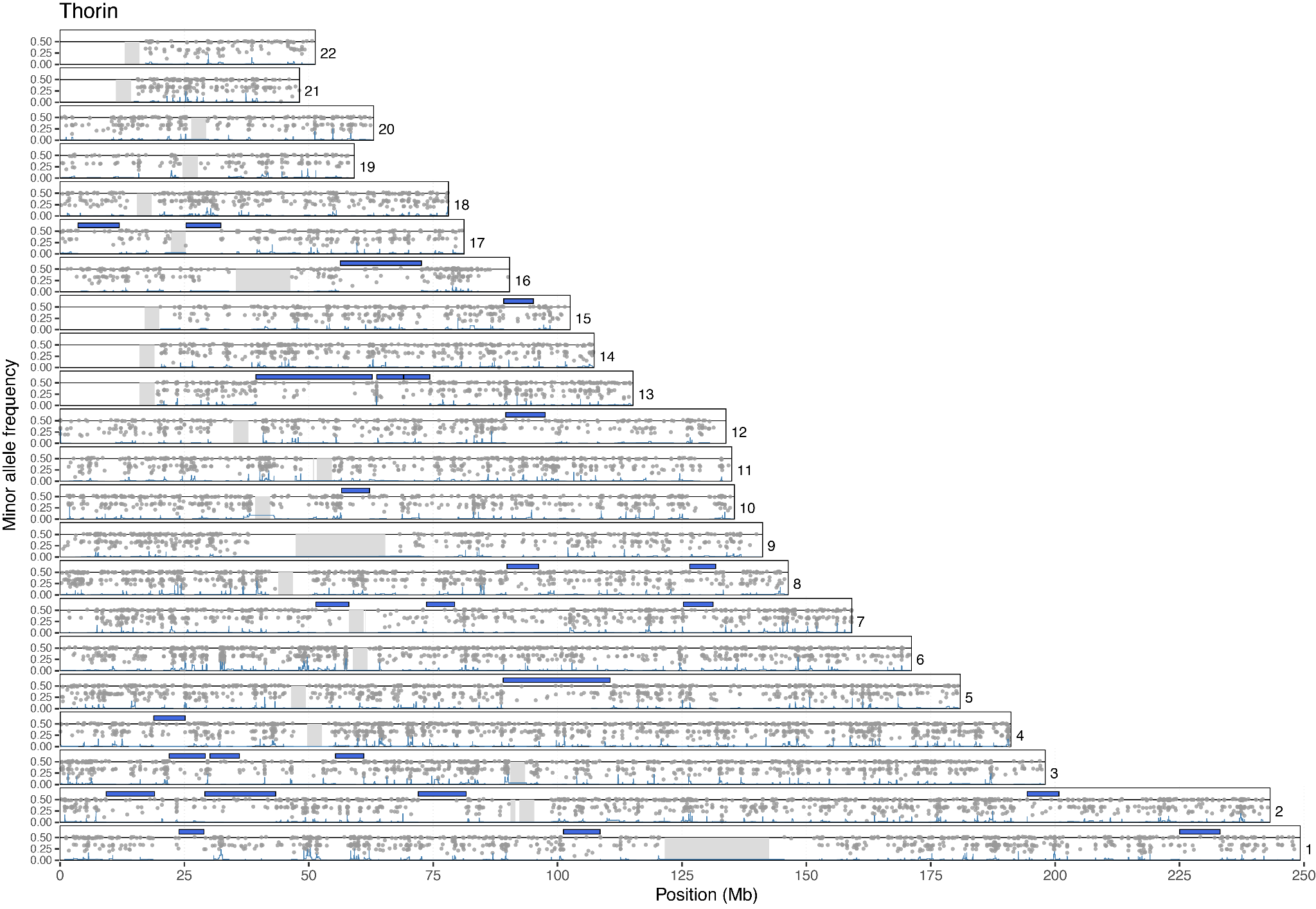

**Fig. S22. ROH calls in the Thorin genome.** Plot shows the distribution of read minor allele frequencies (grey dots) and fused lasso regression coefficients (blue line) for the Thorin genome. Colored segments indicate ROHs with length ≥ 5Mb called for each chromosome. Centromeric regions are indicated with grey boxes. A small amount of random jitter was added to the minor allele frequency values to improve visualization and reduce overplotting.

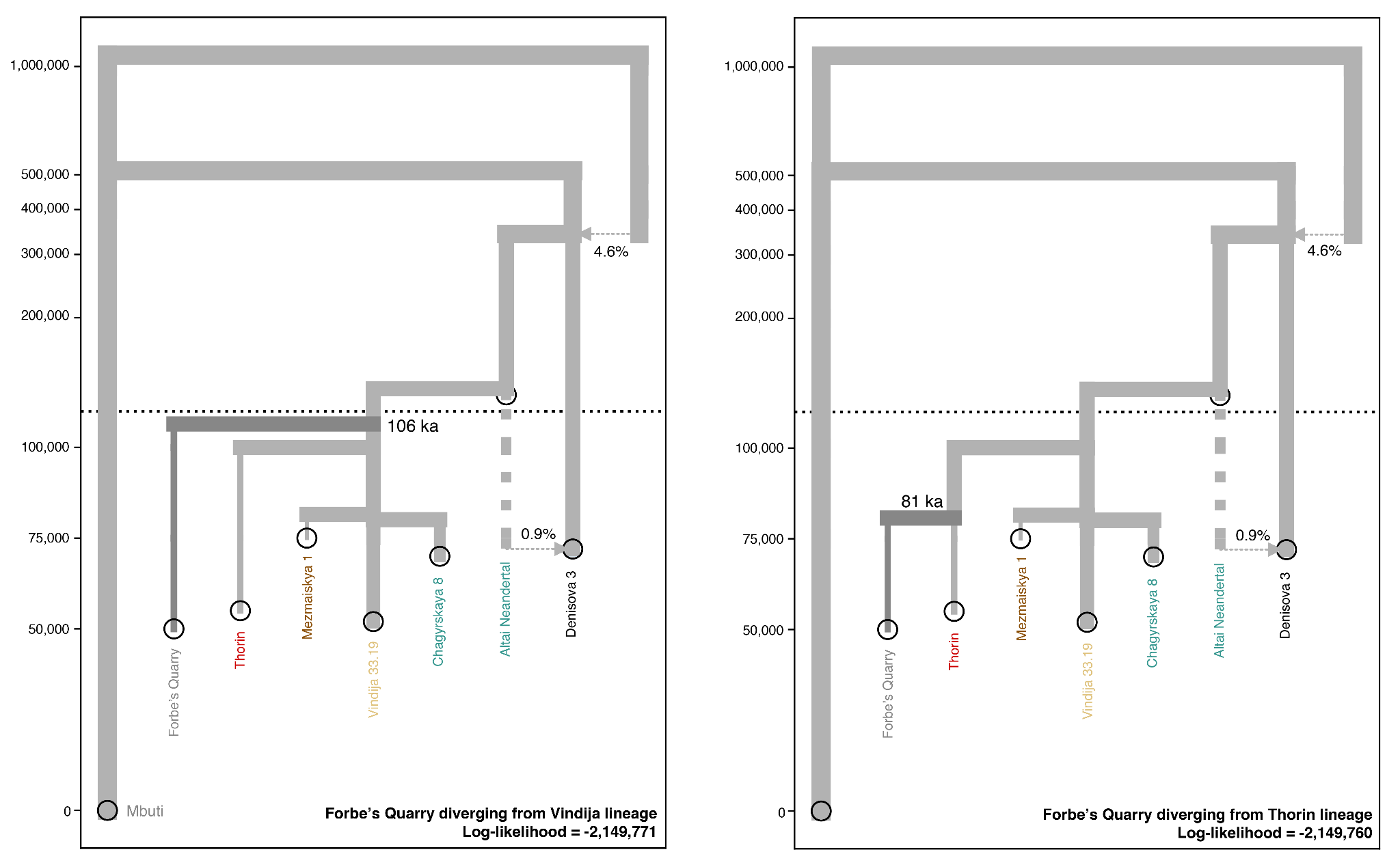

**Fig. S23. Demographic models with Forbes’ Quarry Neanderthal.** Demographic models contrasting two different scenarios of fitting the Forbes’ Quarry Neanderthal, either diverging from the Vindija-like lineage (left) or from the Thorin lineage (right). Point estimates for FQ divergence time and final Log-likelihood for the best fit are indicated for each model.

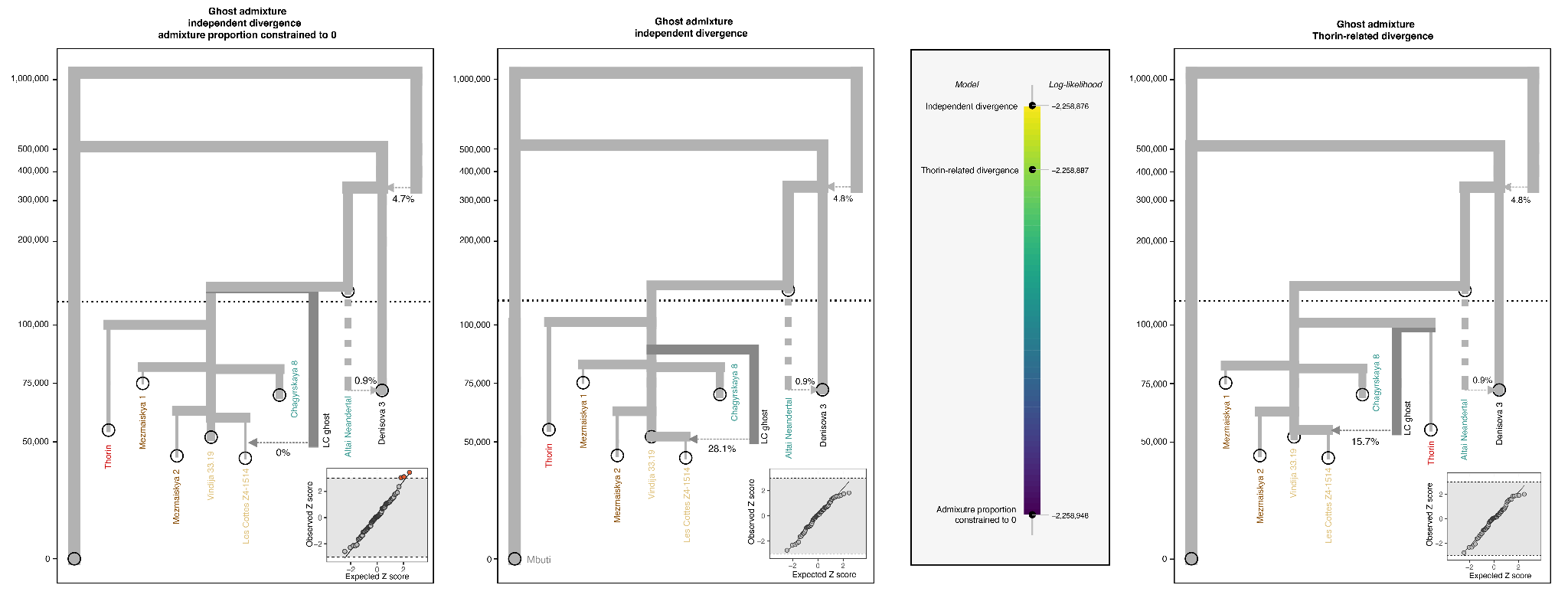

**Fig. S24. Ghost admixture into Les Cottes Z4-1514.** Demographic models contrasting different scenarios of admixture from a ghost lineage into Les Cottes Z4-1514. (left, middle) Best-fitting models with ghost admixture from a lineage (“LC ghost”, darker gray) diverging from the ancestral European Neanderthal lineage, with admixture proportion constrained to 0 (left) or free (middle). (right) Best-fitting model with ghost lineage diverging from Thorin lineage. Insets in demographic models show model fit using quantile-quantile plots of Z-scores for *f*_4_* statistics predicted from the model versus those observed in the data. Points marked in red indicate statistics with |Z|>3. Log-likelihood values for the three models are indicated in the colored bar chart (middle).
